# Mutualistic interactions between *Escherichia coli* and *Bifidobacterium bifidum* enable degradation of human milk oligosaccharides in healthy infants

**DOI:** 10.1101/2025.03.24.645027

**Authors:** David Seki, Shaul Pollak, Magdalena Kujawska, Raymond Kiu, Antia Acuna-Gonzales, Lucy Crouch, Cassie Bakshani, Peter Chivers, Monique Mommers, Nils van Best, John Penders, Lindsay J Hall

## Abstract

The development of the human gut microbiota during infancy is marked by frequent colonization of *Enterobacteriaceae*, a bacterial family notoriously associated with various diseases. Yet, despite their prominence in the absence of illness, their exact ecological role during healthy maturation of the infant gut remains poorly explored. Here, we analyse longitudinal stool samples from healthy, term-born, breastfed neonates (n=41) at two, six, and eleven months post-delivery, as well as microbiota of related mothers (n=30) with shotgun metagenomic sequencing, complemented by novel computational approaches and experimentation. Strain-resolved profiling indicates that dominant *Bifidobacterium* are frequently shared between infants and parenting mothers, while *Escherichia coli* originate from other sources, yet often persist within individuals. Despite their ecological differences, these genera co-exist, and both display evolutionary adaptations related to the utilization of human milk oligosaccharide (HMO) degradation products. We demonstrate that interactions between *E. coli* and *Bifidobacterium bifidum* are mutualistic in co-culture, where *E. coli* supplies cysteine to its auxotrophic partner, facilitating the cooperative degradation of 2′-fucosyllactose (2′FL), the predominant HMO. In turn, the liberated monosaccharides support *E. coli* proliferation and niche occupation. These findings reveal a fundamental cross-feeding interaction during development of healthy infant gut microbiota.

## Introduction

Infant health is tied to the postnatal assembly of the gut microbiota. Despite major research interest^1–4^, our understanding of the principles underlying microbial community assembly is still lacking. This is largely due to difficulties in identifying true microbe-microbe and host-microbe interactions in natural populations, whose dynamics arise from the interplay between deterministic factors such as diet^5^ and geographical location^6^ with countless other stochastic components.

The early life breastmilk diet, rich in human milk oligosaccharides (HMOs), provides prebiotic substrates to colonising microbes and facilitates reproducible iterations of host-specific microbiome assembly across generations. It is commonly believed that HMOs are preferentially utilised by commensal bacteria, prominently exemplified by *Bifidobacterium* in neonates^7^. However, the genomic potential for utilisation of HMOs appears to be widely distributed among microbiota members^8^. In addition, many ‘primary degraders’ of HMOs must express costly enzymes to break down complex HMO molecules outside of cells before they can absorb smaller, more universally nourishing sugar fragments^9^. In theory, this creates additional niches for recipients of such breakdown products (‘secondary degraders’) that do not invest in costly primary degradation of HMOs but compete for their consumption. However, the environmental relevance of HMO cross-feeding in the neonatal intestine is not fully understood.

*Bifidobacterium* are widely recognized as infant symbionts – given several indications that in return for habitation, they induce immunological tolerance^10^ and provide colonisation resistance against pathogens during early-life development^11^. But loss of such symbiotic relationships is not unusual and allows for gastrointestinal (GIT) overgrowth by pathobionts - a term used to label common inhabitants of a given environment, that may promote disease when host conditions become altered^12^. In neonates, *Enterobacteriaceae* are considered pathobionts, with *Enterobacteriaceae*-driven infectious diarrhoea being a major cause of neonatal sepsis and death^13^, especially in low- and middle-income countries (LMICs), where *Escherichia coli* is often found in high abundances^14^. Because *Enterobacteriaceae* do not effectively metabolise HMOs^15^, a critical question arises regarding the alternative resources that support their persistence in the infant gut. Moreover, despite being widespread, it remains unclear whether these pathobionts solely pose a constant threat, or if their presence may provide potential benefits to neonates under healthy conditions.

To reveal factors underlying composition and function of gut microbiota in neonates, we employed metagenomic sequencing of stool samples in a Dutch cohort of healthy neonates (n=41). Notably, these infants represent a selected subset of the larger LucKi Birth cohort^16^, chosen for their rare status of exclusive breastfeeding post-delivery before gradually transitioning to a more complex diet during the first year. This well-defined dietary trajectory provides a unique model system to investigate organising principles of infant microbiome community dynamics. To explore it in depth, we developed a new computational pipeline - MAJIC (Mean Across Jaccard Index Checkerboards), and profiled microdiversity of infant and maternal gut microbiota, revealing ecological aspects for the transmission of strains among individual hosts. Furthermore, without relying on annotation of genes, we identified ecological links between *Bifidobacterium* and *Escherichia* over the acquisition of HMO degradation products in neonates. Lastly, our experimental findings demonstrate that in co-culture, *E. coli* actively supplies cysteine to auxotrophic *B. bifidum*, facilitating the cooperative breakdown of 2’-fucosyllactose (2’FL), one of the predominant HMOs.

## Results

### Species-level composition of exclusively breast-fed infants during the initial year of life

Although the order of colonizing bacteria is well described^4,17,18^ the mechanisms driving assembly of the early-life gut microbiome remain poorly understood. Here, we employed shotgun metagenomics to sequence 78 longitudinal stool samples from 41 healthy neonates, including an early time-point (n=18) at 2 months post-delivery at which each infant was exclusively breast-fed, a time-point at 6 months post-delivery (n=25), commonly marking the onset of weaning^19^, and a late time-point at 11 months post-delivery (n=35), by which complex foods were introduced. In comparison to the broader LucKi Birth cohort^16^, where 74.1% of infants are no longer exclusively breastfed by 6 months, our cohort represents a rare subset. Additionally, to evaluate potentially relevant components of vertical transfer between mother and infant we sequenced 30 stool samples of respective mothers at 2 weeks post-delivery (Figure 1A and Table 1).

**Figure 1:**
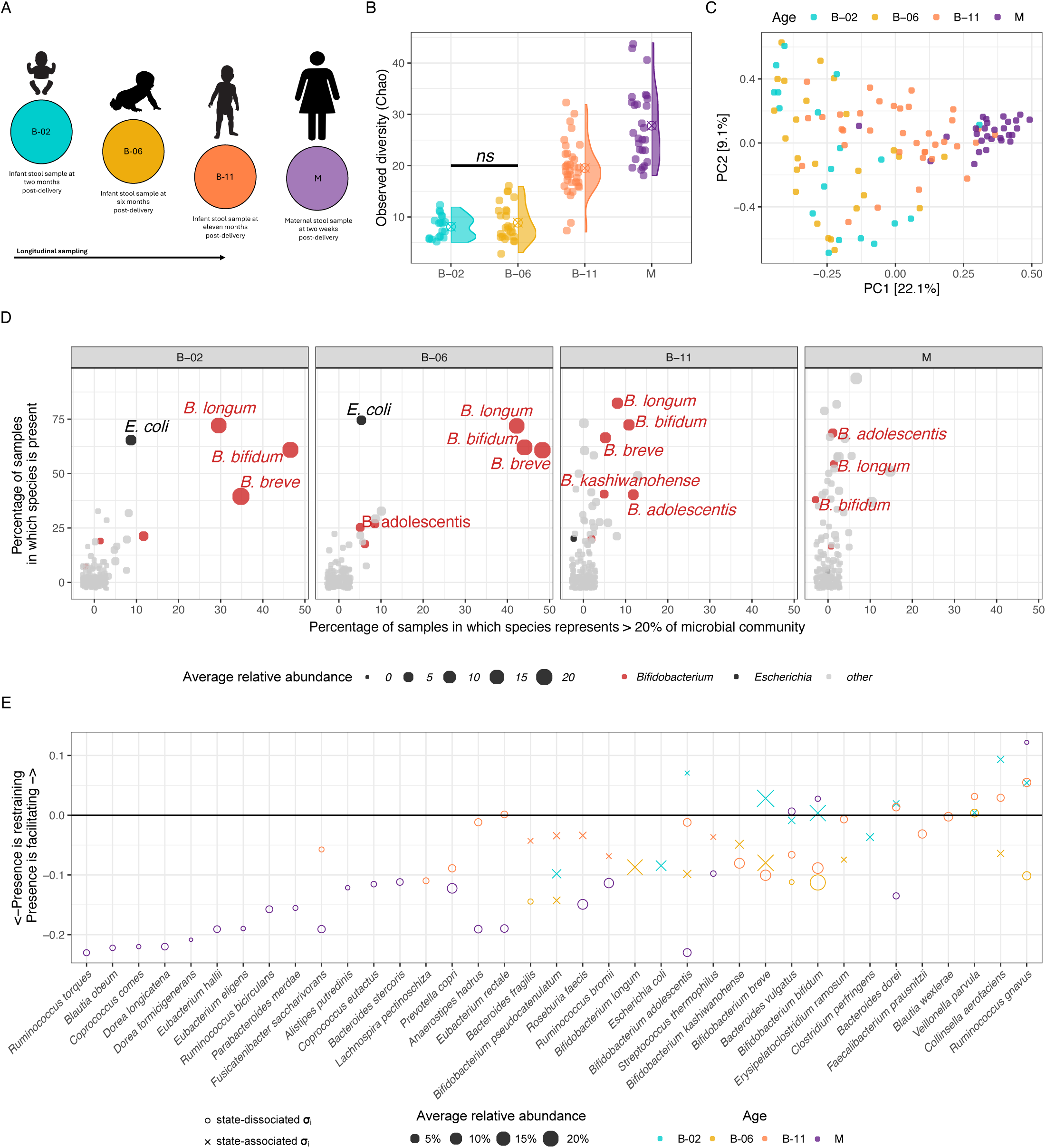
Characteristics of infant gut microbiota during the initial year post-delivery at species-level resolution. A) Overview of study-design. Samples were obtained from babies at two-, six-, and eleven months post-delivery (B-02, B-06, B-11), and one maternal sample was obtained two weeks post-delivery (M-02) F) Principal component analysis (PCA) of relative species counts. B) Microbial richness measured by count of unique bacterial species per age-group (Chao). C) Principal component analysis (PCA) of relative species counts over Hellinger distance. D) Scatterplot highlighting prevalence (percentage of samples in which species are present) and dominance (percentage of samples in which species represent >20% of the microbial community) for *Bifidobacteria* and *Escherichia coli* in maternal and infant samples. E) Differences in mean Jaccard dissimilarity Δµ_i_ = µ_i_^+^ - µ_i_^-^, wherein negative values indicate that microbiota are more restrictive when focal species is missing [𝞼_i_^-^ (µ_i_^-^)], and conversely more facilitating when Δµ_i_ > 0, therefore focal species being present [𝞼_i_^+^ (µ_i_^+^)]. It was identified via comparisons to shuffled data whether focal species (𝞼_i_) is state-associated (X) or state-dissociated (O), and colour indicates age-groupings.

**Table 1:** Subject cohort demographics.

First, we excluded species which occurred in <10% of samples with a minimum relative abundance of 1% per respective age-group. Resulting matrices served all subsequent analyses. Overall, the number of observed species was similar at 2 and 6 months post-delivery (Chao-1, t.test, p.adj = 0.23) and increased significantly thereafter (Chao-1, t.test, p.adj < 0.001, respectively; Figure 1B). Principal component analysis (PCA) indicated that infant gut microbiotas were most dissimilar to each other at two months post-delivery, gradually becoming more similar to same-aged infants and to the maternal microbiota over time (Figure 1C). This trend that was more pronounced between infants and their own mothers compared to unrelated mothers (Figure S1A). Microbiota composition varied significantly with age (PERMANOVA, p.adj < 0.05), except between two and six months post-delivery (PERMANOVA, p.adj = 0.731), highlighting the selective pressure exerted by a breastmilk-based diet. *Bifidobacterium breve* was the most abundant species in infants, followed by *Bifidobacterium bifidum,* and *Bifidobacterium longum* subspecies *longum.* As long as breastmilk was supplied, *E. coli* occurred at low counts, but consistently prevalent (66.67% and 72% prevalence at two and six months post-delivery), alongside initially dominant *Bifidobacterium* (Figure 1D, Figure S1B, and Table 2).

**Table 2:** Abundance distribution of microbiota.

In ecology, checkerboard units (C-scores) measure the extent to which pairs of species avoid co-occurring in similar habitats, with higher scores indicating stronger competition^20^. To assess the tendency of microbial species-species exclusion in the infant gut during early-life, we calculated the average number of checkerboard units for each unique species pair. C-score distributions differed from respective randomised distributions within (ks.test, p < 0.0001) and decreased with age (t-test, respective p.adj < 0.001), except between two and six months post-delivery (t-test, p.adj = 0.62) (Figure S1C). However, species-resolved co-occurrence analysis revealed a limited number of significant correlations between species before weaning, including, for example, a negative relationship between *B. longum* subsp. *longum* and *B. pseudocatenulatum* at two months post-delivery, but no other significant putative interactions between core GIT microbiota (Figure S1D). This suggests that a milk-based diet and other host factors initially drive microbial selection, creating an environment where *E. coli* co-exists with dominant *Bifidobacterium,* despite lacking the genetic capacity for HMO degradation^15^. As weaning progresses, the selective pressure from milk diminishes, leading to an increase in both negative and positive species-species interactions, ultimately reducing the abundance of early-dominant species.

### Infant gut microbiota follows deterministic succession amidst stochastic dynamics

While host-driven factors such as immune development and a milk-based diet play a key role in shaping infant microbiomes, the extent to which microbiome-intrinsic factors influence primary microbial succession in the gut remains poorly understood. To address this, we developed a new computational pipeline - MAJIC (Mean Across Jaccard Index Checkerboards) to quantify the influence of focal species on community composition. Our approach involves comparing microbial communities in which focal species (i) is present (𝞼_i_^+^) to those where it is absent (𝞼_i_^-^). To ensure robust statistical testing, we generate 10,000 random 𝞼_i_^+^ and 𝞼_i_^-^ datasets by shuffling the presence/absence labels of focal species (i) each time. This randomization scheme accounts for differences in sample sizes and abundance distributions of individual species. For each focal species, we calculate the mean Jaccard dissimilarity of samples in 𝞼_i_^+^ (µ_i_^+^), and in 𝞼_i_^-^ (µ_i_^-^), as well as the difference in mean Jaccard dissimilarity (Δµ_i_ = µ_i_^+^ - µ_i_^-^), comparing these values to the random distributions generated from shuffled data. A negative Δµ_i_ (< 0) indicates that microbiota are more constrained in the presence of the focal species (since µ_i_^+^ < µ_i_^-^), whereas a positive Δµ_i_ (> 0) indicates greater community variability when the focal species is present (Figure S2A).

At two months post-delivery, infant microbiotas exhibited considerable flexibility. Dominant species (e.g. *B. breve*, *B. bifidum, B. longum subsp. longum*), showed no significant differences between µ_i_^+^ values and those obtained from shuffled inputs (ks.test, p > 0.05, respectively), suggesting no strong association with specific community states. These species were therefore classified as ‘state-dissociated’. By six months post-delivery, µ_i_^+^ distributions remained similar, but the number of ‘state-associated’ species increased. At eleven months post-delivery and in maternal samples, overall microbiota flexibility declines, accompanied with more in ‘state-associated’ species (Figure S2B). Analysis of Δµ values revealed that maternal microbiotas were more constrained in the absence of *Ruminococcus gnavus*, suggesting that this putative keystone species plays a role in niche expansion when present. Similarly, infant microbiomes lacking *R. gnavus* or *Collinsella aerofaciens* were more restricted at two and eleven months post-delivery. However, at six months, the absence of either species had a facilitating effect, suggesting that their niche-defining roles are less relevant during the weaning transition. *Veillonella parvula* also displayed a slight facilitating effect across all infant time-points, while *Bacteroides dorei* showed a similar trend at two and eleven months post-delivery (Figure 1E). These findings suggest that despite their recognised importance for infant development, dominant *Bifidobacterium* species do not function as putative keystone species in the classical ecological sense^21^. Instead, their loss is likely compensated by functional equivalent species, maintaining overall composition and functional stability despite individual species’ absence.

Our analysis extends beyond presence/absence patterns to identify pairs of state-associated microbes that quantitatively influence each other’s abundance within the community. For each focal species (i) as defined above, we identified any species (j) whose abundance patterns significantly differed between matrices containing species (i) (𝞼_i_^+^) and those without it (i) (𝞼_i_^-^) using a Wilcoxon rank-sum test. At two months post-delivery, most significant associations occurred between ‘state-dissociated’ species, suggesting they were likely stochastic in nature. However, the overall number of significant associations increased by eleven months post-delivery, while maternal samples exhibited a reduced number of such interactions. Interestingly, all significant associations between ‘state-associated’ species were exclusively negative (Figure S2C), indicating potential competitive interactions that become more pronounced at the microbiota matures.

### Strain-level microdiversity of exclusively breast-fed infants during the initial year of life

To gain a better ecological understanding of *Bifidobacteria* and *Escherichia* within complex infant gut microbiotas, strain-level information is essential. Therefore, we conducted we conducted de-novo assembly of metagenome assembled genomes (MAGs) from metagenomic reads, resulting in 458 unique dereplicated genomes (Table 4). On average, MAGs accounted for 75 ± 14% of metagenomic reads in infants at two months, 78 ± 5% at six months, and 75±7% at eleven months post-delivery, while covering 68±7% of maternal metagenomes. All reads were subsequently mapped to this set of dereplicated genomes, and inStrain^22^ was used to profile the nucleotide diversity (π), assessing both within (intra-π) and across (inter-π) individual variation.

Intra-π for all MAGs were not significantly correlated with their coverage within samples (Spearman, p.adj > 0.05, respectively, Figure S3A). However, when aggregated at the genus-level, *Bifidobacterium*-intra-π showed a negative correlation with combined bifidobacterial coverage (Spearman, cor −0.190, p.adj = 0.0499), suggesting that higher bifidobacterial strain-diversity may be associated with a decrease in bifidobacterial abundance. Overall, intra-π remained stable between infants at two and six months post-delivery, but increased significantly thereafter (t.test, p.adj < 0.05 for every other pairwise comparison). Notably, *E. coli* MAGs consistently displayed elevated intra-π, suggesting presence of multiple strains co-existing within individuals, whereas bifidobacterial MAGs rarely showed such patterns (Figure 2A).

**Figure 2:**
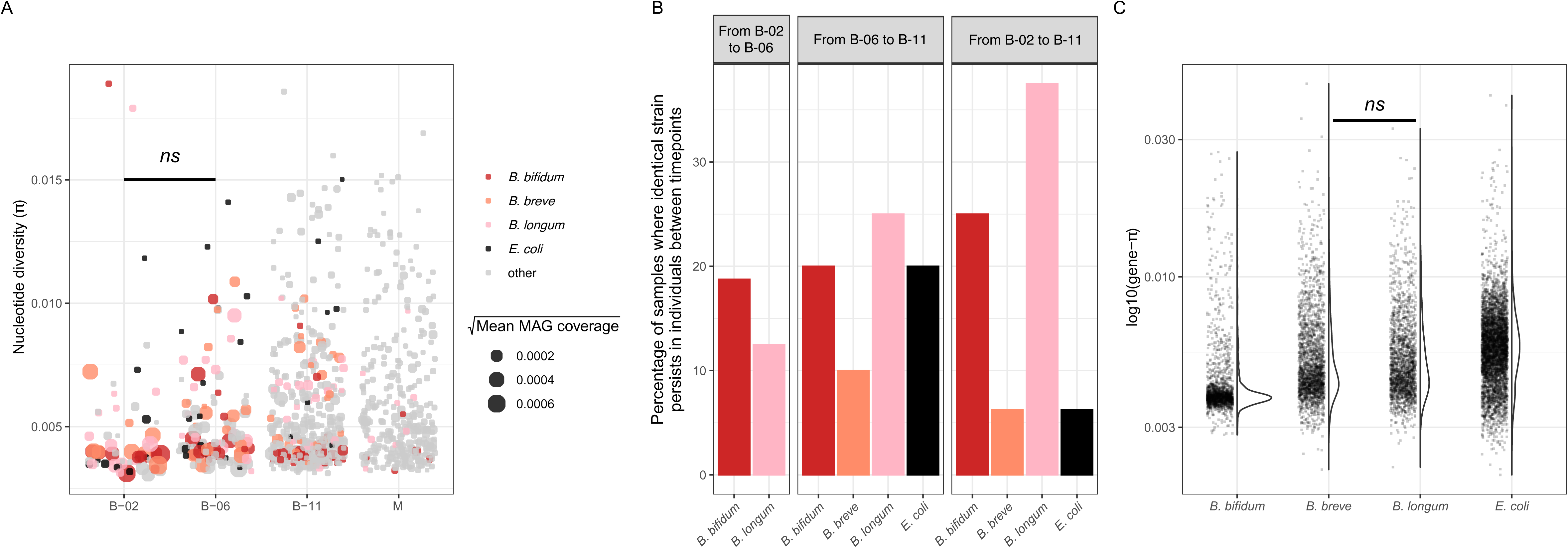
Microdiversity and strain-sharing among infant and maternal gut microbiota. A) Dot-plot visualizing the distribution of overall nucleotide diversity (π) according to age-groups. Each dot represents a metagenome-assembled-genome (MAG) dereplicated at 95% average nucleotide identity (ANI), and size of dots represent mean coverage MAGs. MAGs of *B. bifidum, B. breve, B. longum*, and *E. coli* are coloured in red, salmon, pink, and black. B) Count of strain-persistence events where identical strains of *B. bifidum, B. longum, B. breve,* and *E. coli* re-occurred between adjacent timepoints within the same individual. C) Violin / Dot-plot visualizing the distribution of overall gene nucleotide diversity (gene-π) in *B. bifidum, B. breve, B. longum*, and *E. coli*. Each dot represents a gene in respective MAGs.

To investigate strain-level transmission patterns, we next applied a threshold of pop-ANI 99.5% to distinguishing between identical strains and strain-variants, and quantified strain-sharing events between individuals. Identical *B. adolescentis* strains were shared among 22% of maternal microbiota, while *B. longum subsp. longum* strains were shared among 31% of infants at eleven months post-delivery (Figure S3B). Furthermore, pop-ANI values were significantly higher between parenting mothers and their infants, as compared to random mother-infant pairs at two and eleven months post-delivery (t-test, p.adj < 0.0001 and 0.008, respectively; Figure S3C). Notably, B. *longum* subsp. *longum* strains were frequently shared between infants and their mothers, whereas no such co-occurrence was observed for *E. coli* strains in mother-infant pairs (Figure S3D). Next, we assessed strain persistence within infants across time for the four most abundant and prevalent species in the infant gut microbiome (*B. bifidum, B. longum subsp. longum, B. breve*, and *E. coli*). Between two and six months post-delivery, identical *B. bifidum* and *B. longum subsp. longum* strains persisted within individuals in 12.5% of cases, while no such persistence was observed for B. *bre*ve. From six to eleven months, identical *E. coli* strains persisted in 10% of cases, and from two to eleven months, *B. longum subsp. longum* strains persisted in 25% of infants (Figure 2B). These findings indicate that *B. adolescentis* is frequently shared among maternal microbiota, while *B. longum subsp*. *longum* strains are commonly transmitted between infants and parenting mothers early post-delivery. In contrast, bifidobacterial strains showed less consistent co-occurrence between maternal and infant samples, suggesting alternative colonization routes and persistence mechanisms in infants. *E. coli* displayed high microdiversity overall, and although not shared between mothers and infants, strains persisted within infants across the first year post-delivery, suggesting extrinsic sources as the origin of colonization.

We hypothesized that the observed differences in host-associated transmission patterns would be reflected in the genetic nucleotide diversity (gene-π) within dominant *Bifidobacterium* and *Escherichia* species. Specifically, given that *E. coli* exhibited limited mother-to-infant transmission as well as higher pop-ANI values, we expected weaker host-specific selective pressures, leading to higher nucleotide diversity across a broader range of genes. Conversely, for *Bifidobacterium*, where frequent strain transmission and persistence between hosts occur, we anticipated stronger purifying selection on genes essential for host adaptation. To identify these genes, we applied EggNOG-mapper v6 for functional annotation^23^, and profiled gene-π in dominant *Bifidobacterium* and *Escherichia* species. As expected, *E. coli* displayed the highest gene-π on average. Additionally, gene-π did not significantly differ between *B. breve*, *B. longum subsp. longum* (t.test, p.adj > 0.05), suggesting that despite differences in transmission dynamics, their ecological roles and adaptive strategies are similar. However, *B. bifidum* exhibited a particularly narrow gene-π distribution that significantly differed from the remaining species (t.test, p.adj < 0.0001, Figure 2C), indicating stronger genetic constraints on its adaption. Next, we categorised genes with gene-π values above the 99^th^ percentile as ‘loose’, and those below the 1^st^ percentile as ‘restricted’. Notably, ‘loose’ genes included a starch-binding outer membrane protein from the SusD/RagB family in *B. bifidum,* - potentially involved in HMO transport. Conversely, ‘restricted’ genes, aside from several housekeeping genes, included a putative chitinase in *B. bifidum* and a multiple sugar transporter system in *B. breve*, both likely involved in HMO acquisition (Table 3). These findings highlight conserved functions essential for strain-specific adaptations to the host environment.

**Table 3:** Most restricted or loose genes in MAGs of dominant *Bifidobacteria* and *E. coli*.

### Coevolving ecological strategies for the consumption of HMOs feature dependencies between *Bifidobacterium* and *E. coli*

HMOs serve as the primary carbon source for colonizing infant gut microbes, and therefore central ecological drivers of primary succession^24^. To overcome the lack of traditional gene annotation and gain a more holistic understanding of microbial traits essential for acquisition of HMOs in complex communities, we employed a recently developed annotation-agnostic approach to infer microbial trophic strategies from genomic data^25^. We hypothesized that glycoside hydrolases (GHs) are necessary for HMO degradation, costly to produce, and benefit not only the GH producers, but also all potential consumers of the resulting degradation products. As such, GHs function as model public goods, where their utility extends beyond enzyme synthesis to encompass a suite of associated traits, largely unknown but likely including chemotaxis, biofilm formation, and membrane transport, that enhance the efficient capture and utilization of GH-derived breakdown products. Public goods like GHs are subject to rapid evolutionary shifts between parasitism and cooperation due to trade-offs between production costs and competitive advantages. Furthermore, their evolutionary trajectories are shaped by high rates of gene deletion and horizontal transfer. By leveraging these dynamics, we can trace patterns of co-evolution between the GHs (public goods) and battery-traits across microbial species with distinct ecological strategies, providing deeper insights into how microbial communities adapt to HMO-rich environments^25^.

Because of their central role in HMO degradation, we sought to identify protein families that have co-evolved with GH2 family enzymes across our set of 486 unique dereplicated genomes. In total we identified 2803 such protein families (𝜺_GH2_), with an average of and 302 ± 284 per genome (𝜺^i^_GH2_). To further investigate the degree of polymer degradation specialization in different microbial lineages, and the associated strategies, we compared the number of GH2 enzymes encoded in a genome (O_GH2_) to two derived quantities based on the co-evolutionary patterns of GH2-related genes encoded within single genomes. The first, E_GH2_ – was estimated using an elastic-net linear regression model fitted across all genomes. This value represents the expected number of GH2 enzymes a genome ‘should’ encode based on its genomic background of 𝜺^i^_GH2_. If true coevolution were occurring, E_GH2_ and O_GH2_ would be strongly correlated, as recently demonstrated for chitinases and their coevolved genes^25^. Indeed, we observed a strong correlation between E_GH2_ and O_GH2_, supporting the validity of our approach and the genuine co-evolution of 𝜺_GH2_ across diverse microbial phyla (Spearman, R = 0.83, p < 0.00001). However, our data also suggest that the specialized organisms for GH2 breakdown product utilization (high E_GH2_ and high O_GH2_) primarily include species from the genera *Bacteroides* and *Parabacteroides*, rather than typically dominant members of the infant gut microbiome (Table 4).

**Table 4:** Number of observed and predicted GH-2 domain-containing enzymes per MAG.

The final parameter we examined, f^i^_CAZy_, represents the fraction of genes coevolving with GH2, that are themselves CAZymes. If the selection of coevolved genes were random, the correlation between O_GH2_ and f_CAZy_ would be expected to follow a diagonal line of identity. However, our analysis revealed a notable deviation, with *Bifidobacterium* forming a distinct cluster, alongside species from *Escherichia*, *Enterococcus*, *Clostridium*, *Erysipelatoclostridium*, *Klebsiella*, and *Megamonas*. This clustering suggests a non-random coevolutionary dynamic, likely driven by shared selective pressures that require an expanded repertoire of CAZymes for efficient colonization in HMO-rich environments. These findings highlight putative functional associations between these species (Figure 3A). Notably, *B. bifidum* deviated from this pattern, as its GH2-associated genes were exclusively non-CAZymes, suggesting an alternative metabolic adaption.

**Figure 3:**
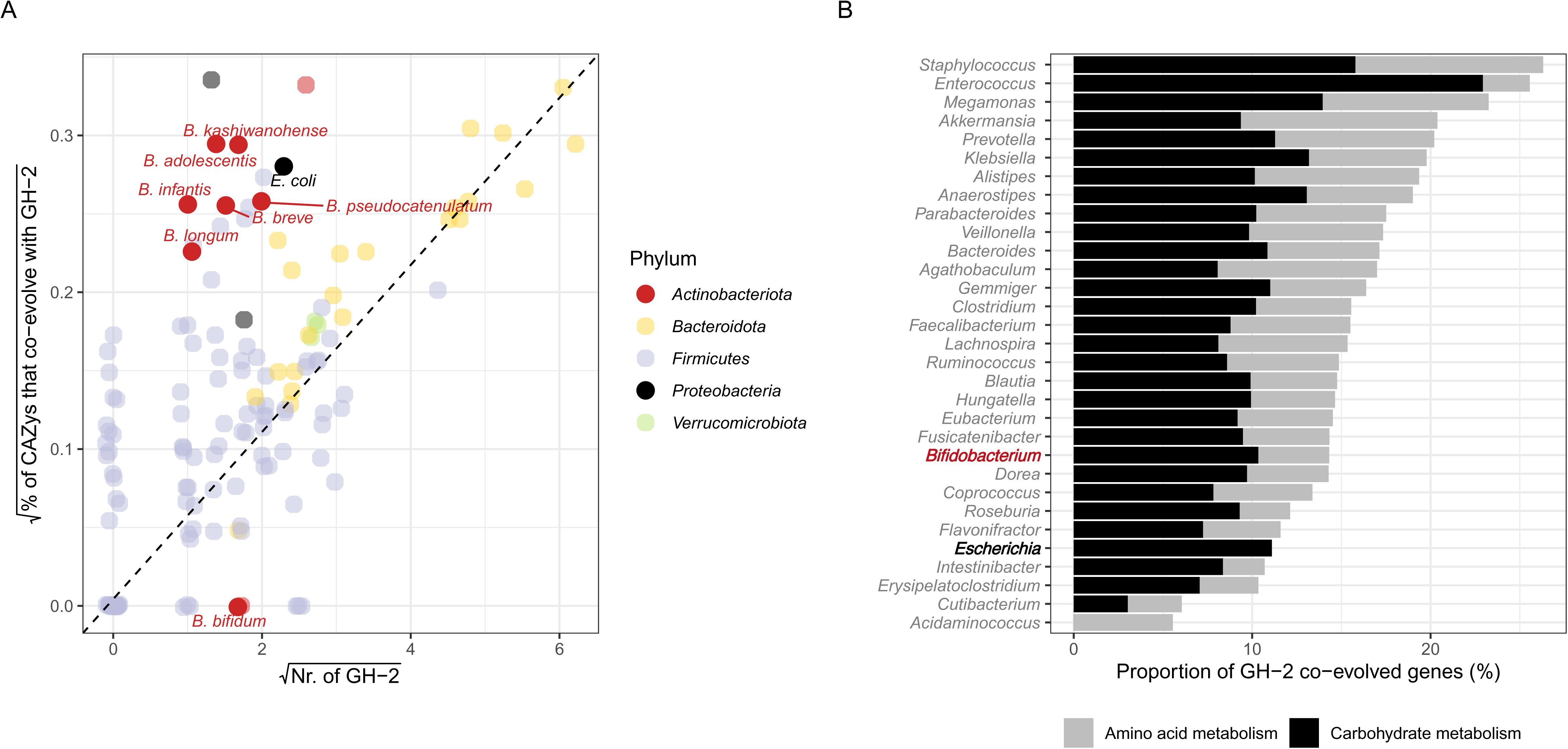
Coevolution of microbiota with GH2 domain-containing enzymes. A) Scattering of observed number of GH2 domain-containing enzymes against fraction of coevolved genes that are other carbohydrate active enzymes as well. Each dot represents a MAG, coloured by corresponding phylum. B) Bar-charts summarizing coevolved genes that correspond to amino acid or carbohydrate metabolism, which we could map to kegg-orthology (ko) groups via EggNOG mapper v6. Distribution of ko-groups among listed microbial genera significantly differed from shuffled random distributions. *Bifidobacteria* and *Escherichia* are highlighted in red and black.

To further explore the functional significance of these coevolved genes, we annotated them using EggNOG-mapper v6^23^. Notably, while all genera displayed GH2 coevolving genes that could be attributed to amino acid metabolism, *E. coli* uniquely lacked such associations (Figure 3B and Figure S4). This suggests that while *E. coli* mirrors the dynamics of *Bifidobacterium*, which are specialized in HMOs degradation, it adopts the ecological role of a versatile prototroph. Additionally, it may be less reliant on simultaneous amino acid degradation to access GH2 breakdown products, despite possessing the functional capacity to do so^26^.

### *B. bifidum* and *E. coli* cooperate over the degradation of 2′-O-fucosyl-lactose

We hypothesized that *Bifidobacterium* and *E. coli* leverage complementary metabolic roles in extracellular HMO degradation, where *Bifidobacterium* spp. act as primary degraders, breaking down HMOs into monosaccharides and smaller oligosaccharides that *E. coli* can utilize as carbon sources for growth.

To test this hypothesis, we focused on the metabolism of 2′-O-fucosyl-lactose (2’FL), the most abundant breastmilk HMO^27^. First, we isolated representatives of the three most abundant *Bifidobacteriaceae* from infant stool – *B. longum* subsp. *longum, B. breve,* and *B. bifidum*, and assessed their growth on 0.1% 2’FL (w/v) as the primary carbon source in modified M9-medium (m-M9)^28^. In pure cultures, only *B. bifidum* was able to grow at 37°C under anoxic conditions (Figure 4A), and none of the isolates grew in m-M9 without 2’FL (Figure S5A). This confirmed that the additives to m-M9 were insufficient to support growth without an external carbon source. Importantly, *B. bifidum* failed to grow in m-M9 + 0.1% 2’FL when casamino acids or cysteine were omitted, indicating this isolate is a cysteine auxotroph with additional metabolic dependencies (Figure S5B)

**Figure 4:**
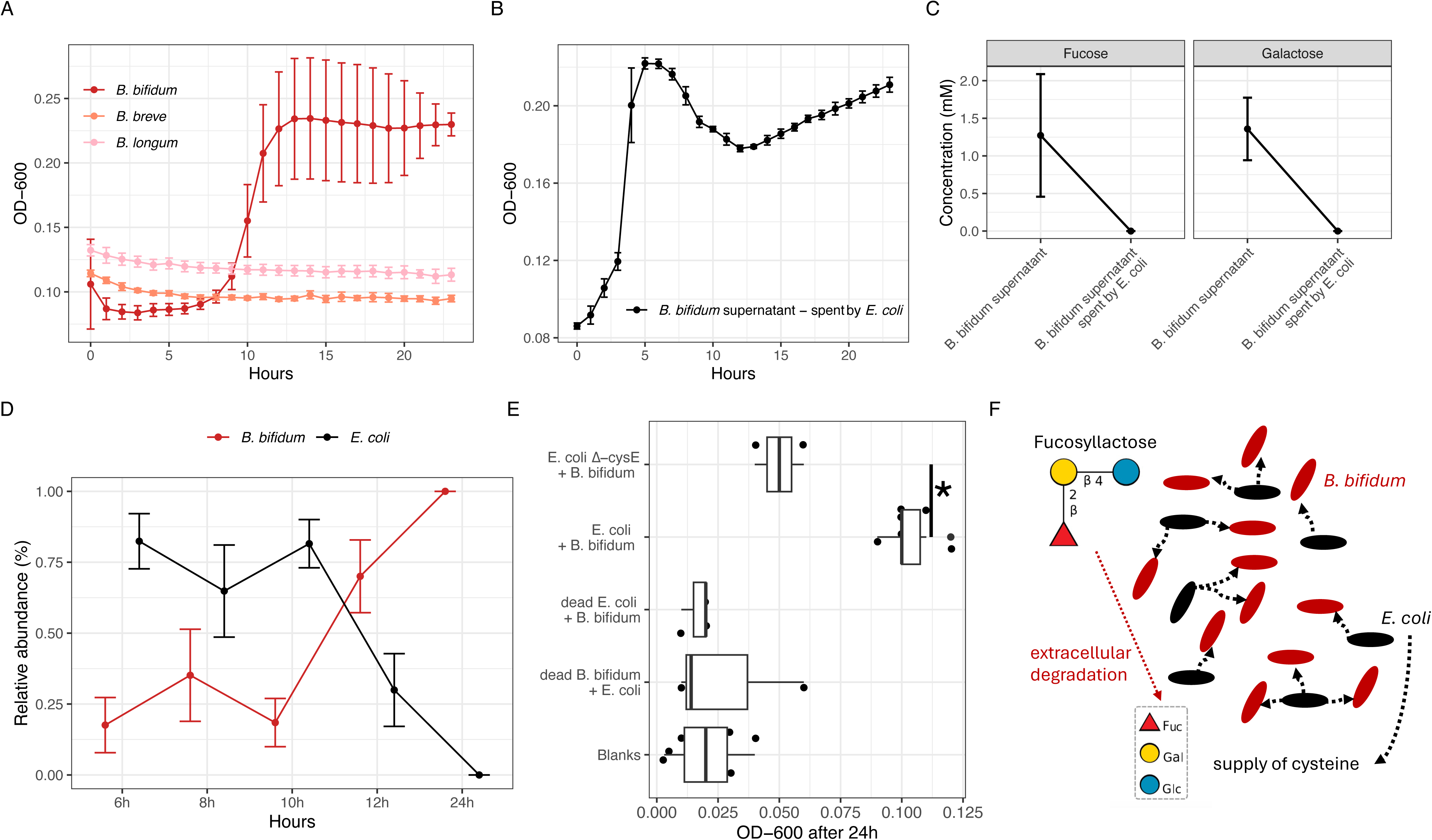
Cooperative degradation of 2’-O-fucosyl-lactose between *E. coli* and *B. bifidum*. A) Growth of *B. bifidum* (red), *B. breve* (salmon), *B. longum* (pink) in modified M9-medium (m-M9) with 2’-O-fucosyl-lactose (2’FL) as the only source of carbon, measured by optical density (OD-600). B) Growth of *E. coli* in supernatant *of B. bifidum* previously grown in m-M9 with 2’FL as the only carbon-source. C) Concentration of fucose and galactose in m-M9 + 2’FL *B. bifidum* supernatants spent by *E. coli* at pH 6,3. D) Relative abundances of B. bifidum and E. coli in co-culture in m-M9 + 2’FL without any additions of cysteine or other amino acids. Relative abundances were estimated by fluorescent gram-stain and manual count. E) Growth of *B. bifidum* and *E. coli* co-cultures with respective controls, including a cysteine mutant (E. coli Δ-cysE) which is incapable of cysteine production. Growth was assessed by optical density measurements (OD-600) after 24 hours. Blanks included sterile m-M9 + 2’FL medium without cysteine or other amino acids. F) Visual conclusion of the article. Asterisks represent adjusted p-values: ∗p < 0.05. Dots in line-plots show group median, error bars show standard deviation.

Next, we analysed whether *E. coli* K-12 could directly metabolize 2’FL or its individual monosaccharides - fucose, galactose, and glucose as carbon sources. While *E. coli* K-12 could not metabolize 2’FL, it exhibited slight growth in m-M9 only, in the absence of 2’FL, suggesting either a preference for amino acid metabolism in the absence of polysaccharides or possibly inhibitory effects of 2’FL on growth (Figure S5C). However, *E. coli* K-12 efficiently grew when provided with all three monomers, including fucose (Figure S5D). Next, we tested whether *E. coli* K-12 could grow in the spent medium from *B. bifidum* cultures, observing growth only when the pH was adjusted to its initial value of 6.3 (Figure 4B). Repeating the experiments with higher 2’FL concentrations (0.5% w/v) confirmed that *E. coli* could utilize degradation products released by *B. bifidum*. (Figure S5E). Using thin layer chromatography (TLC), and high-performance anion exchange chromatography (HPAEC), we determined that *B. bifidum* efficiently degraded 2’FL, but left galactose and fucose unconsumed, which were subsequently utilized by *E. coli* K12 (Figure 4C and Figure S6A-D).

We hypothesized that under nutrient-limiting conditions, prototrophic *E. coli* K12 could provide essential amino acids to an auxotrophic partner, thus facilitating coexistence. To test this, we co-cultured *E. coli* K12 and our cysteine auxotroph *B. bifidum* isolate in m-M9 + 0.5% 2’FL(v/v) without added cysteine and casamino acids. After 24 hours, we observed robust co-culture growth (Figure S5F). Fluorescent Gram-staining confirmed the stable co-existence of both species throughout the experiment, with *B. bifidum* becoming dominant after 12 hours (Figure 4D and supplementary file 1). Furthermore, no significant growth was observed when one partner was ethanol-killed before inoculation, confirming the necessity of active metabolic interaction. Additionally, when we repeated the co-culture using an *E. coli* K12 mutant (Δ-cysE) lacking cysteine biosynthesis, growth was significantly impaired (Figure 4E). We therefore conclude that prototrophic *E. coli* K12 facilitate the growth and metabolic activity of auxotrophic *B. bifidum* under nutrient-limiting conditions by alleviating its cysteine dependency. In turn, *B. bifidum* enables the degradation of 2’FL, providing *E. coli* with essential carbon sources, thereby establishing a mutualistic relationship between these two species.

## Discussion

*Bifidobacterium* and *E. coli* are both core members of neonatal gut microbiota^29–31^. Here, we investigate their co-existence in exclusively breastfed, term-born infants, revealing mutualistic interactions that enhance human milk oligosaccharide (HMO) degradation. Strain-sharing analysis highlights fundamental ecological differences: obligate anaerobic *Bifidobacterium* rely on vertical and horizontal transmission between individuals, whereas facultative anaerobic *E. coli* colonizes from diverse external sources. Despite these distinct origins, both species persist remarkably once established. Evolutionary-guided genome analysis suggests that their co-existence is driven by shared selective pressures, fostering cooperation in HMO degradation. Supporting this, we demonstrate that *E. coli* supplies cysteine to auxotrophic *B. bifidum*, facilitating the joint degradation of 2’FL and promoting mutual benefits in co-culture (Figure 4F).

Successful colonization of an environment requires microbial species to overcome multiple constraints, including competition for nutrients, spatial niche occupation, and host selection. However, cooperation within microbial social networks is equally crucial, and the neonatal gut provides a prime example. Here, given the vast structural heterogeneity of HMOs^32^, it seems unlikely that a single species could specialize in the degradation of all oligosaccharides. Instead, cooperation in HMO metabolism becomes a central selective barrier of colonization. To overcome this, *Bifidobacterium* balance diverse strategies for efficient acquisition and degradation, involving both aspects of cooperation and competition. For instance, *B. bifidum*, an extracellular degrader, breaks down 2′FL in co-culture with *B. breve*, but refrains from utilizing the released fucose, while both species compete for the liberated lactose^33^. However, why does distantly related *E. coli* occur so often in these social networks, given its inability to degrade HMOs?

Fundamentally, HMO degradation can be viewed as a snowdrift game, where the stability of cooperation depends on the balance between individual advantage and collective benefit. Their outcomes are likely shaped by several ecological pressures, including abundances of interacting species and key-nutrient availability, which determine the ‘cost of cooperation’^34^. Cysteine is such a key-nutrient, given the widespread auxotrophy among *Bifidobacterium*^35^. In healthy infants, cysteine is sourced from both the breast milk diet^36^ and endogenous synthesis^37^, with its secretion at mucosal surfaces supporting the colonization of microbes and maintaining redox balance^38^. However, premature infants for instance have impaired capacities for the synthesis of cysteine, which both exacerbates oxidative stress in the gastrointestinal tract^39^ and limits its availability to microbiota. We argue that the resulting deficiency would increase the ‘cost of cooperation’, driving shifts in microbial interactions, and elevating the importance of prototrophic bacteria such as *E. coli*, which may gain an ecological advantage by fulfilling the shortage. Indeed, we find extracellularly feeding *B. bifidum* and *E. coli* to dominate in extremely premature infants^40^.

Beyond cysteine, other key-nutrients such as vitamins and transition metals could furthermore play essential roles in early-life microbial social networks. While *E. coli* possesses efficient iron acquisition systems^41^, *Bifidobacterium* may critically depend on its availability. Similarly, *Bifidobacterium* require B and K vitamins for growth and metabolism, which could as well be synthesized by other microbes and shared within the community^42^. We argue that while such dependencies may help explain *E. coli*’s persistence at low levels as a commensal, they could also serve as initial stepping stones that enable subsequent pathological overgrowth, causing severe disease^43^ and facilitating further spread of antimicrobial resistances^44^, an especially pressing issue in the Global South. Future research should prioritize understanding how microbes compete and cooperate over key-nutrients in their social networks, as uncovering these dynamics can inform the strategic selection of probiotics and dietary interventions to promote health in children.

## Material and Methods

### Experimental model, clinical definitions, stool sample collection

Stool samples were collected within the context of the LucKi Gut study, an on-going longitudinal study embedded within the larger LucKi Birth cohort study^16^ among newborns and their families. Pregnant women residing in the South Limburg region of the Netherlands were recruited through obstetrics and gynaecology clinics, lactation information sessions, and advertisements at pregnancy yoga classes, baby clothing stores, and on social media. Infants born prematurely (gestational age ≥32 weeks) were excluded. One maternal sample and 9 infant fecal samples were collected throughout the first 14 months of life.

Participants received faecal sampling starter kits consisting of stool collection tubes (Sarstedt, REF 80.623.022), cold transport containers (Sarstedt, REF 95.1123), safety bags, gloves, faeces collection devices (Fe-Col, REF FC2010), questionnaires, instructions and consent forms. The samples were collected at home and immediately stored at −20°C in their home freezers. Samples were thereafter transported to the family’s well-baby clinic using frozen transport container to preserve the cold chain. From there, samples were transported to the laboratory, where frozen faecal matter was aliquoted and stored at −80°C until further analyses. At each faecal sampling time-point, parents also completed a questionnaire gathering information on the infant’s lifestyle, health, development, medication use, and feeding practices, as well as maternal health (during pregnancy), diet, and medication use. Samples collected at 2 weeks (maternal sample), and 2, 6, and 11 months (infant samples) post-partum were used for the purpose of the present study. Written informed consent was obtained from both parents/legal caregivers prior to enrolment in the study. This research confirmed to the principles of the Helsinki Declaration. Ethical approval was obtained by the Medical Ethical Committee of Maastricht University Medical Center (study number: METC-15-4-237).

### Isolation of DNA, metagenomic sequencing, and read-based processing

DNA was extracted using the Power Soil Pro Kit (Qiagen) following the manufacturer’s protocol for stool samples, with the inclusion of one negative control per extraction-batch, consisting of nuclease free water to allow for assessment of contamination. DNA was eluted in 40 μl nuclease free water and stored at −20°C until further analysis. DNA was quantified using Qubit 2.0 dsDNA BR assay kit (ThermoFisher Scientific, Q32850). Library preparation was done following the manufacturer’s instructions using the DNA flex kit with 8 mer UDI. Shotgun metagenome sequencing (short-read) on DNA samples were performed at Azenta Life Sciences on Illumina NovaSeq platform at paired-end reads 2x 150bp. Short-read metagenomic data were processed using HUMAnN 3^45^ with default setting. Briefly, HUMAnN 3 first estimates community composition with MetaphlAn 4^46^, second it maps reads to a community pangenome with bowtie2^47^ and third it aligns unmapped reads to a protein database using DIAMOND^48^.

### De novo assembly and processing of metagenome-assembled genomes (MAGs)

Raw sequence reads were quality filtered (-q 20) with fastp v.0.20.0^49^ before de novo genome assembly using SPAdes v.3.14.1^50^ at default parameters. CheckM v.1.1.3^51^ was used for assessment of metagenome assembled genomes (MAGs). Low quality genomes (contamination > 20% and completion < 70%) excluded from further analysis. These were taxonomically classified using GTDBtk (v. 2.1.0)^52^. The MAG dataset was dereplicated with standard settings via dRep^53^, resulting in a final set of 458 unique quality MAGs (Table 4). Dereplicated MAGs were furthermore concatenated into a single fasta file, and bowtie2^47^ was used to create a mapping index from it, as well as to calculate the percentage of reads per sample that were mapping to our set of dereplicated MAGs. A ‘scaffold-to-bin’ was created using ‘parse_stb.py’ from dRep^53^, and Prodigal^54^ was used to profile all genes from the concatenated MAG file, thereby creating a ‘gene’ file. Combining these files, ‘inStrain profile’ was used to screen within-sample nucleotide diversity (π) and gene-nucleotide diversity (gene-π), and ‘inStrain compare’ was used to profile across-sample population average nucleotide identity (popANI)^22^. The ‘gene’ file was furthermore used for annotation of all genes using EggNOG mapper v. 6.0^23^.

### Detection of glycoside-hydrolase-linked genes

We expanded a previous analysis that detected genes that coevolve with chitinases^25^ by analyzing 21,559 high quality species-representative genomes from GTDB r214 (> 99% completeness, < 1% contamination). Genes were called with Pyrodigal^55^, and full-length proteins were clustered at 70% similarity using diamond deepclust followed by diamond recluster (both with --approx-id 70)^48^. Partial reading frames were then added to clusters using diamond blastp (--query-cover 100 --id 80 -b 20 -c 1 -k 10). Glycosyl hydrolases were called with dbcan4^56^. All genes were annotated using eggnog-mapper v5^57^. To look for genes that co-evolve with GH2, we only considered protein clusters that appear in >9 genomes, which reduces computational load and false positive rates due to random fluctuations^25^. Ancestral states were reconstructed using maximum parsimony, as implemented in the mpr function of the PhyloTools Julia package^58^. We detected co-evolution as correlated gain-loss events using canonical correlation analysis (CCA). We used CCA because there are many GH2 protein clusters, and CCA uncovers how the optimal linear combination of their gain/loss vectors correlates with a target vector. These correlations were compared to null distributions generated from randomized vectors, where events from different genes were randomized, therefore controlling for the mean tendency of nodes to display gain/loss events^25^. To analyze MAGs that were assembled de-novo from our own data, we first assign MAG proteins to the protein homology clusters defined above using diamond blastp (--query-cover 80 --subject-cover 80 - k 3 -c 1). Each MAG protein was assigned to the best matching protein cluster based on e-value. The number predicted number of GH2 enzymes in each MAG was calculated using the random forest regression model from above, and compared to the observed number of GH2 proteins encoded by the MAG. To identify the fraction of GH2 co-evolved genes that themselves are carbohydrate active enzymes, we compared annotated coevolving genes against the PFAM database using PFAM.db v3.8.2^59^.

### Growth assays with *Bifidobacteria* and *E. coli*

*B. bifidum*, *B. longum subsp. longum*, and *B. breve* were isolated from infant stool samples by spreading 10^-7^ dilutions of faecal slurries onto MRS agar with cysteine (0.1% w/v; Sigma-Aldrich) and mupirocin (50 mg/l; Sigma-Aldrich) (m-MRS). Plates were incubated at 37°C under anaerobic conditions for 48 hours. Per plate, several colonies were picked at random and grown in liquid m-MRS, repeating the 48 hours long incubation. From incubations with visible growth, DNA was extracted using the Power Soil Pro Kit (Qiagen) following the manufacturer’s protocol and finally eluted in 40µl nuclease-free water. DNA isolated from pure bacterial cultures was subjected to whole-genome-sequencing via Illumina NextSeq 2000. Obtained sequencing reads were pre-processed with fastp v.0.23.2^49^. Unicycler v.0.4.9^60^ with the “–mode conservative” option was used to produce assemblies, after which contigs below 1000 bp were filtered out. Genome completeness and contamination were estimated using CheckM v.1.2.0^51^, and we retained sequences with completeness > 99% and contamination < 1%. GTDB-Tk v.2.1.0^61^ was used to classify assemblies to the strain level. Thereby, we identified several *B. bifidum*, *B. longum* subsp. *longum*, and *B. breve,* and selected one representative respectively for our growth assays.

Bifidobacterium isolates were pre-grown in MRS-medium + cysteine (0.1% w/v; Sigma-Aldrich) for 48 hours. Thereof, 2% (v/v) was transferred to modified M9 medium^28^ with 0.1% or 0.5% 2’-O-fucosyl-lactose (2’FL; Glycome, IRE), following a 24 hours incubation at 37°C under anaerobic conditions (Coy Laboratory Products, USA) to probe their capabilities of metabolizing 2’FL. Modified M9 had the following additions: biotin (10mg/l), para-aminobenzoic acid (10mg/l), thiamine (400mg/l), nicotinic acid (400mg/l), pyridoxine (400mg/l), pantothenate calcium (200 mg/l), riboflavin (200mg/l), and a diluted casamino-acid extract (0.01% w/v; Sigma-Aldrich). pH was adjusted to 6.3 and media were filter-sterilized (Sartorius 0.22µm filter; Sigma-Aldrich) before inoculation. *E. coli* K12 was pre-grown in brain-heart infusion (BHI; Sigma Aldrich) for 24 hours before being inoculated (1% v/v) in m-M9 or m-M9 spent by *B. bifidum*. To generate supernatants of spent media, cultivates were centrifuged at 13.000 rpm for 10 minutes, supernatants were collected, pH was adjusted to 6.3, and lastly filter-sterilized (Sartorius 0.22µm filter; Sigma-Aldrich). Growth of all cultures was monitored under anaerobic conditions in a plate-reader (Thermo Multiskan, Thermo Fisher, USA). For end-point assessments, optical density (OD-600) of cultures was measured by spectrophotometry (Camspec, Spectronic, UK). E. coli BW25113 Δ-cysE was obtained from the Keio collection^62^. For co-culture experiments, *B. bifidum* and *E. coli* K12 were pre-grown in m-MRS or BHI for 48 or 24 hours under anaerobic conditions at 37°C, respectively. Cell density of inoculates were assessed by Luna FX7 automated cell counter (Thermo Fisher USA) to ensure close to 1:1 ratios of all inoculates. *E. coli* Δ-cysE was pre-grown in BHI + cysteine. Relative abundances of *B. bifidum* and *E. coli* K12 in co-culture were assessed by fluorescent gram-stain (LIVE BacLight, Thermo Fisher, USA) and manual counting. Nine representative pictures were taken per timepoint (Zeiss, GER) to estimate the average relative abundance of either microbe in co-culture (supplementary file 1).

### Quantification of 2’-O-fucosyl-lactose and its monomers

For thin layer chromatography (TLC), Culture supernatant samples (3 μL) were spotted onto silica plates (Sigma Z740230) and resolved in running buffer containing 2:1:1 butanol, acetic acid and dH_2_O, respectively. TLCs were subsequently stained using diphenylamine–aniline– phosphoric acid stain^63^ and developed by heating.

To analyse and quantify monosaccharide release in culture supernatants, we used high-performance anion exchange chromatography with pulsed amperometric detection (HPAEC-PAD). Sugars were separated using a CarboPac PA-1 anion exchange column with a PA-1 guard using a Dionex ICS-6000 (Thermo Fisher) and detected using PAD. Flow was 0.75 mL min^−1^ and elution conditions were 0–25 min 5 mM NaOH and then 25–40 min 5– 100 mM NaOH. Software used was the Chromeleon Chromatography Data System. Monosaccharide/disaccharide standards were included at the following concentrations: Maltose = 0.1 mg/ml, Glucose = 0.1 mM, Fucose = 0.1, 0.075, 0.05, 0.025, and 0.01mM, N-acetylglucosamine = 0.2 mM, Galactose = 0.2 mM, N-acetylgalactosamin = 0.1 mM, and Lactose 0.2 mM. All data were obtained by diluting supernatant samples 1/10 prior to injection.

### Analytics and data visualization

Statistical analysis (Student’s T-test, ANOVA, repeated measures ANOVA, Wilcoxon Test, Chi-square test, and Fisher’s exact test) was performed in ‘R version 4.0’ and the R package ‘rstatix version 0.7.0’^64^. All p-values were adjusted using Bonferroni’s method. Data was visualised via ‘R version 4.0’ and R package ‘ggplot2 version 3.3.340’^65^. ‘Ampvis2’^66^ and ‘phyloseq’^67^ were used for handling of metagenomic counts, metadata, and taxonomy files, including species filtering, analysis of alpha and beta diversity, as well as principal component analysis (PCA). ‘co_occurrence()’ in phylosmith^68^ was used for co-occurrence analysis of microbial species. For randomization of matrices, we used the ‘randomizeMatrix()’ function in ‘Picante’ ^69^ with ‘null.model = ‘frequency’’ over 10.000 iterations. Kolmogorow-Smirnov-Test – ‘ks.test()’ in base R – was used to test for whether differences in randomized and observed matrices were statistically different. ‘c.score()’ function in ‘bipartite’^70^ was used for calculation of checkerboard indices. For Mean Across Jaccard Index Checkerboards (MAJIC), randomized and observed matrices were transformed into presence/absence matrices using ‘phyloseq_standardize_otu_abundance()’ with ‘method = pa’ in ‘metagMisc’^71^, and average Jaccard dissimilarities were calculated using ‘vegdist()’ in ‘vegan’^72^. MAJIC was coded ‘R version 4.0’, looping through focal species 𝞼_i_ to obtain 𝞼_i_^+^ (µ_i_ ^+^), and in 𝞼_i_^-^ (µ_i_^-^) - as described in Figure S2A.

## Resource availability

### Lead contact

Further information for resources and reagents should be directed to and will be fulfilled by lead contacts David Seki or Lindsay J. Hall.

### Materials availability

Infant faecal sample metagenome sequencing raw reads are publicly available in the NCBI Sequence Read Archive (SRA) under accession no. PRJNA1230889.

## Acknowledgments

We thank all families for their participation during this study. We would like to thank the QIB sequencing team for technical assistance with shotgun metagenomics library preparation. This project was funded by the Wellcome Trust Investigator Award no. 220876/Z/20/Z to LJH, and a Biotechnology and Biological Sciences Research Council (BBSRC) Institute Strategic Programme, Gut Microbes and Health BB/R012490/1, and its constituent projects BBS/E/F/000PR10353 and BBS/E/F/ 000PR10356, and by the BBSRC Institute Strategic Programme Food Microbiome and Health BB/X011054/1 and its constituent project BBS/E/F/000PR13631 to LJH. The computational results of this work have been achieved using the Life Science Compute Cluster (LiSC) of the University of Vienna.

## Author contributions

Conceptualization, D.S. and L.J.H; methodology, D.S., M.K., R.K., S.P., L.C., C.B., A.AG.; software, D.S., S.P; investigation, D.S.; writing – original draft, D.S. and L.J.H.; writing – review & editing, D.S., L.J.H., S.P.; resources, L.J.H., L.C., N.vB., J.P., P.C., M.M; and funding acquisition, L.J.H.

---

Supplementary Figure 1:

A) Boxplots summarizing Bray-Curtis distances of pairwise comparisons between microbiota of infants and parenting or non-related mothers. B) C-score distribution for observed species occurrences in given age-groups. C) Heatmap displaying mean average abundance of species in given age groups. Blue colour indicates low mean average abundance, orange colour indicates high mean average abundance. D) Co-occurrence analysis highlighting significant correlations (adjusted p-values < 0.05) between microbial species.

Supplementary Figure 2:

A) Schematic illustration of computational pipeline. B) Distribution of average Jaccard dissimilarities (µ) in presence of state-associated (X) or state-dissociated (O) focal species (𝞼_i_). C) For each focal species (𝞼_i_), we identify differences in abundance patterns of any other species (j) between samples where focal species i (𝞼_i_^+^) is present and samples where it’s absent (𝞼_i_). Bar-chart summarizes these significant differences.

Supplementary Figure 3:

A) Scatter-plot between nucleotide diversity (π) and mean MAG coverage. Each dot represents a MAG, coloured by corresponding phylum. B) Summary of strain-sharing events where strains of *Bifidobacteria,* and *E. coli* are shared among same-aged individuals. C) Differences of pop-ANI between comparisons of infants with parenting and random mothers. D) Summary of strain-sharing events where identical strains of *Bifidobacteria,* and *E. coli* are shared between infants and parenting mothers.

Supplementary Figure 4:

Bar-charts summarizing the relative distribution of GH-2 co-evolving genes per genus across given kegg-orthology (ko) groups. Distribution of ko-groups among microbial genera significantly differed from shuffled random distributions.

Supplementary Figure 5:

A) No-carbon control: growth of *B. bifidum* (red), *B. breve* (salmon), *B. longum* (pink) in modified M9-medium (m-M9) without 2’-O-fucosyl-lactose (2’FL), measured by optical density (OD-600). B) No-amino-acid control: growth of *B. bifidum* in m-M9 + 2’FL lacking cas-amino-acids (brown), cysteine (yellow), or both (orange), measured by OD-600. C) Growth of *E. coli* in m-M9 and respective controls. D) Growth of E. coli on 2’FL monomers fucose (light-green), galactose (light-blue), and glucose (dark-blue). E) Growth of *E. coli* in supernatant of m-M9 + 2’FL previously spent by *B. bifidum* at pH 6,3 and 4,1. F) Co-cultures of *E. coli* and *B. bifidum* in m-M9 with and without each additive. Dots in line-plots show group median, error bars show standard deviation.

Supplementary Figure 6:

Thin layer chromatography (TLC) and matching high-pressure anion-exchange chromatography (HPAEC) for quantification of 2’-O-fucosyl-lactose (2’FL), its monomers, and lactose. A) Supernatants of *B. bifidum* grown in m-M9 with 0.5% 2’FL at pH 6,3. B) Supernatants of *E. coli* grown in *B. bifidum* spent m-M9 with 0.5% 2’FL at pH 6,3. C) Supernatants of *B. bifidum* grown in m-M9 with 0.5% 2’FL at pH 4,1. D) Supernatants of *E. coli* grown in *B. bifidum* spent m-M9 with 0.5% 2’FL at pH 4,1.

Supplementary Figure 1<u>: co-culture of *E. coli* and *B. bifidum*: Quantification of gram-stains.</u>

