## Supplementary figures and images for "Mutualistic interactions between *Escherichia coli* and *Bifidobacterium bifidum* enable degradation of human milk oligosaccharides in healthy infants"

### Figure S1

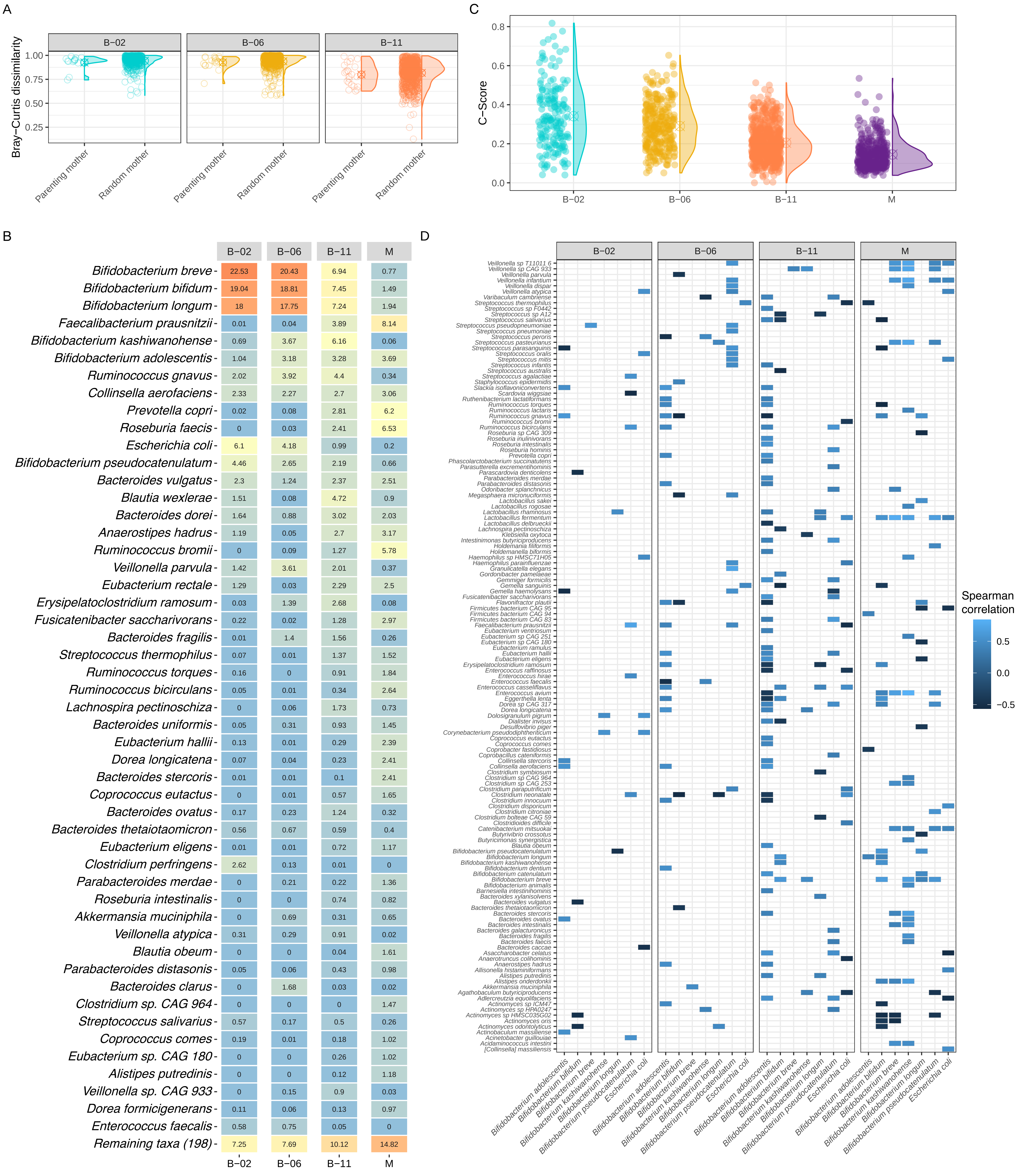

### Figure S2

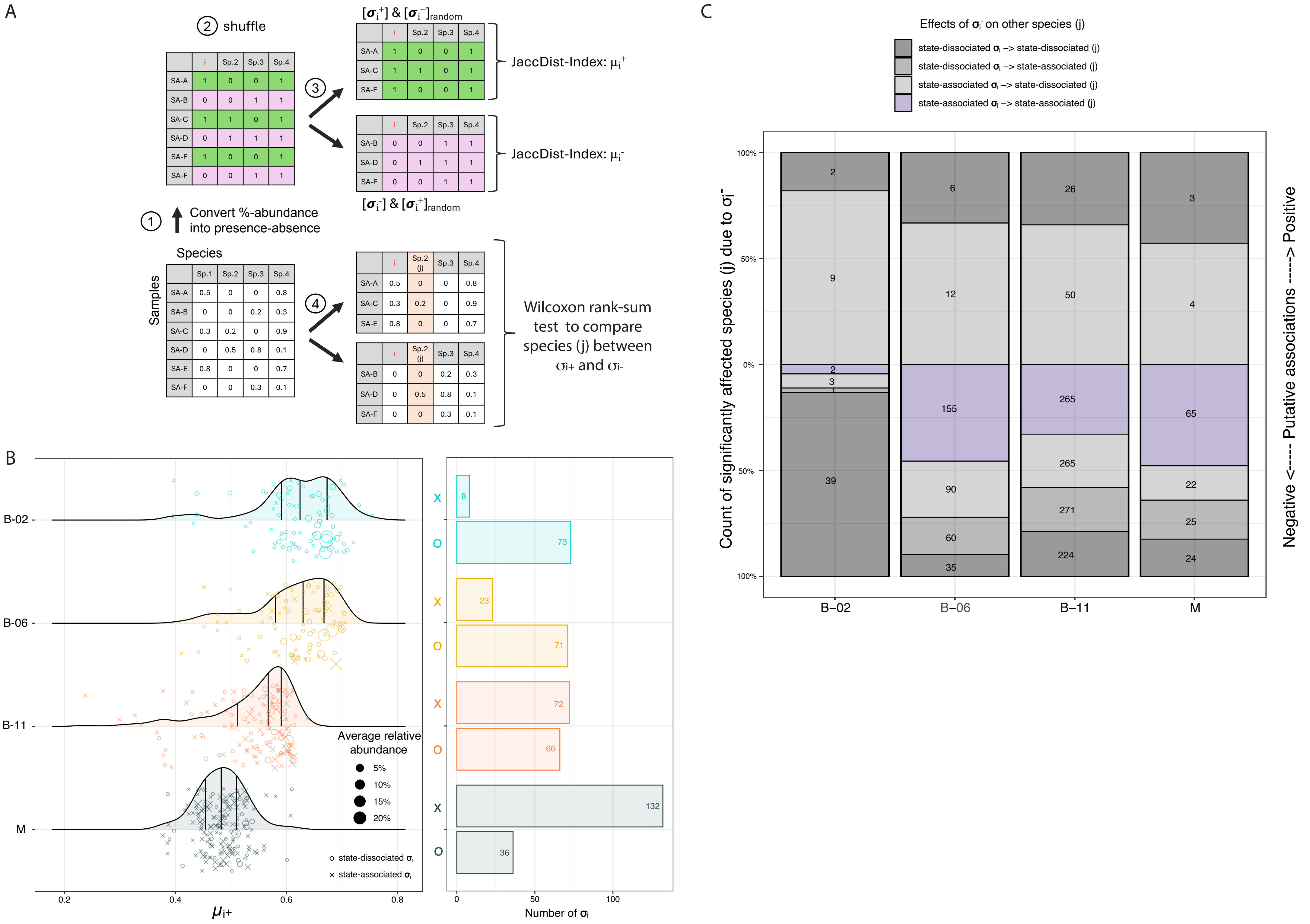

### Figure S3

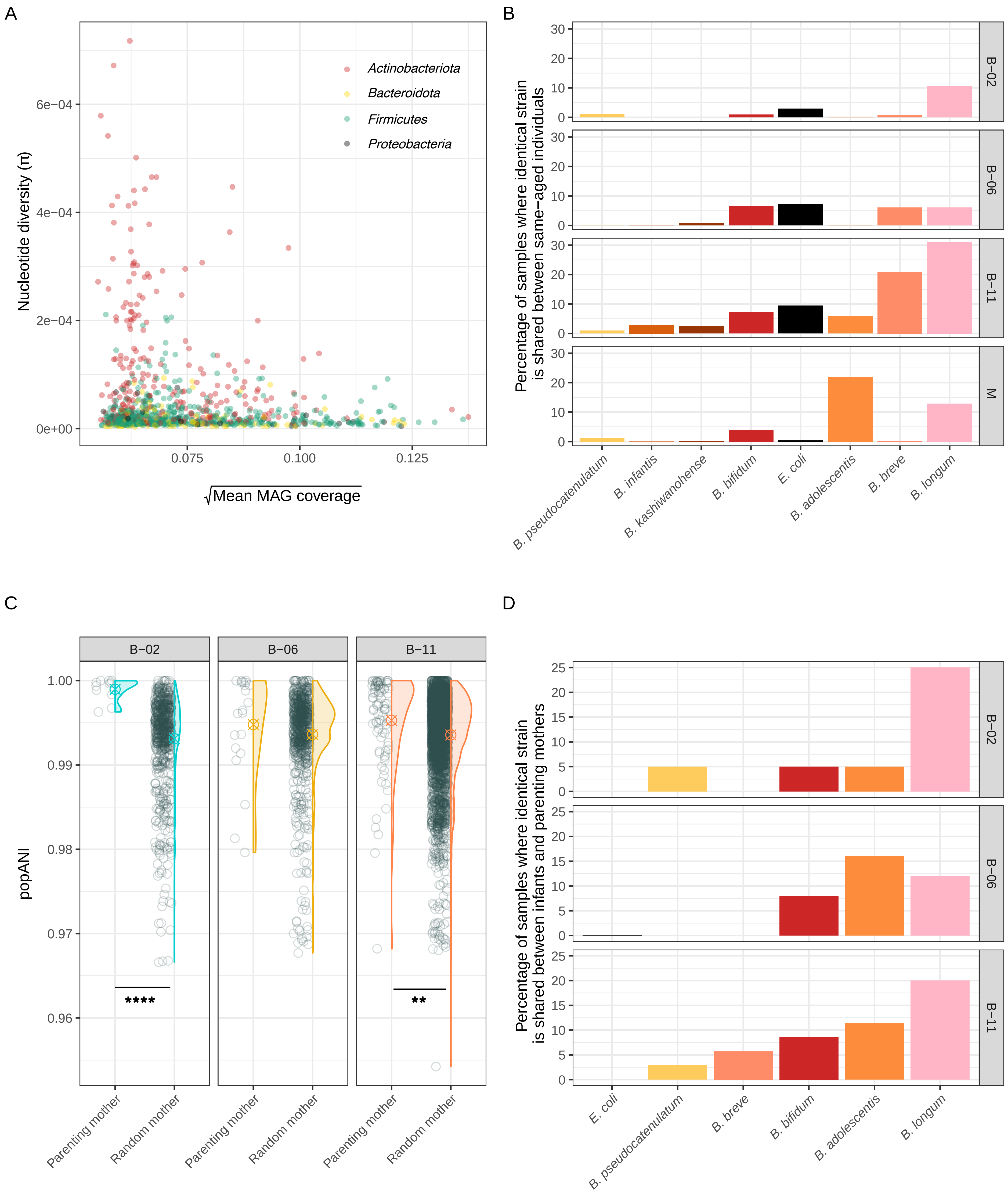

### Figure S4

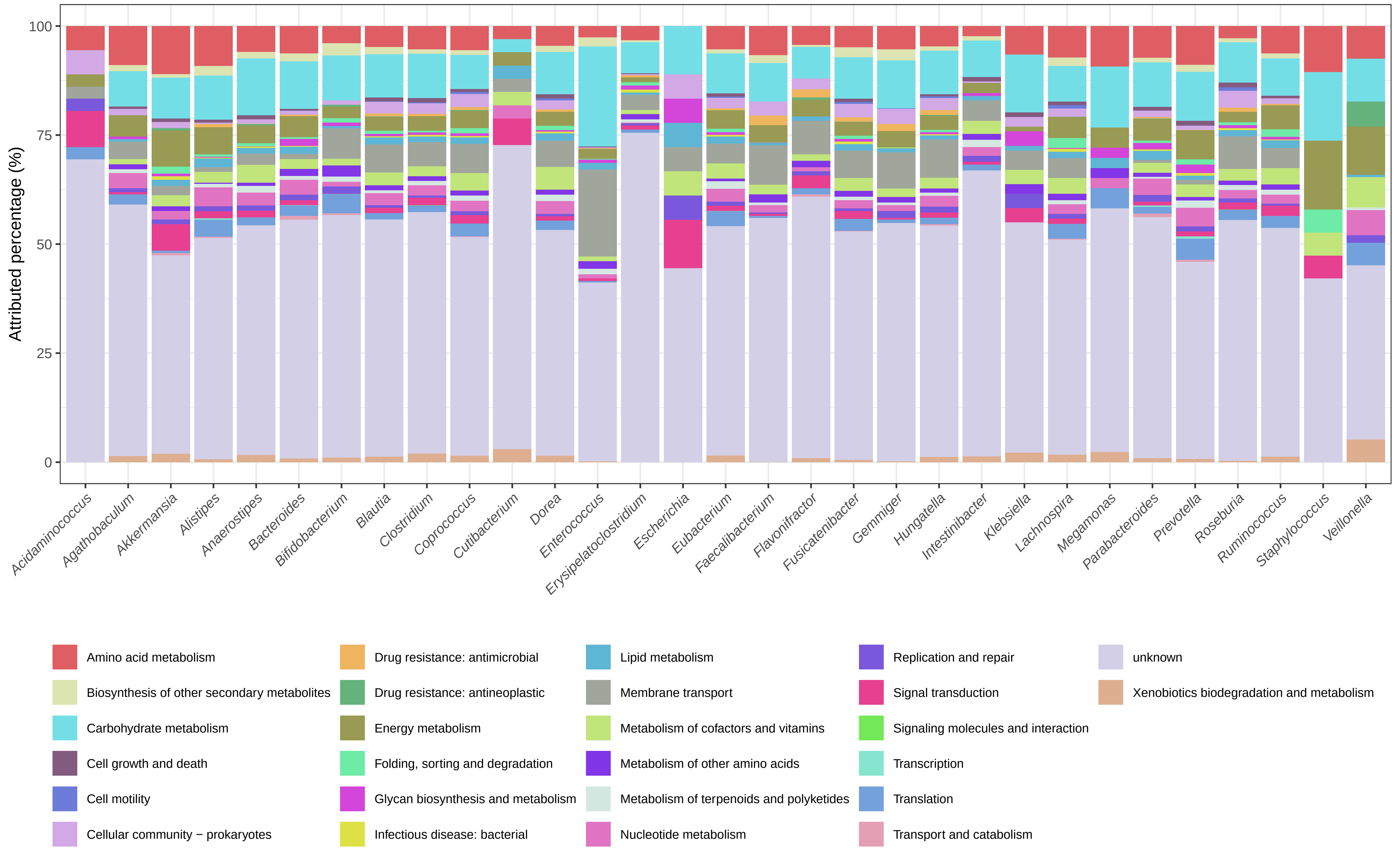

### Figure S5

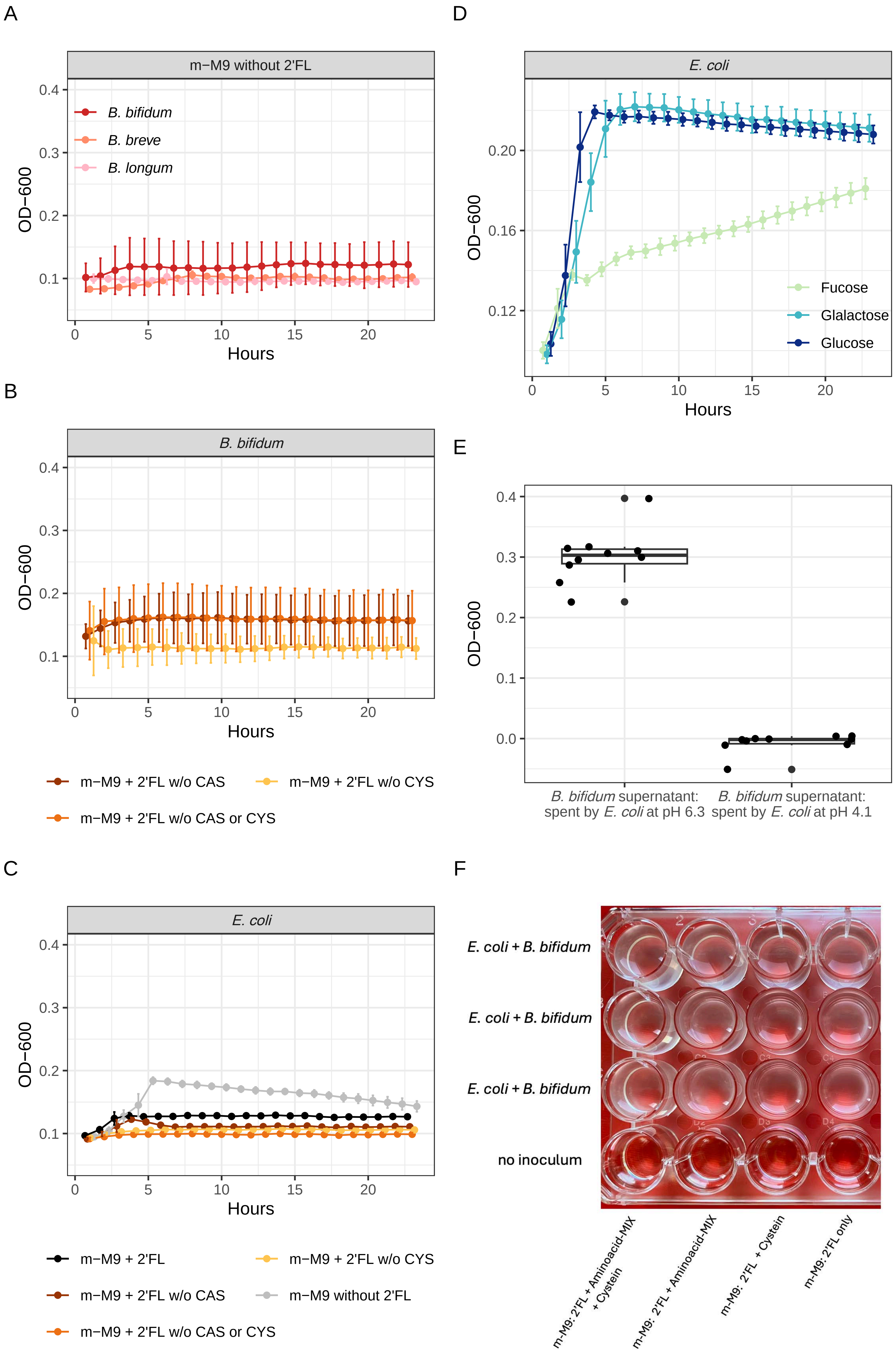

### Figure S6

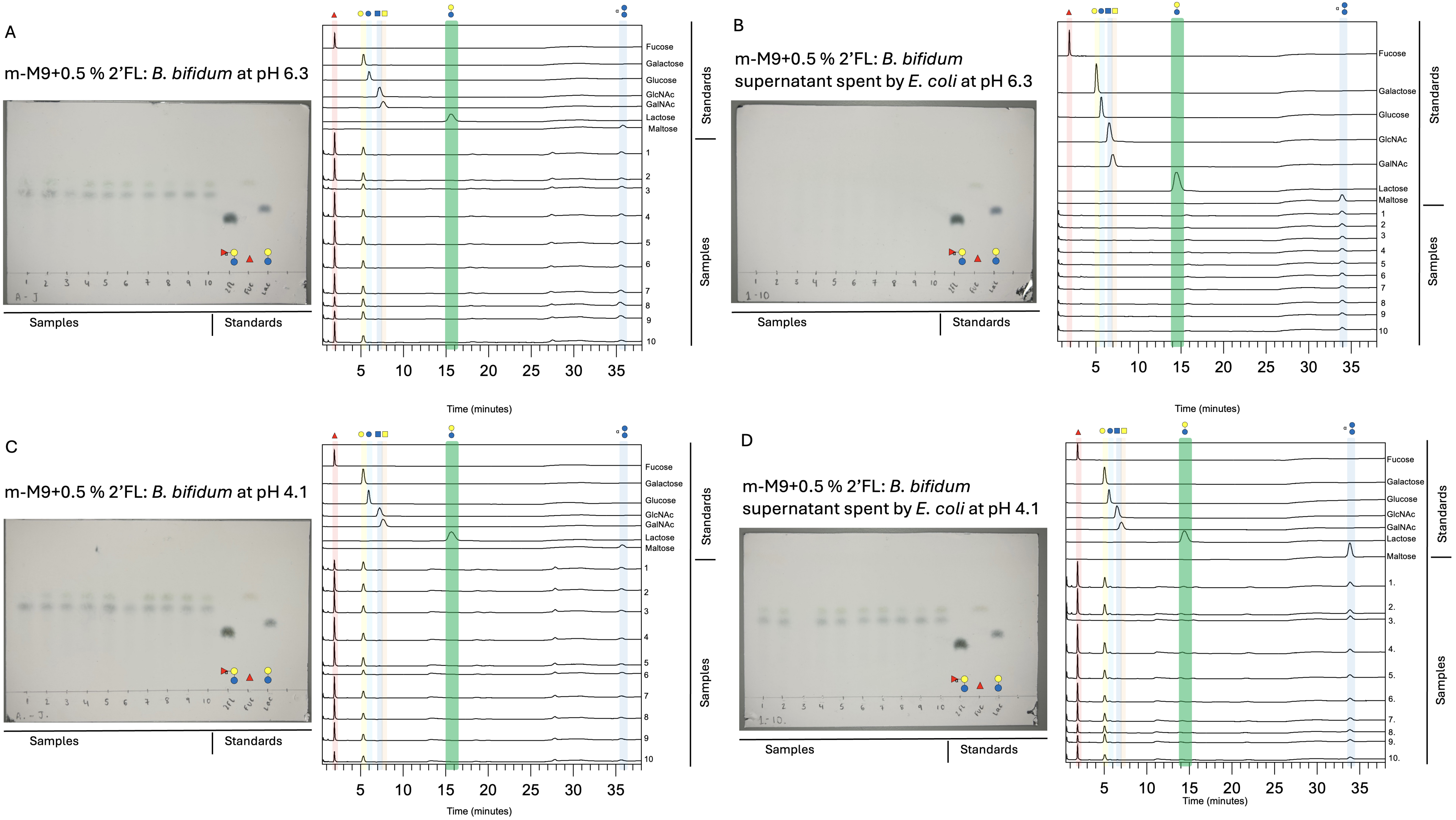
