## Supplementary File 1 for "Mutualistic interactions between *Escherichia coli* and *Bifidobacterium bifidum* enable degradation of human milk oligosaccharides in healthy infants"

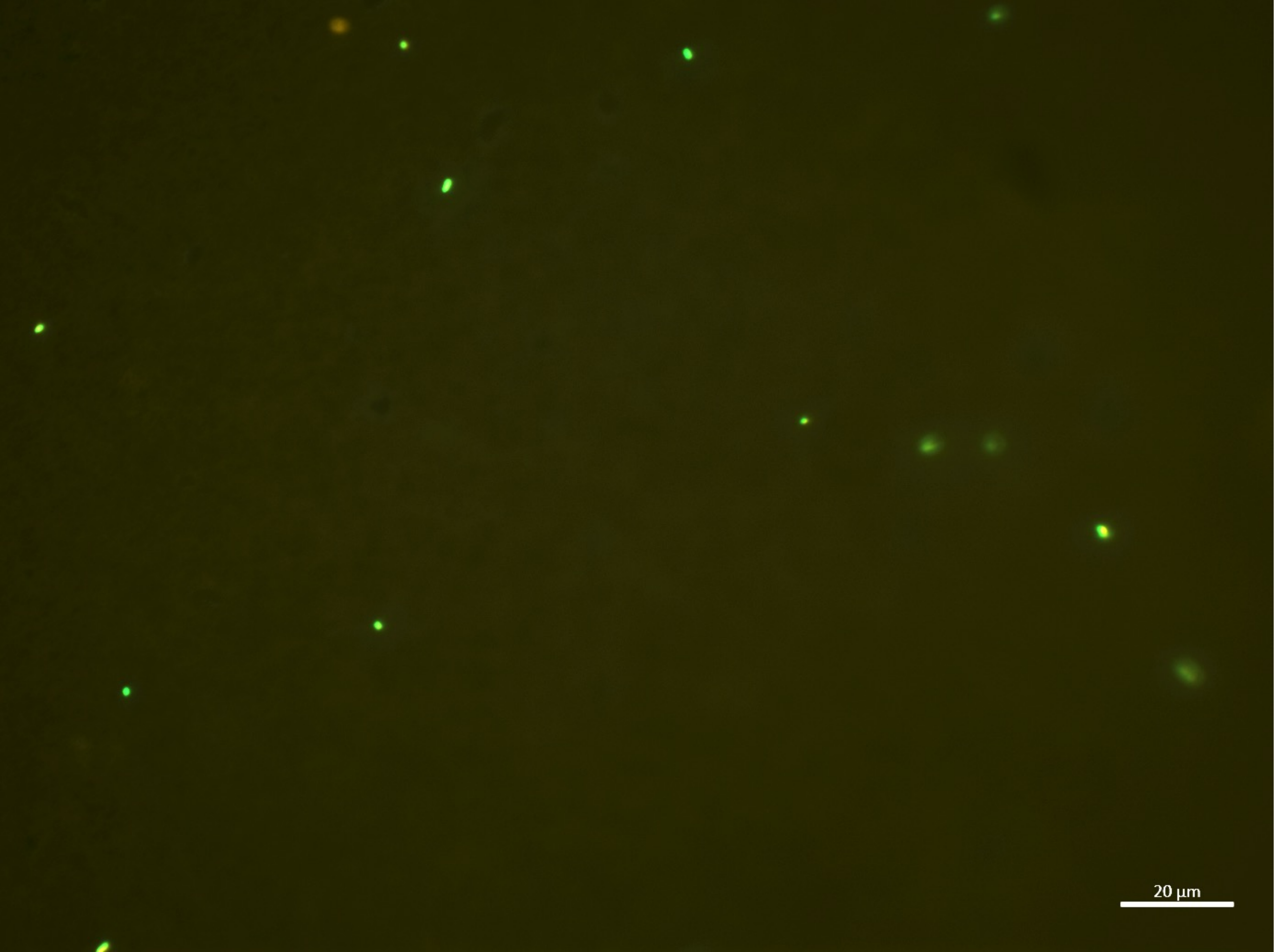

14:00-A

| | |
|---|---|
| **B.bifidum** | **1** |
| **E.coli** | **13** |
| **Sum** | **14** |

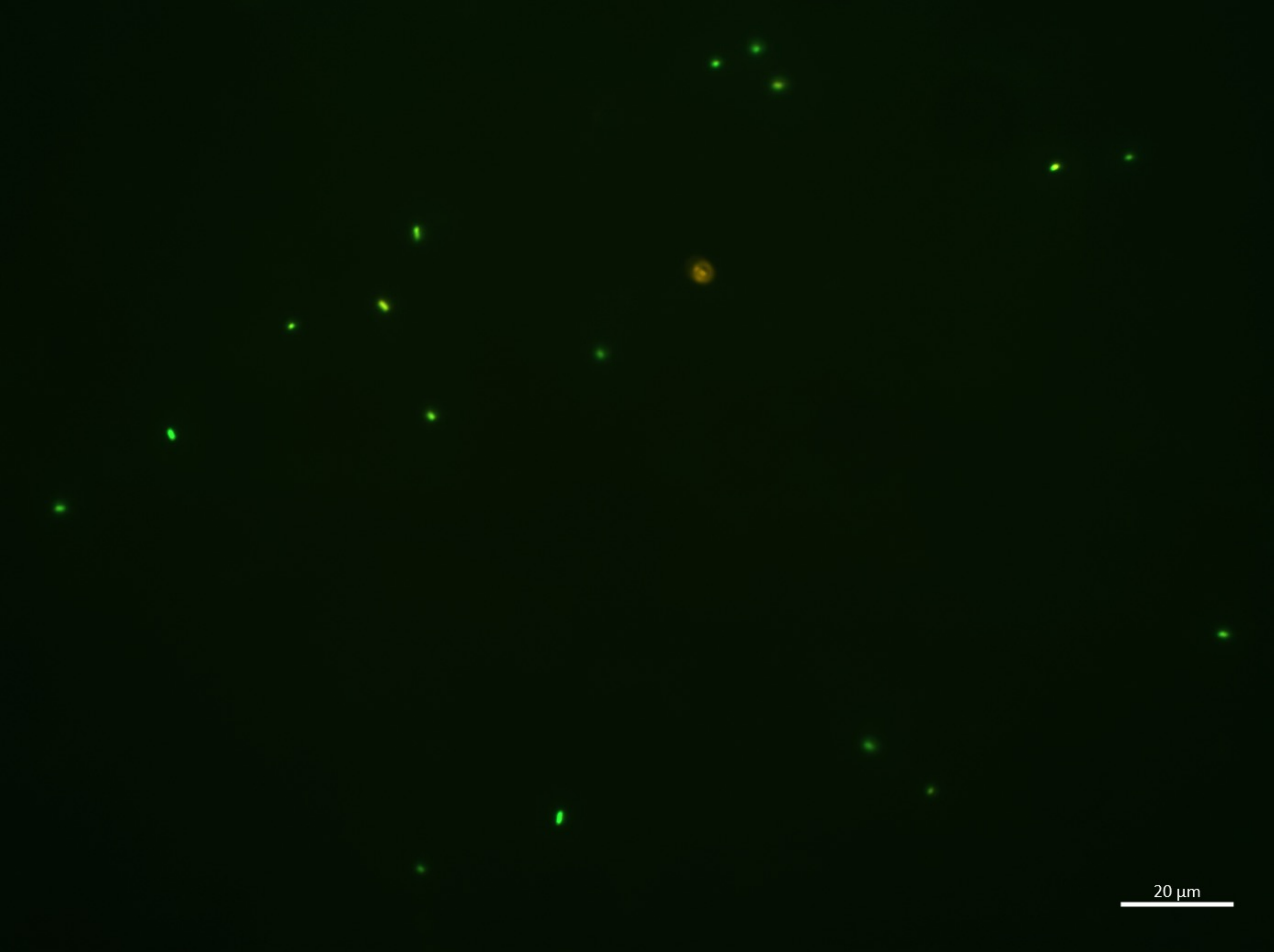

14:00-B

| | |
|---|---|
| **B.bifidum** | **1** |
| **E.coli** | 17 |
| **Sum** | 18 |

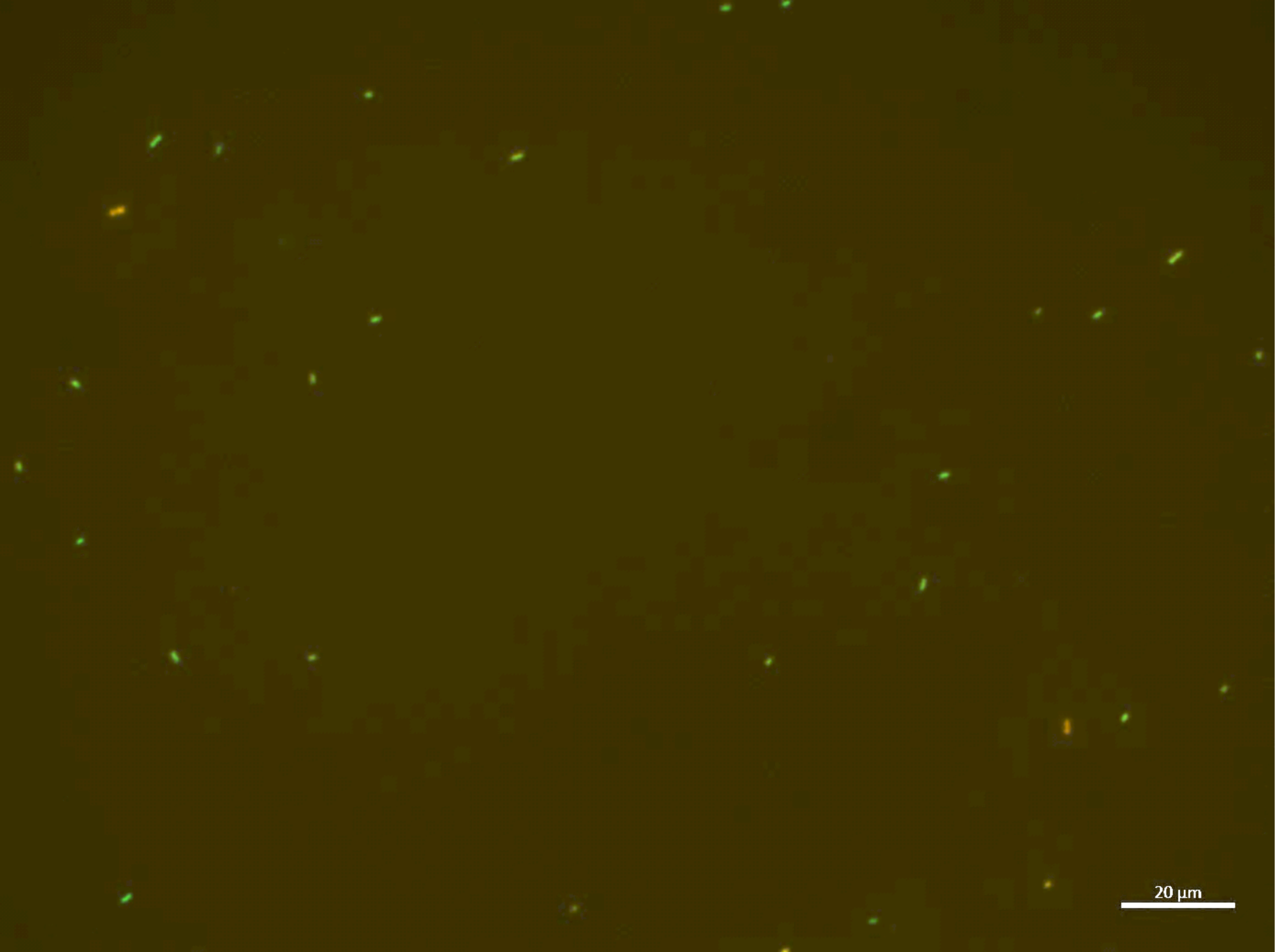

14:00-C

| | |
|---|---|
| **B.bifidum** | **3** |
| **E.coli** | 25 |
| **Sum** | 28 |

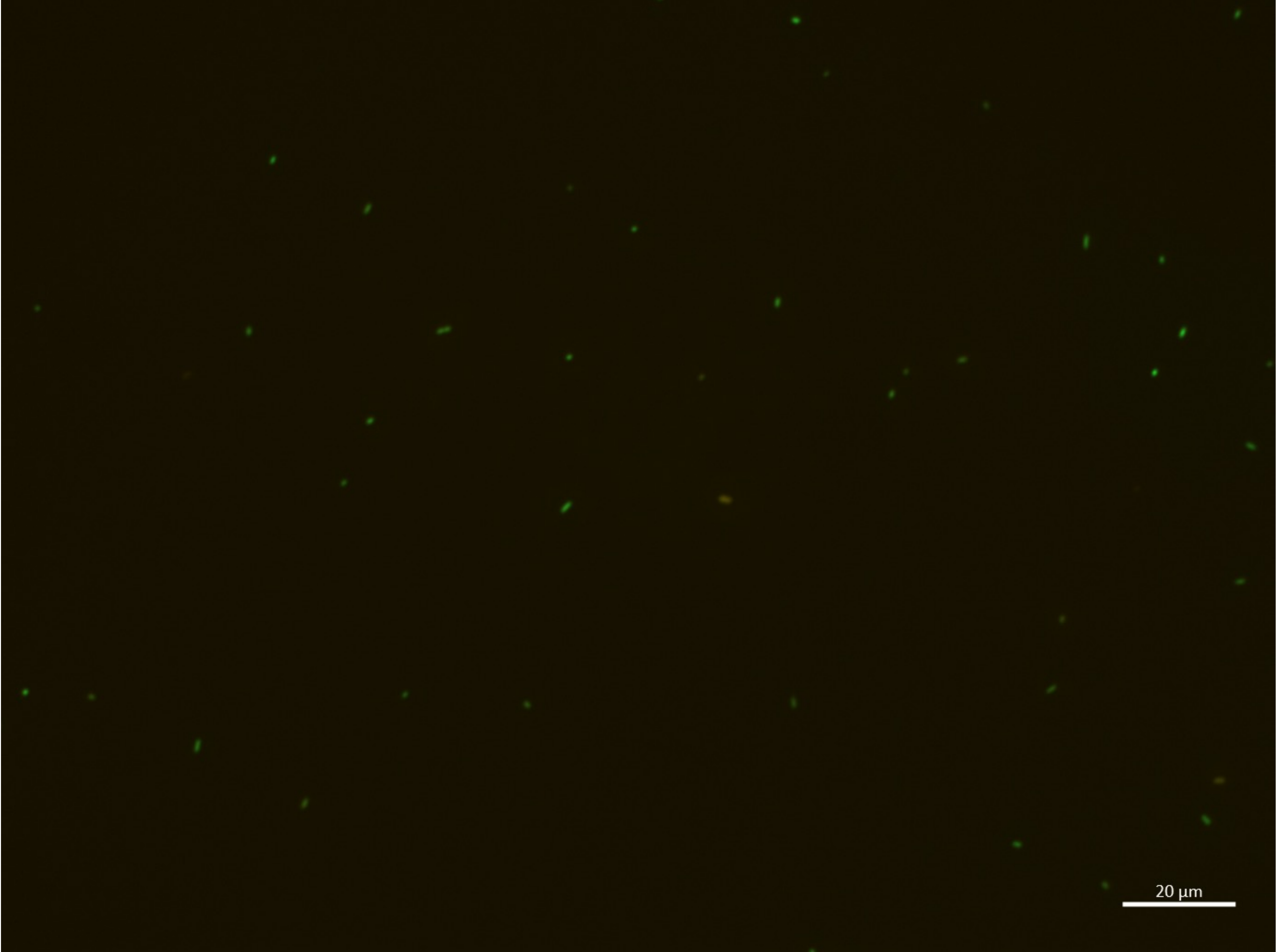

14:00-D

| | |
|---|---|
| B.bifidum | **3** |
| E.coli | 37 |
| **Sum** | 40 |

14:00-E

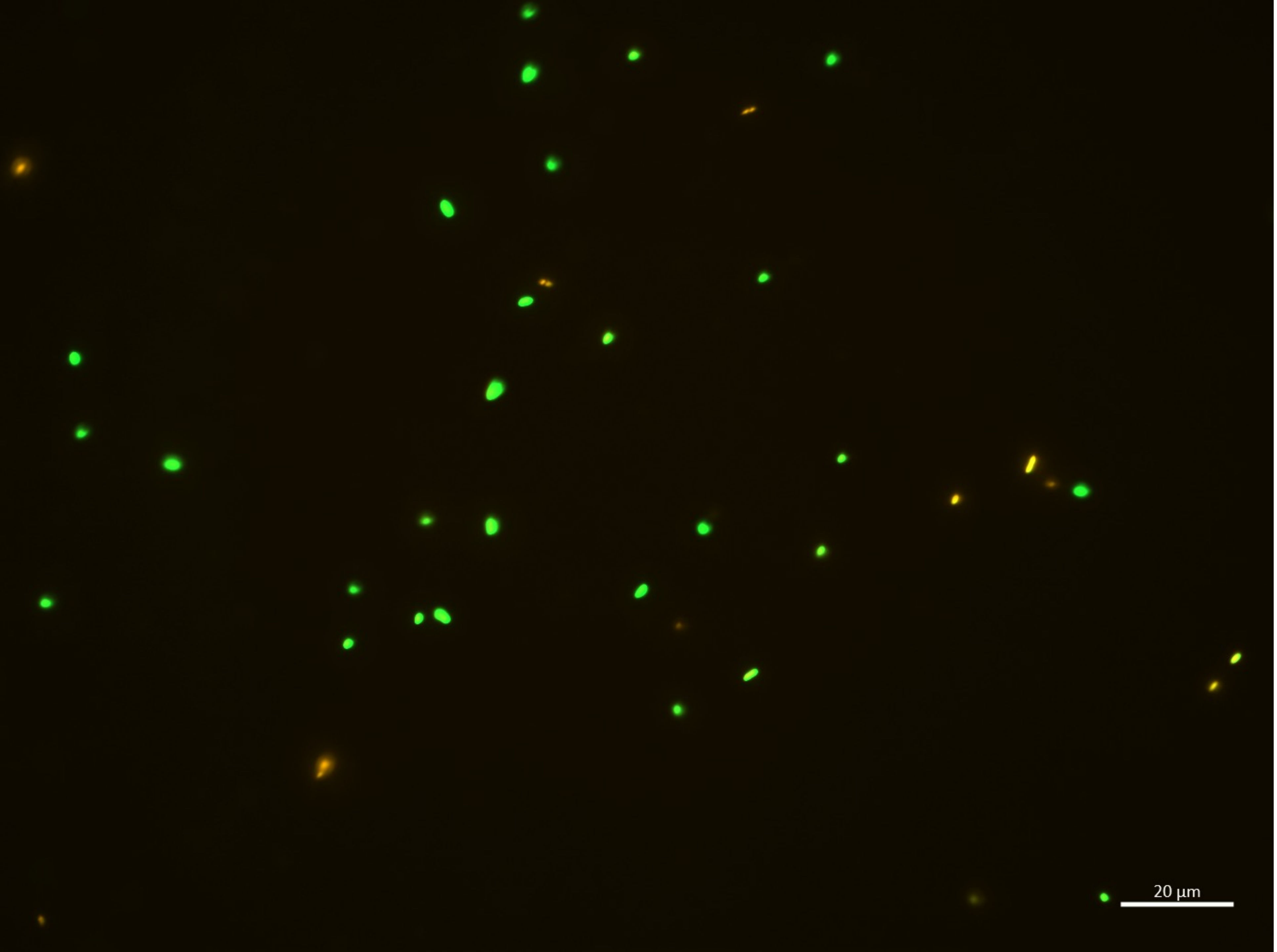

| | |
|---|---|
| **B.bifidum** | **11** |
| **E.coli** | 28 |
| **Sum** | 39 |

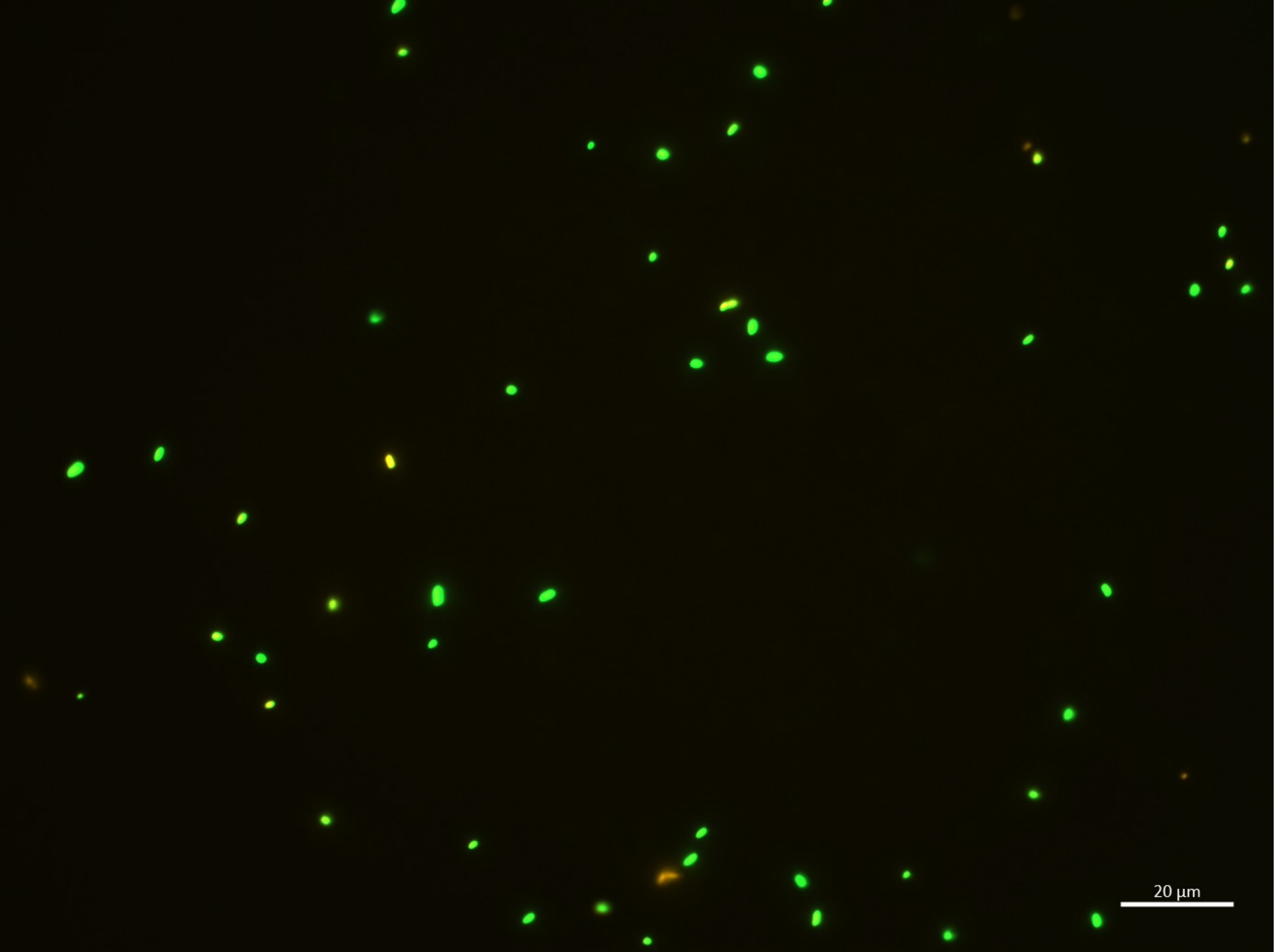

14:00-F

| | |
|---|---|
| B.bifidum | **6** |
| E.coli | 48 |
| **Sum** | 54 |

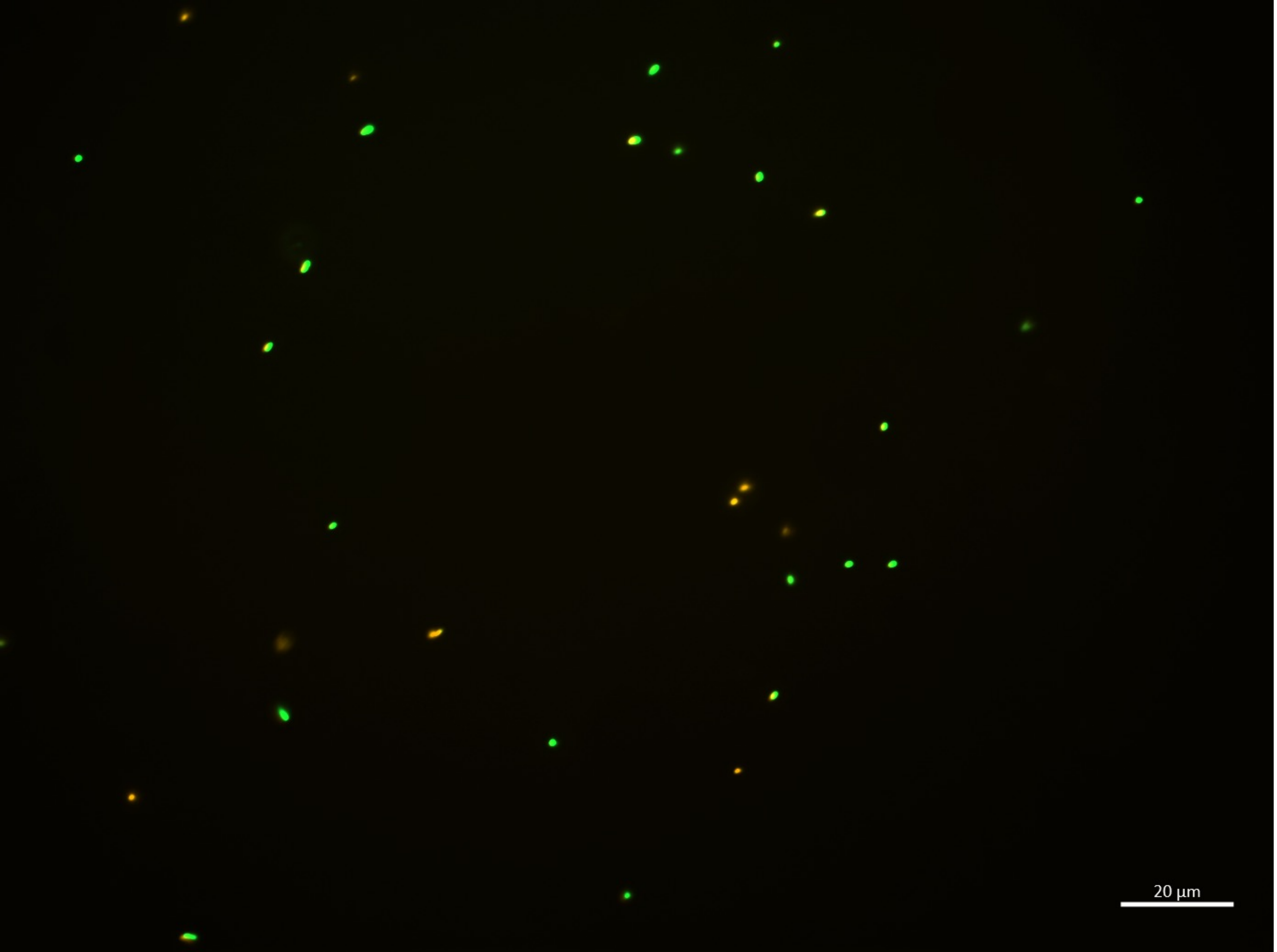

14:00-G

| | |
|---|---|
| B.bifidum | **9** |
| E.coli | 22 |
| **Sum** | 31 |

14:00-H

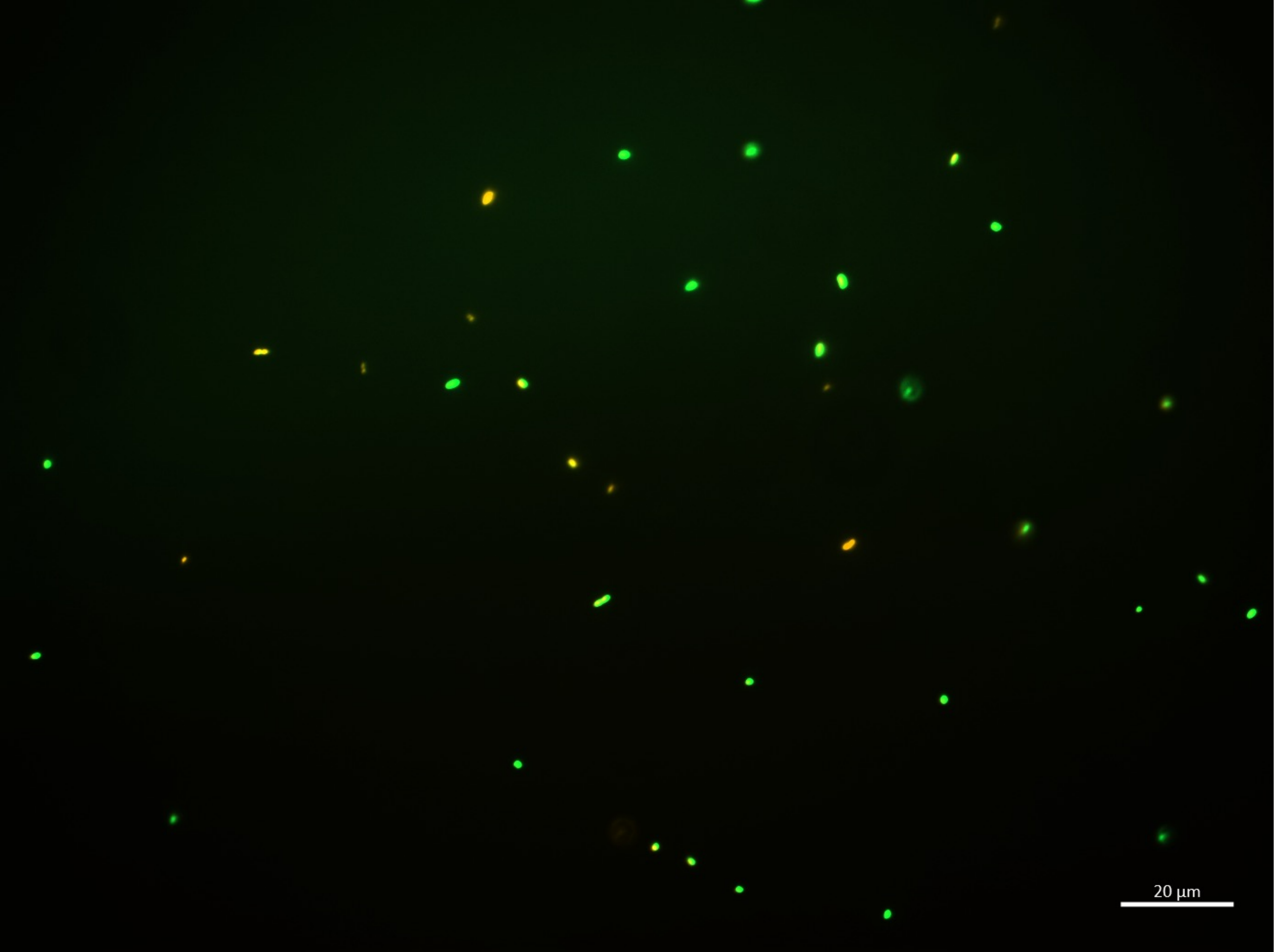

| | |
|---|---|
| B.bifidum | **8** |
| E.coli | 29 |
| **Sum** | 37 |

# 14:00-I

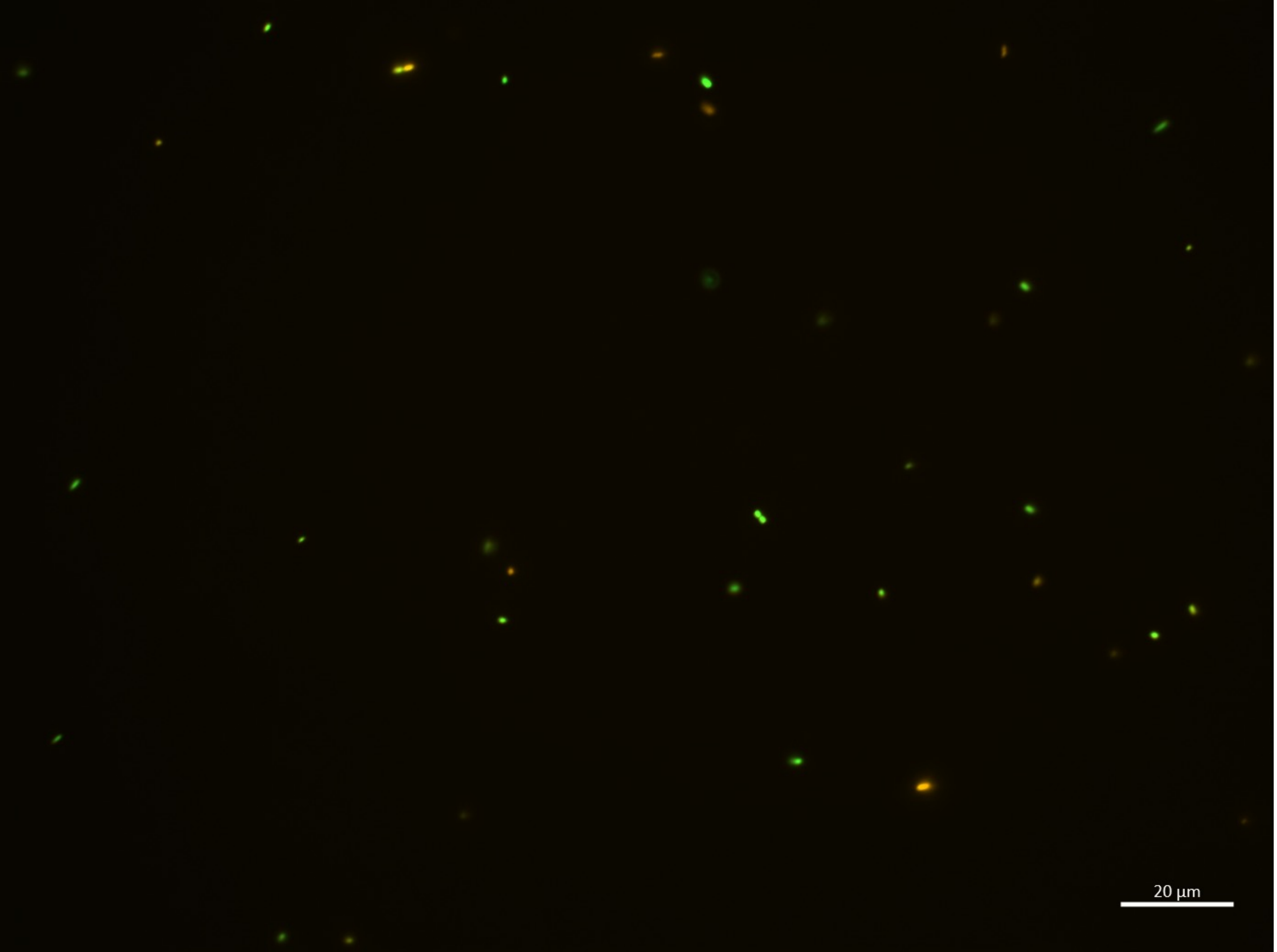

| | |
|---|---|
| B.bifidum | **7** |
| E.coli | 26 |
| **Sum** | 33 |

14:00-J

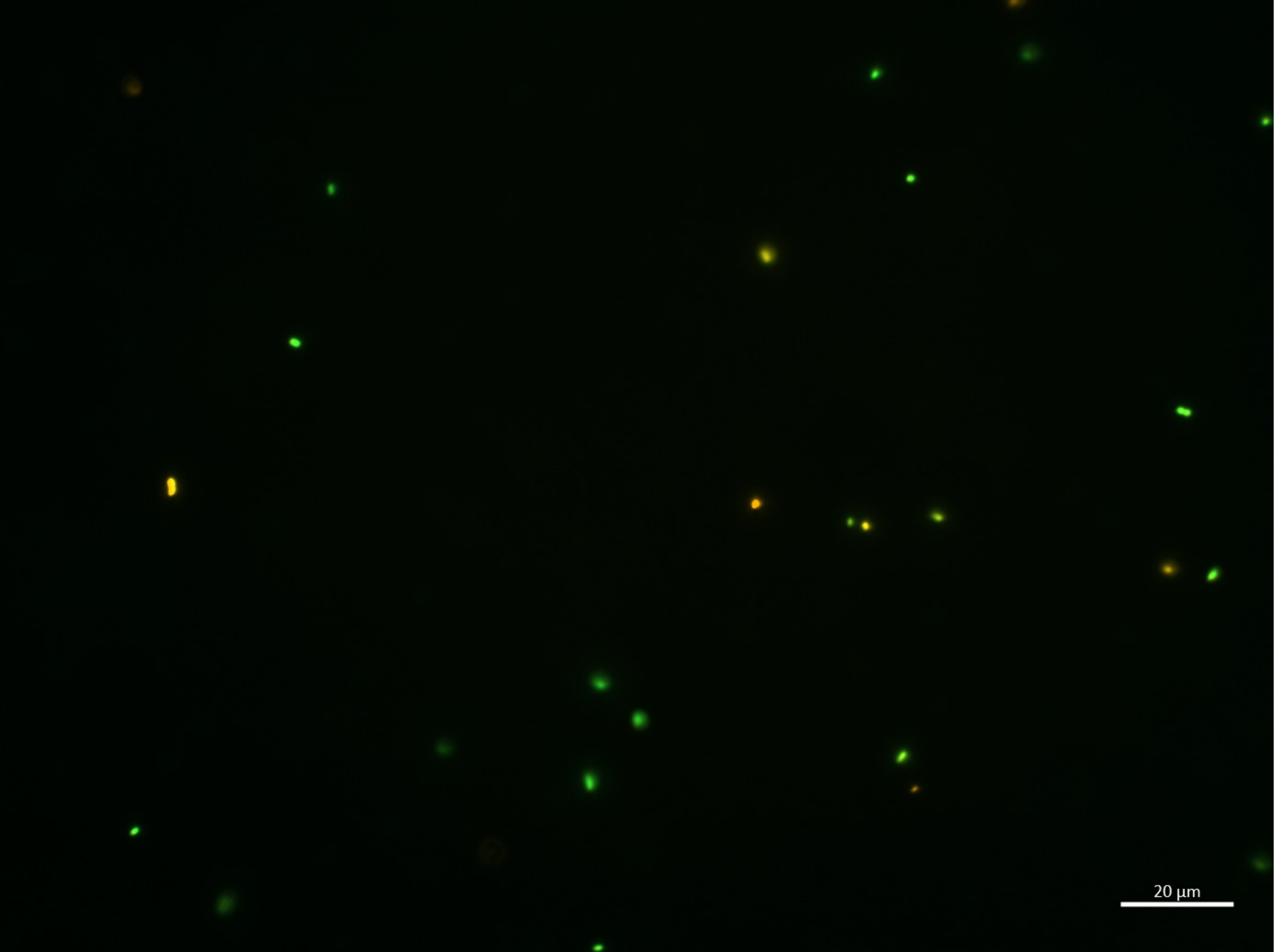

| B.bifidum | **7** |
|---|---|
| E.coli | 19 |
| **Sum** | 26 |

16:00-A

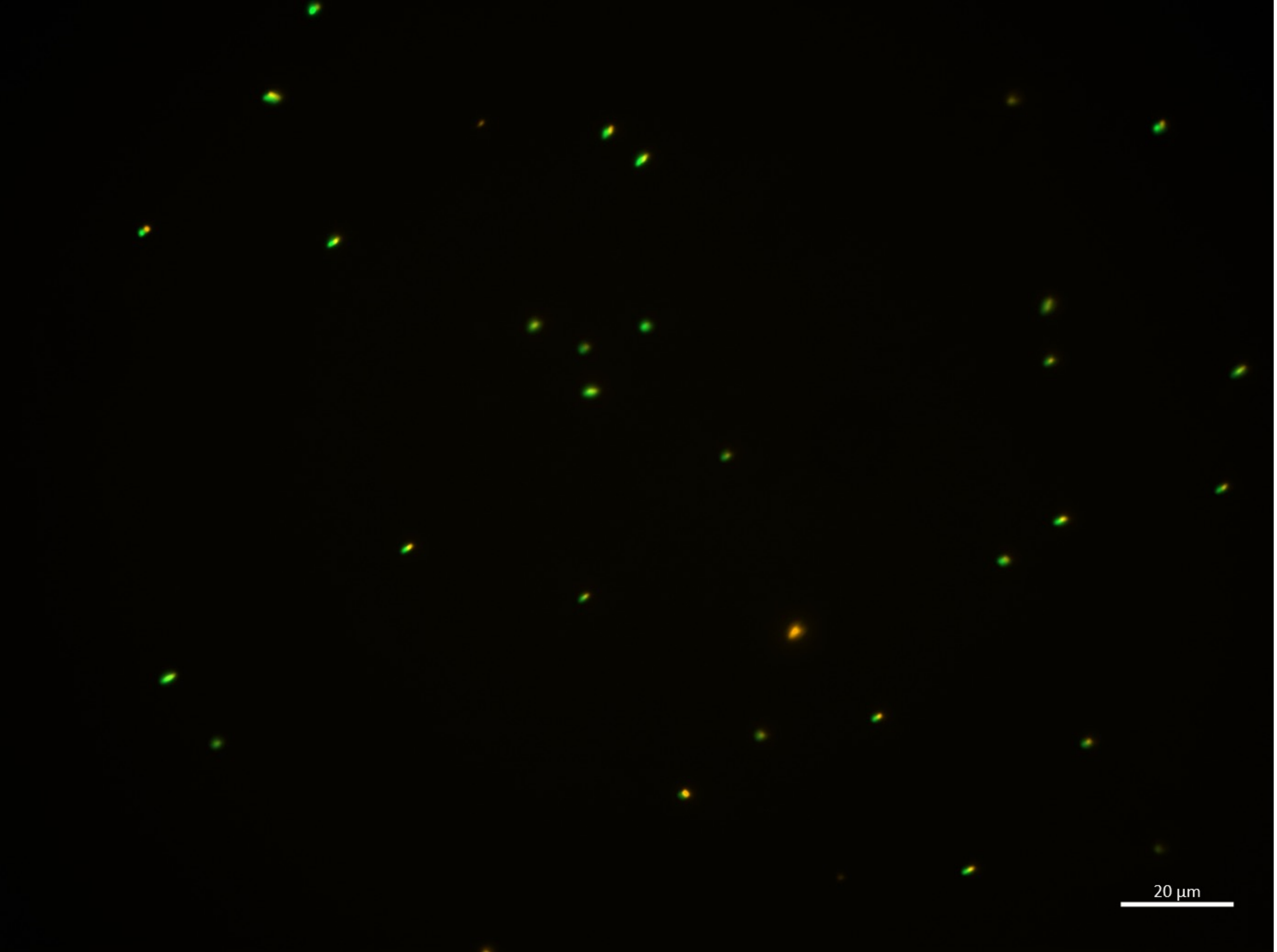

| | |
|---|---|
| B.bifidum | **4** |
| E.coli | 25 |
| **Sum** | 29 |

16:00-B

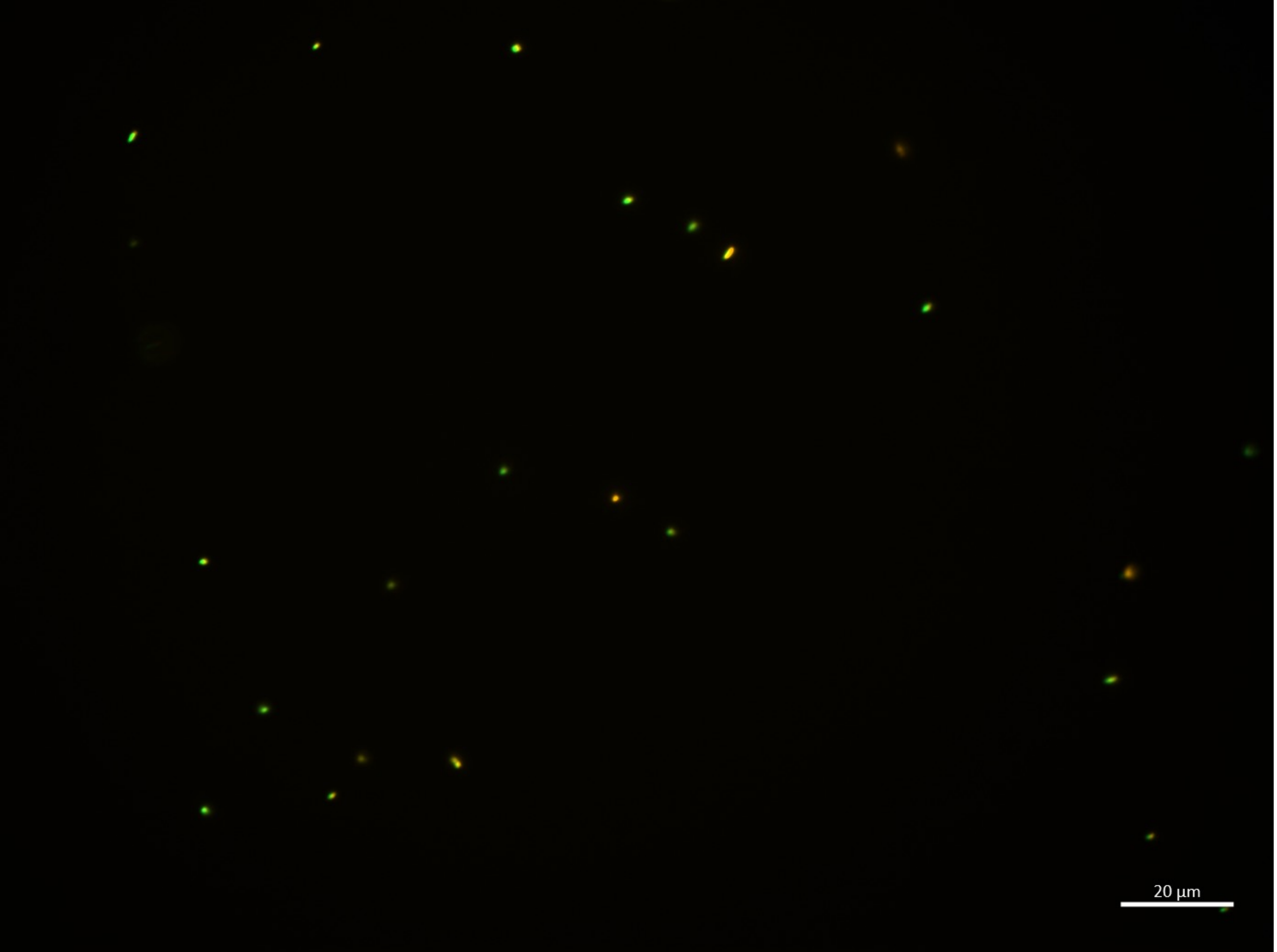

| | |
|---|---|
| B.bifidum | **4** |
| E.coli | 18 |
| **Sum** | 22 |

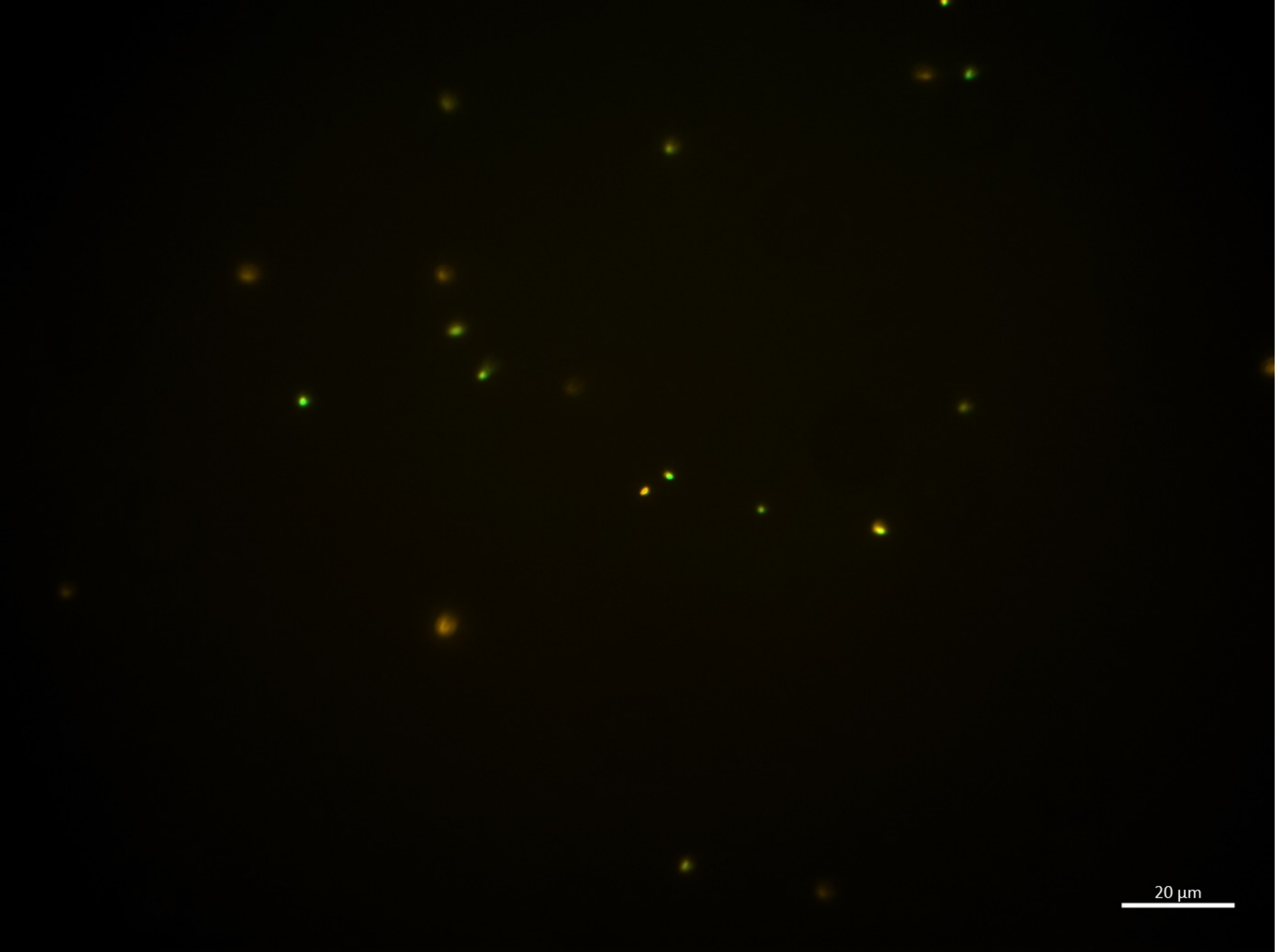

16:00-C

| | |
|---|---|
| **B.bifidum** | **11** |
| **E.coli** | 9 |
| **Sum** | 20 |

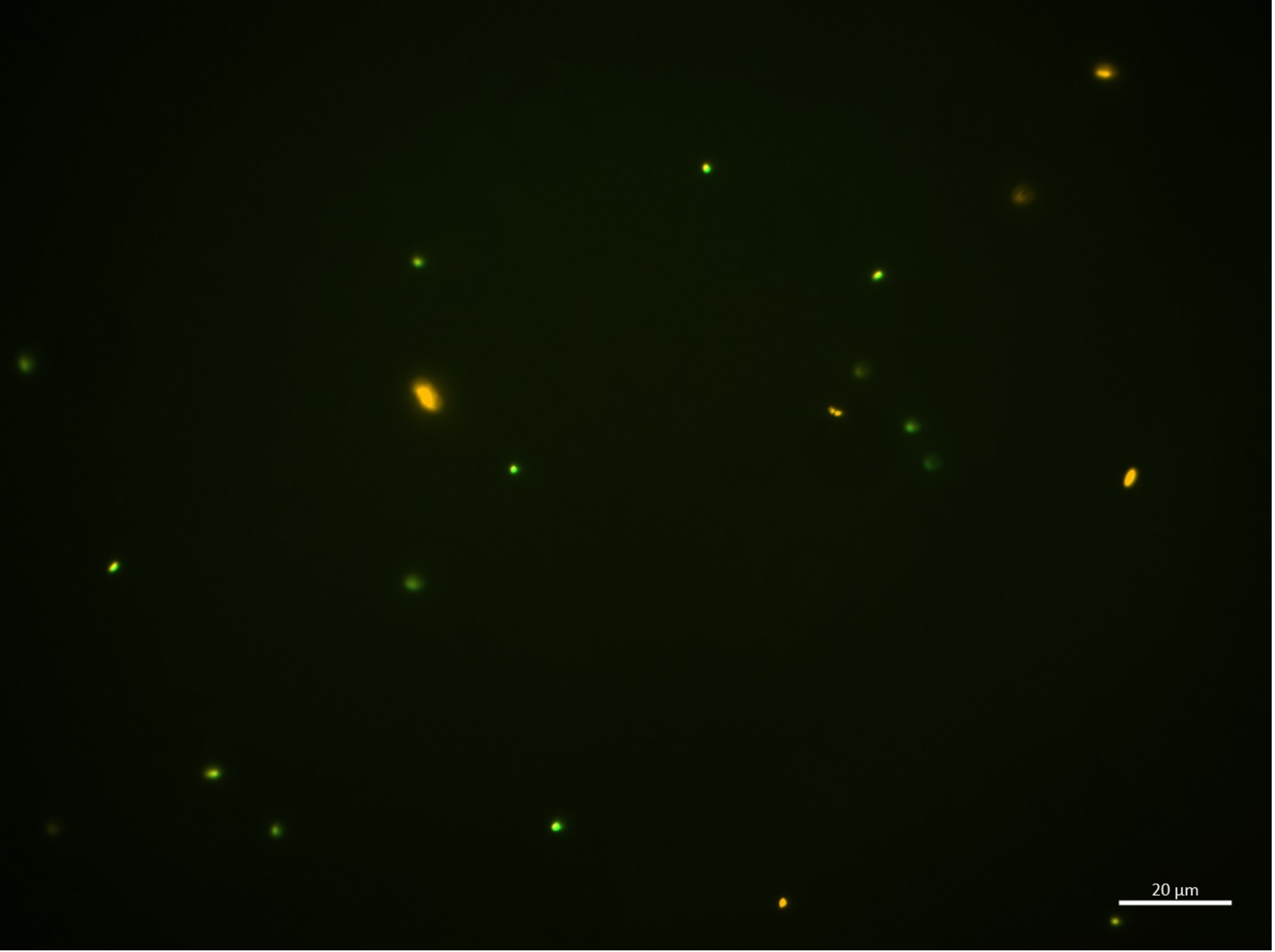

16:00-D

| | |
|---|---|
| **B.bifidum** | **6** |
| **E.coli** | 14 |
| **Sum** | 20 |

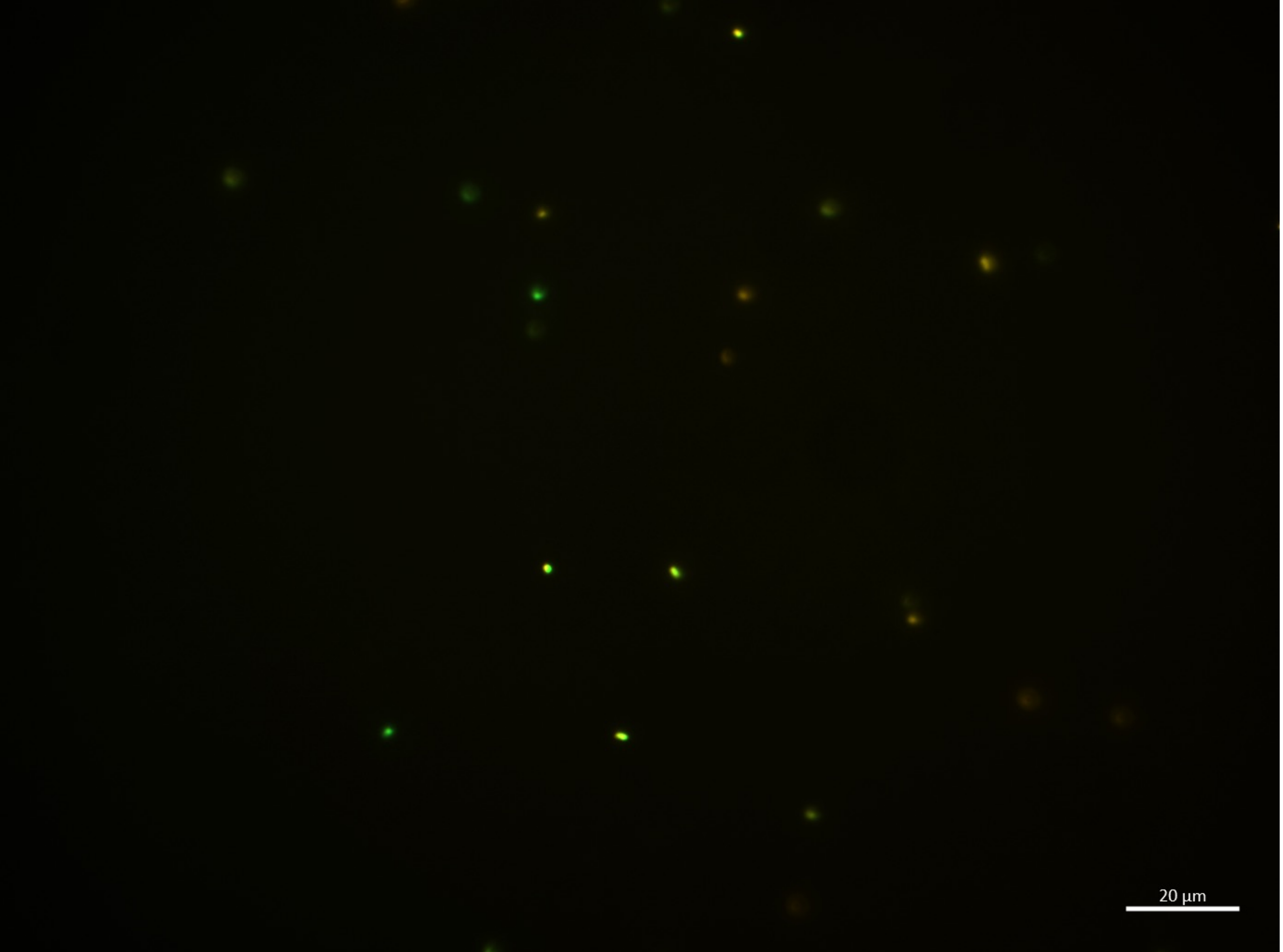

16:00-E

| | |
|---|---|
| **B.bifidum** | **6** |
| **E.coli** | 12 |
| **Sum** | 18 |

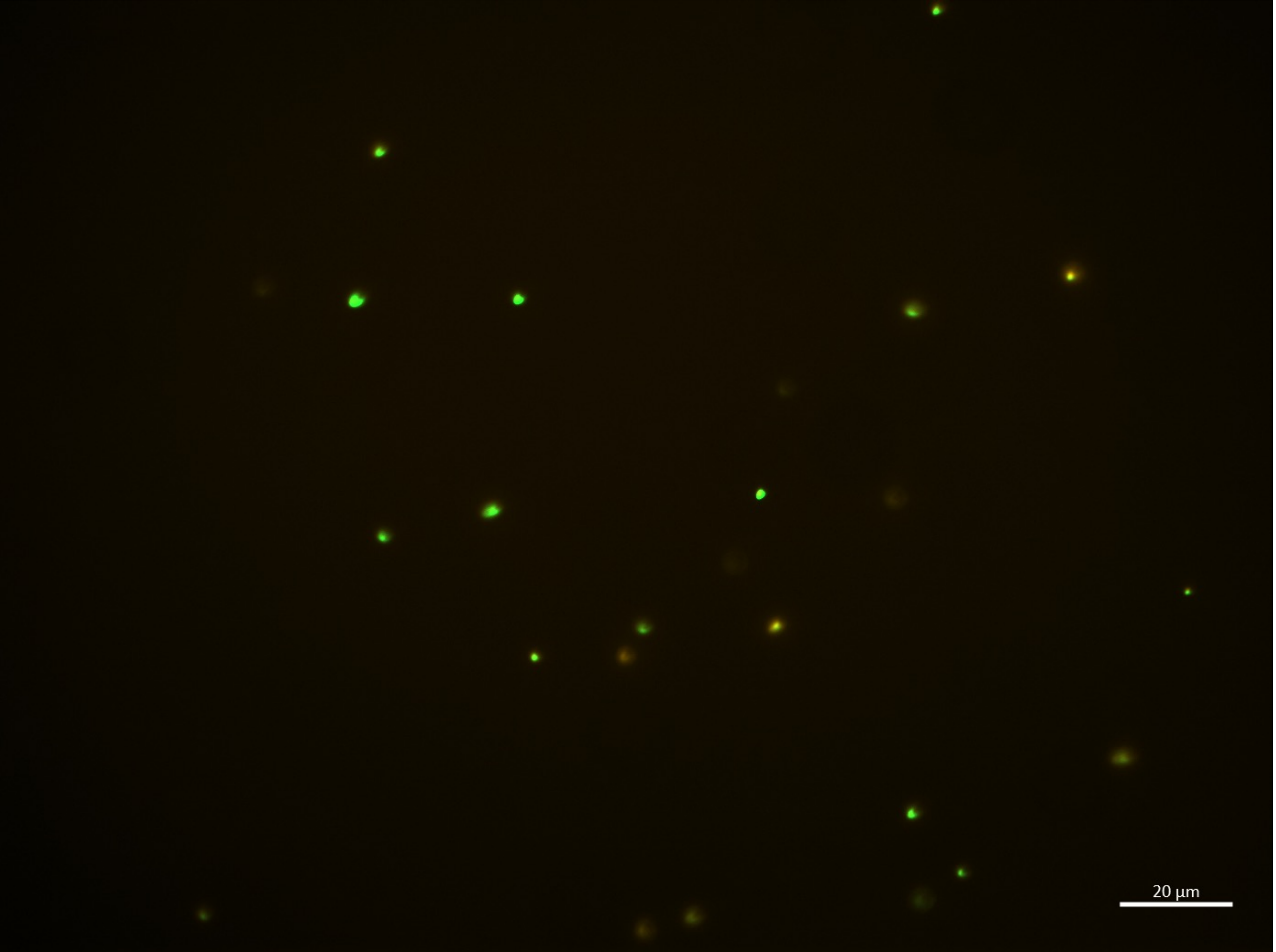

16:00-F

| | |
|---|---|
| B.bifidum | **7** |
| E.coli | 17 |
| **Sum** | 24 |

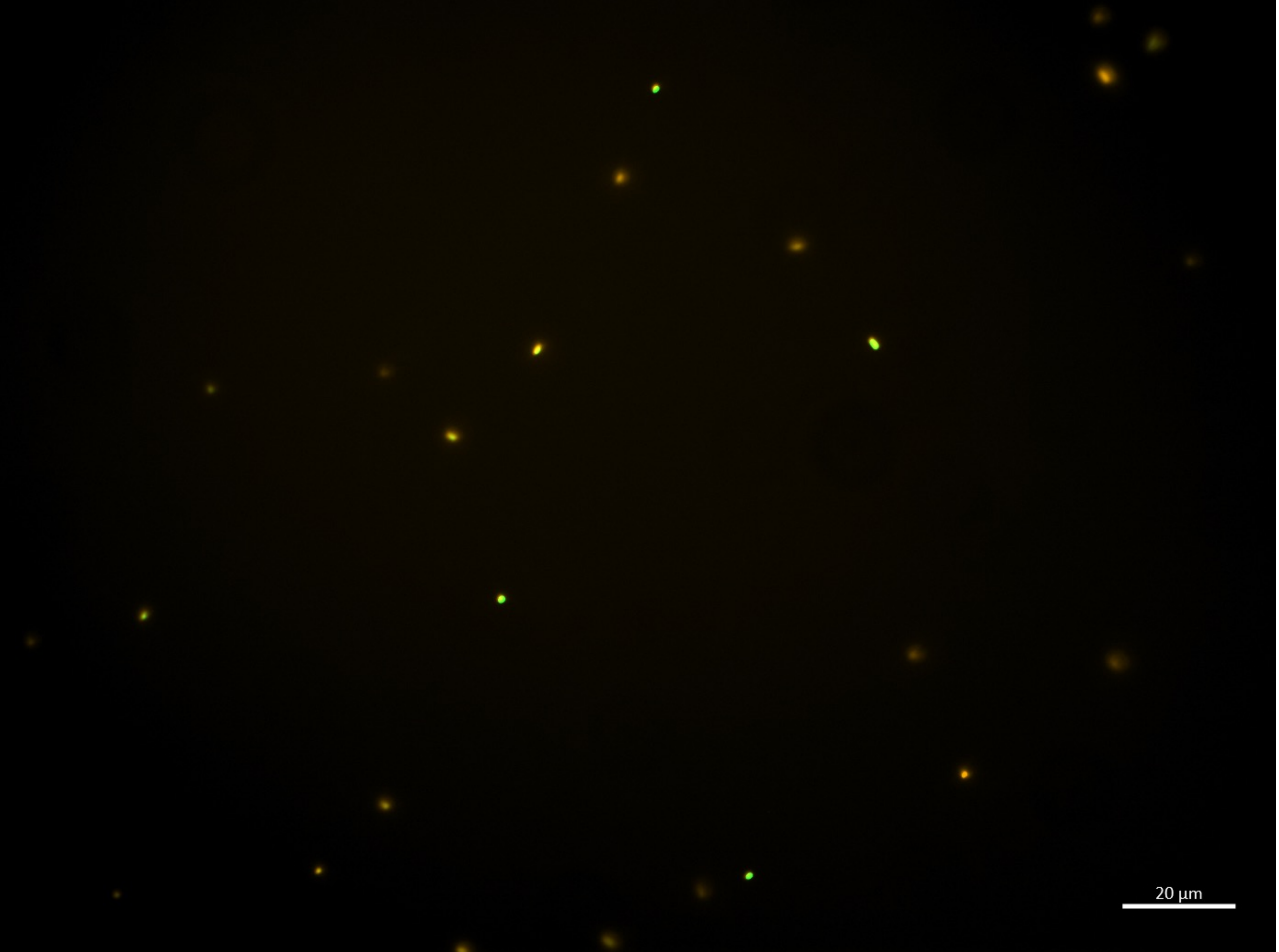

16:00-G

| | |
|---|---|
| **B.bifidum** | **15** |
| **E.coli** | 8 |
| **Sum** | 23 |

16:00-H

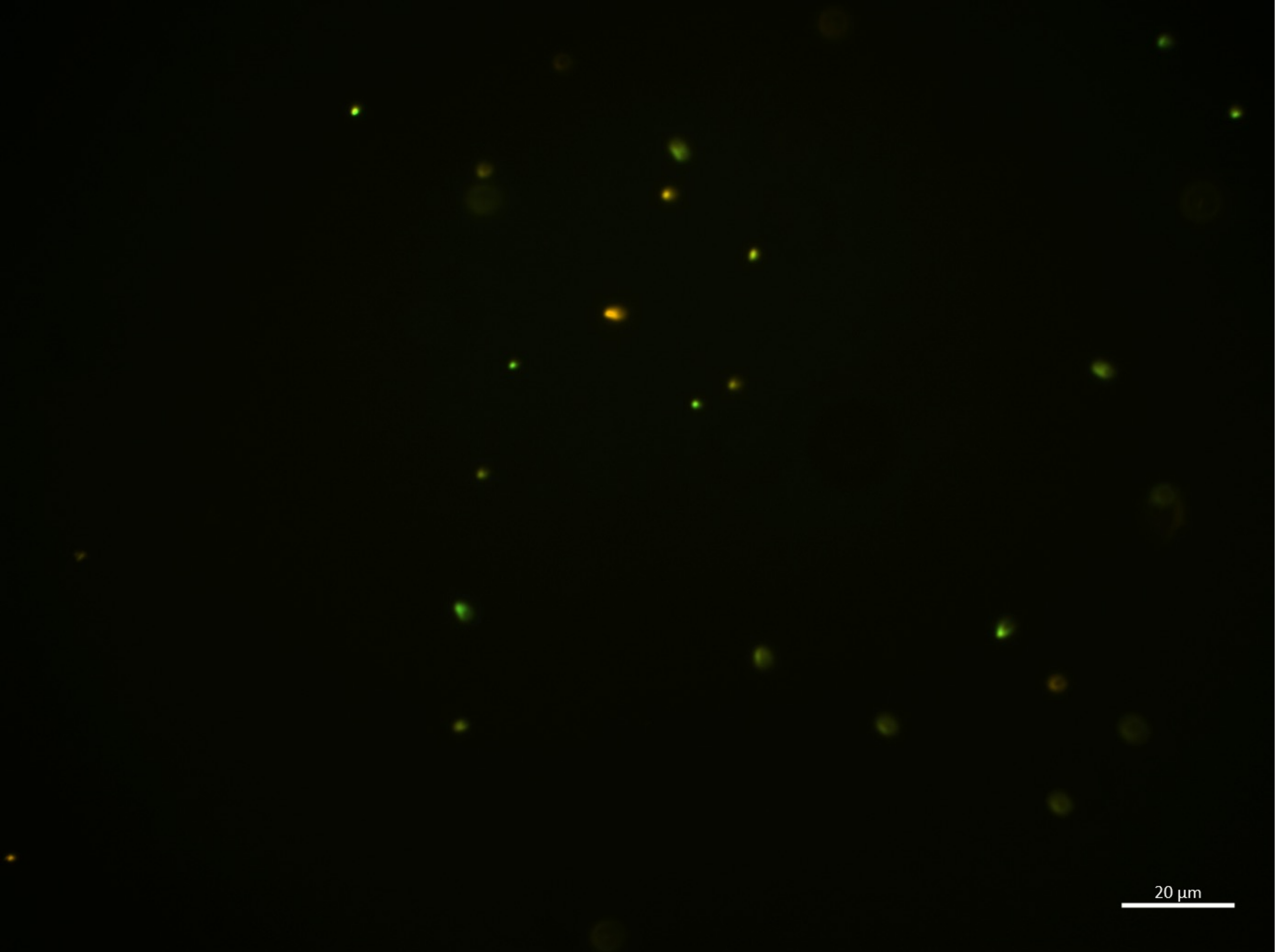

| | |
|---|---|
| B.bifidum | **4** |
| E.coli | 17 |
| **Sum** | 21 |

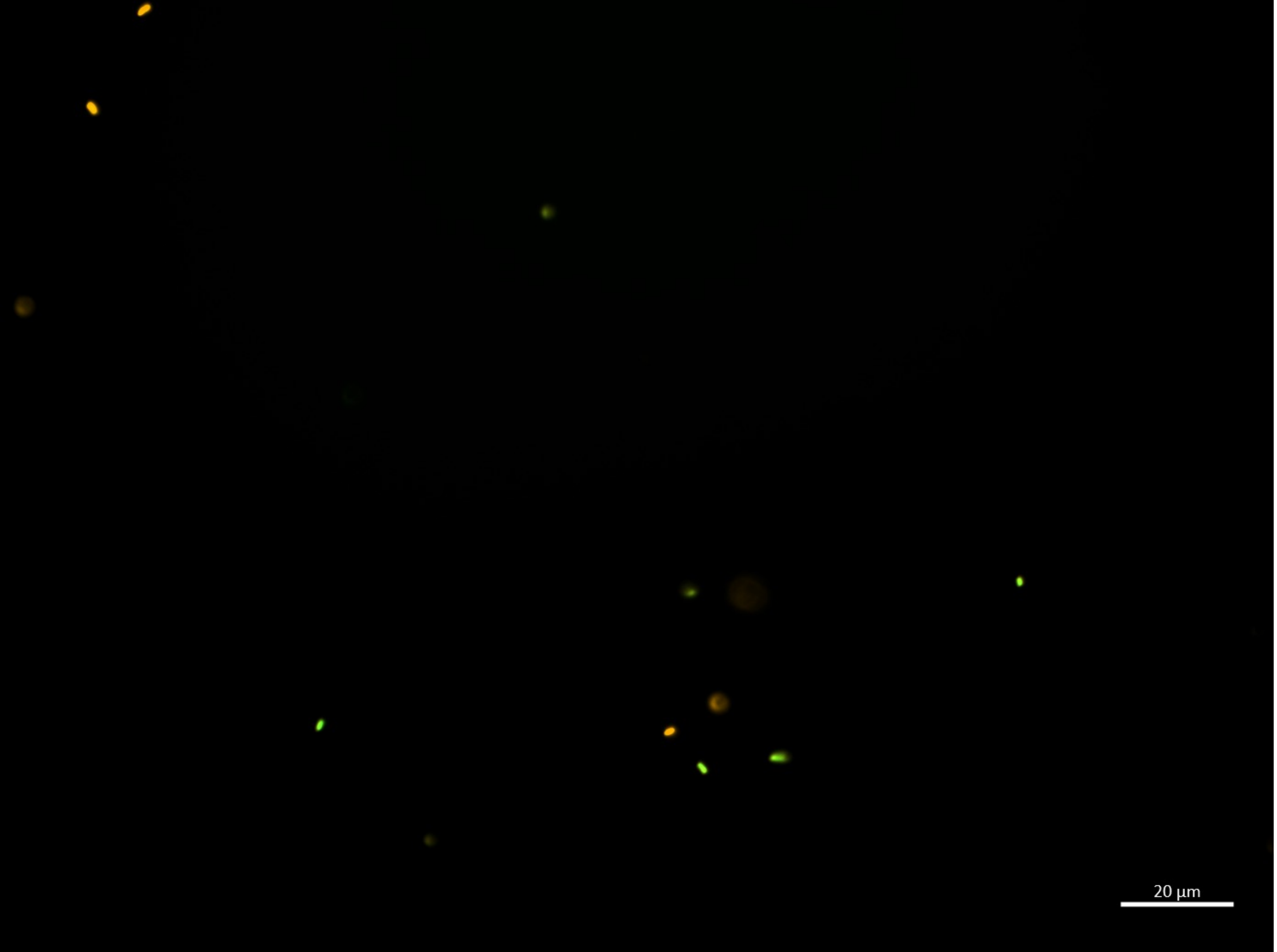

16:00-I

| | |
|---|---|
| B.bifidum | 4 |
| E.coli | 7 |
| Sum | 11 |

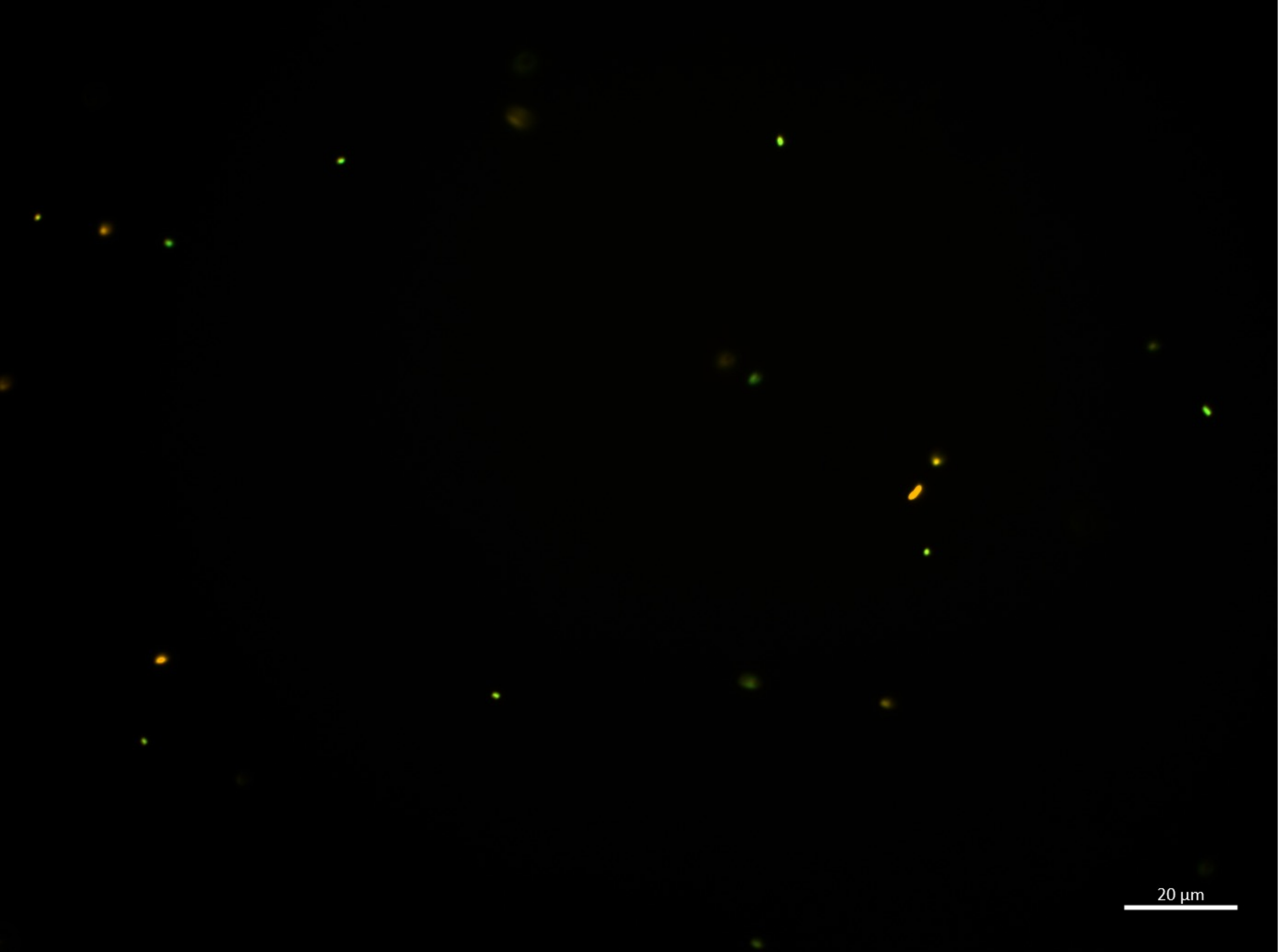

16:00-J

| | |
|---|---|
| **B.bifidum** | **6** |
| **E.coli** | 11 |
| **Sum** | 17 |

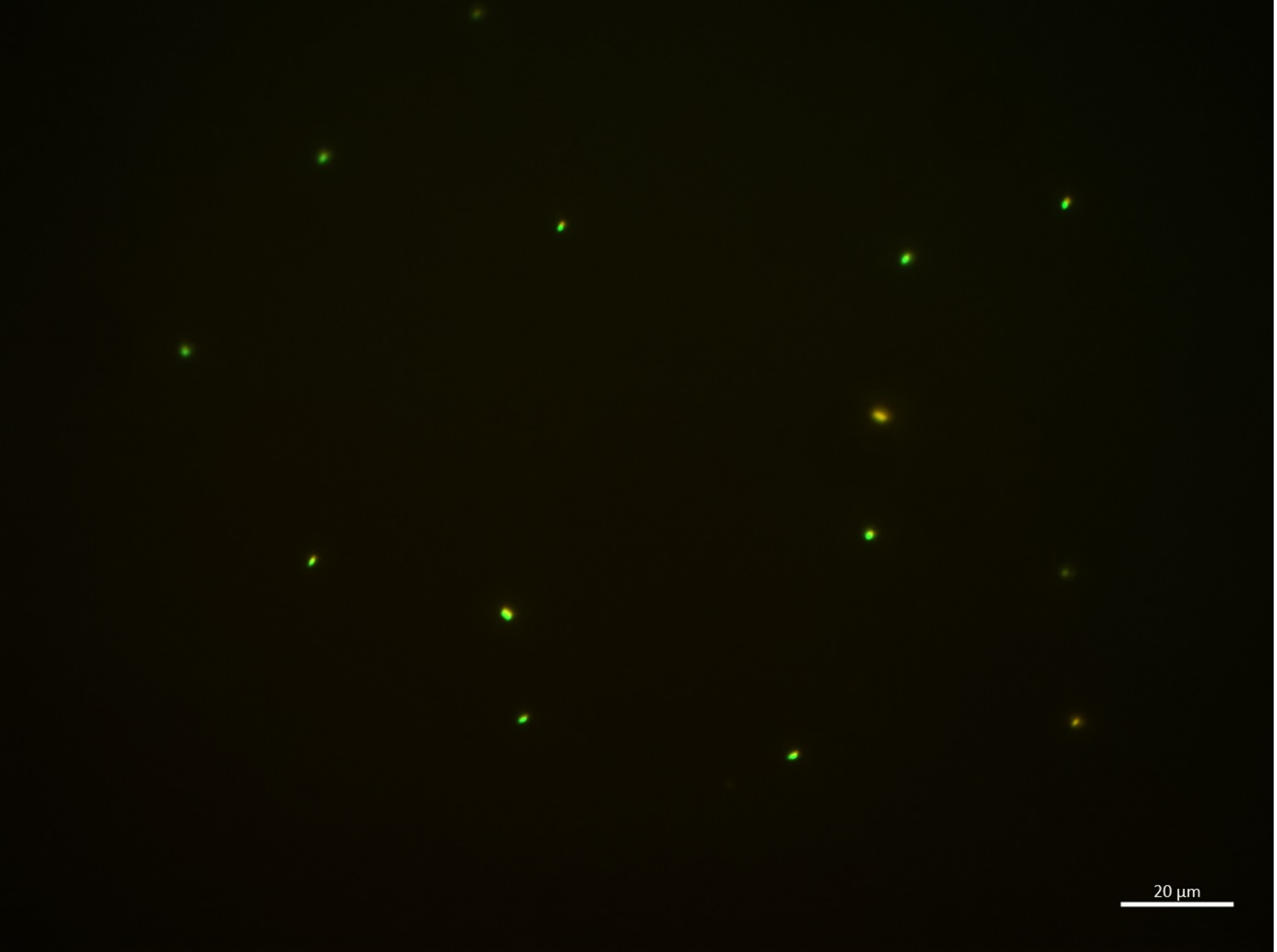

18:00-A

| | |
|---|---|
| **B.bifidum** | **2** |
| **E.coli** | 11 |
| **Sum** | 13 |

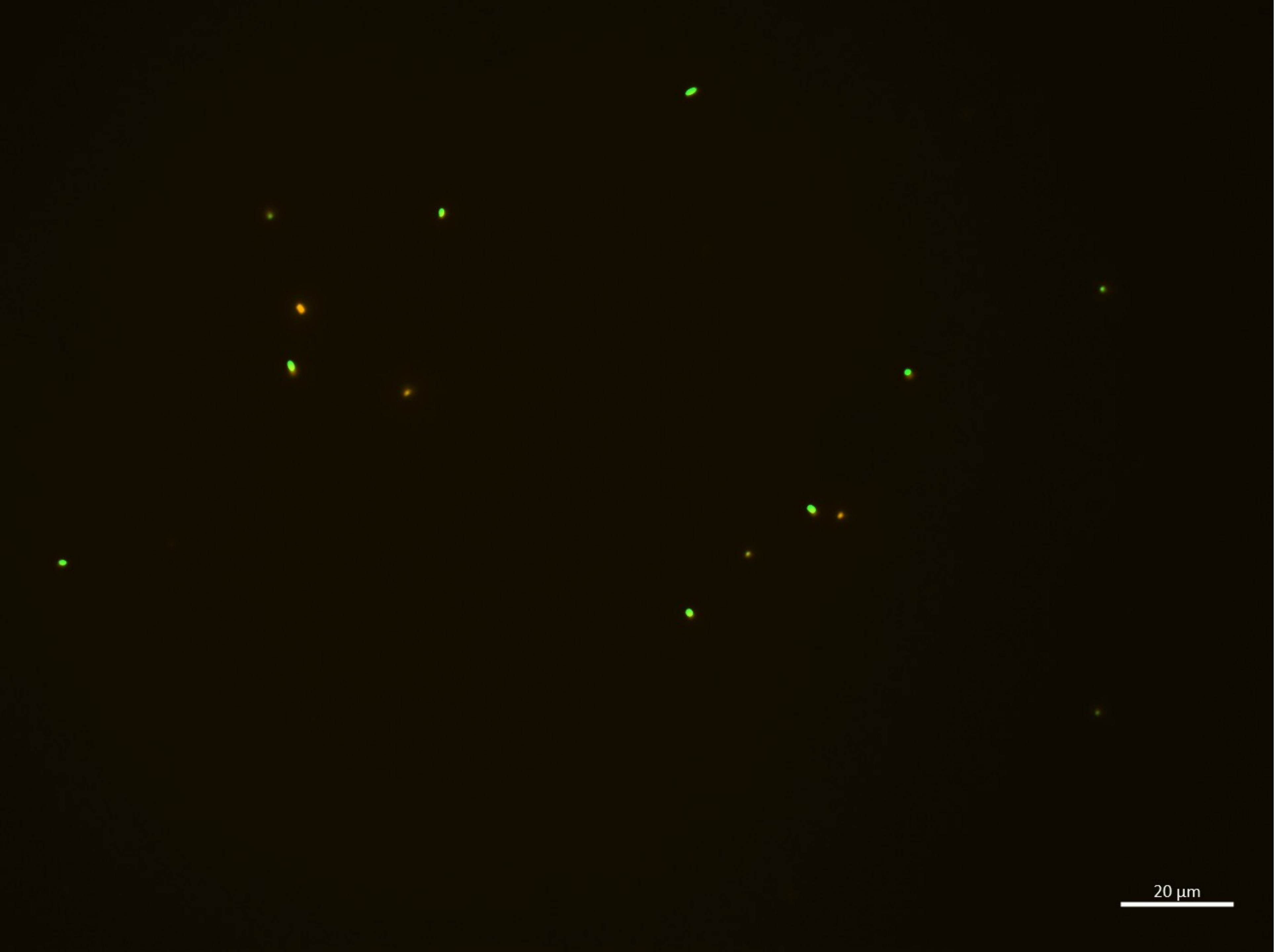

18:00-B

| | |
|---|---|
| **B.bifidum** | **3** |
| **E.coli** | 11 |
| **Sum** | 14 |

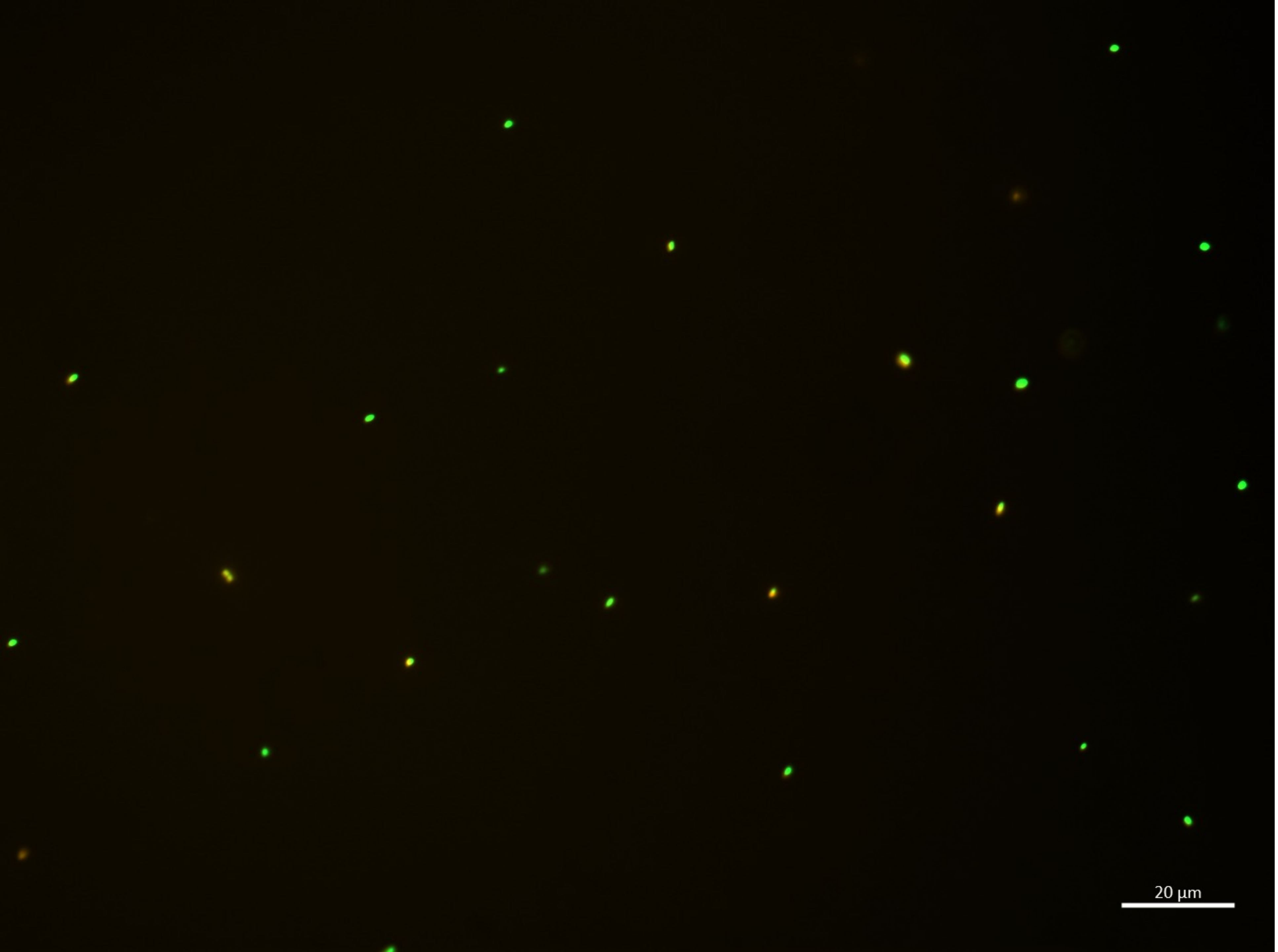

18:00-C

| | |
|---|---|
| B.bifidum | **2** |
| E.coli | 23 |
| **Sum** | 25 |

18:00-D

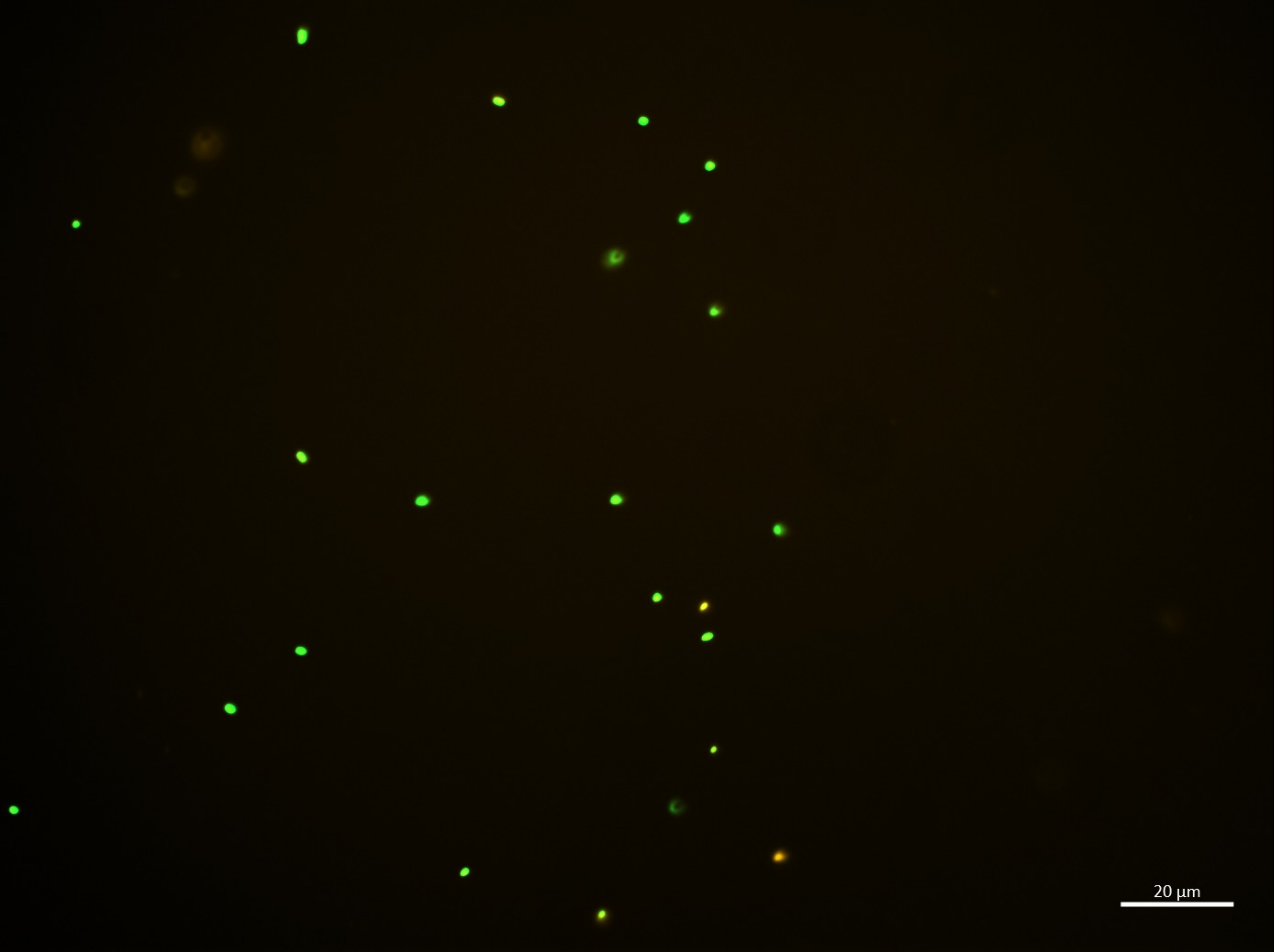

| | |
|---|---|
| **B.bifidum** | **3** |
| **E.coli** | 20 |
| **Sum** | 23 |

18:00-E

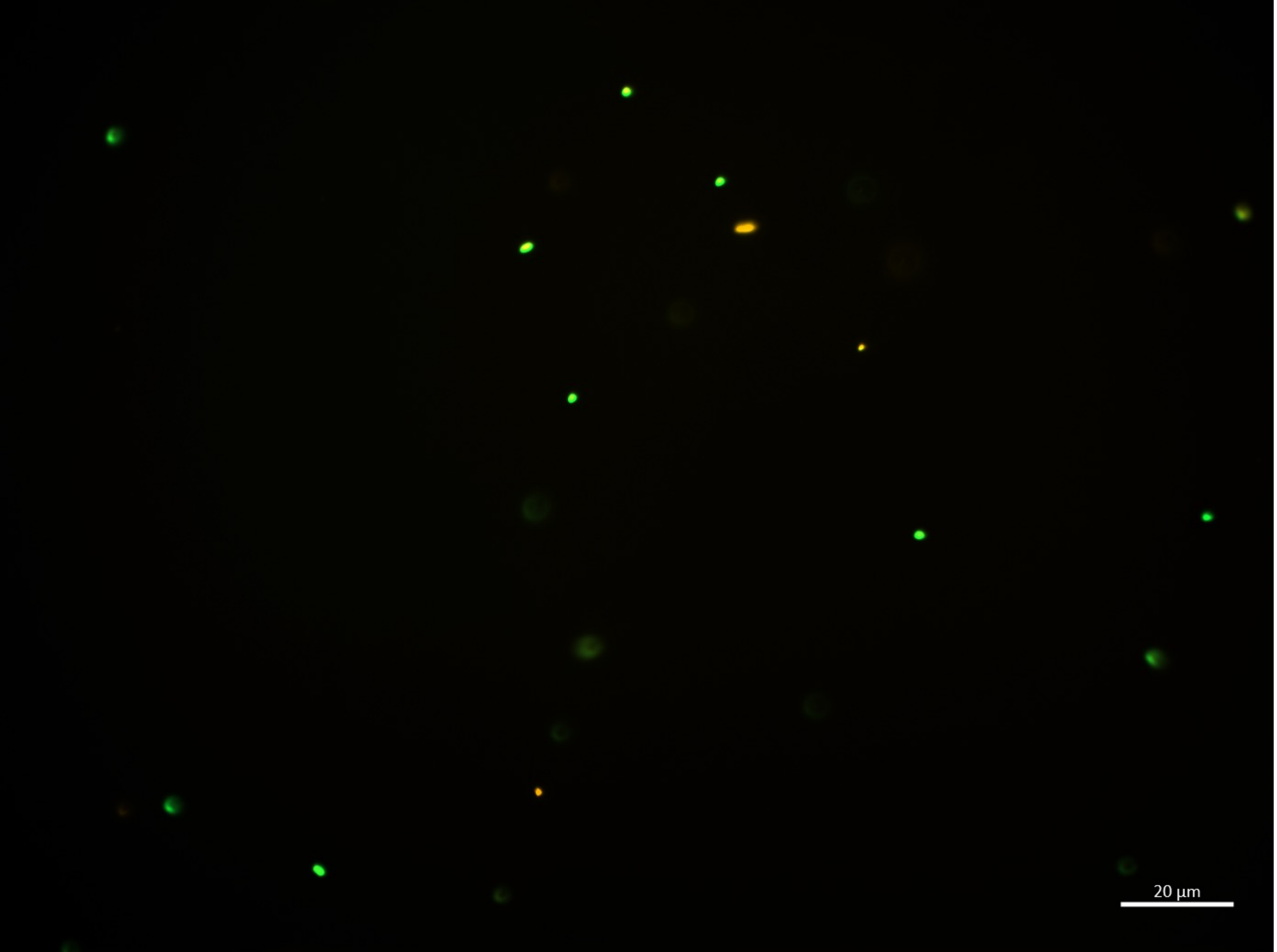

| | |
|---|---|
| B.bifidum | **2** |
| E.coli | 12 |
| **Sum** | 14 |

18:00-F

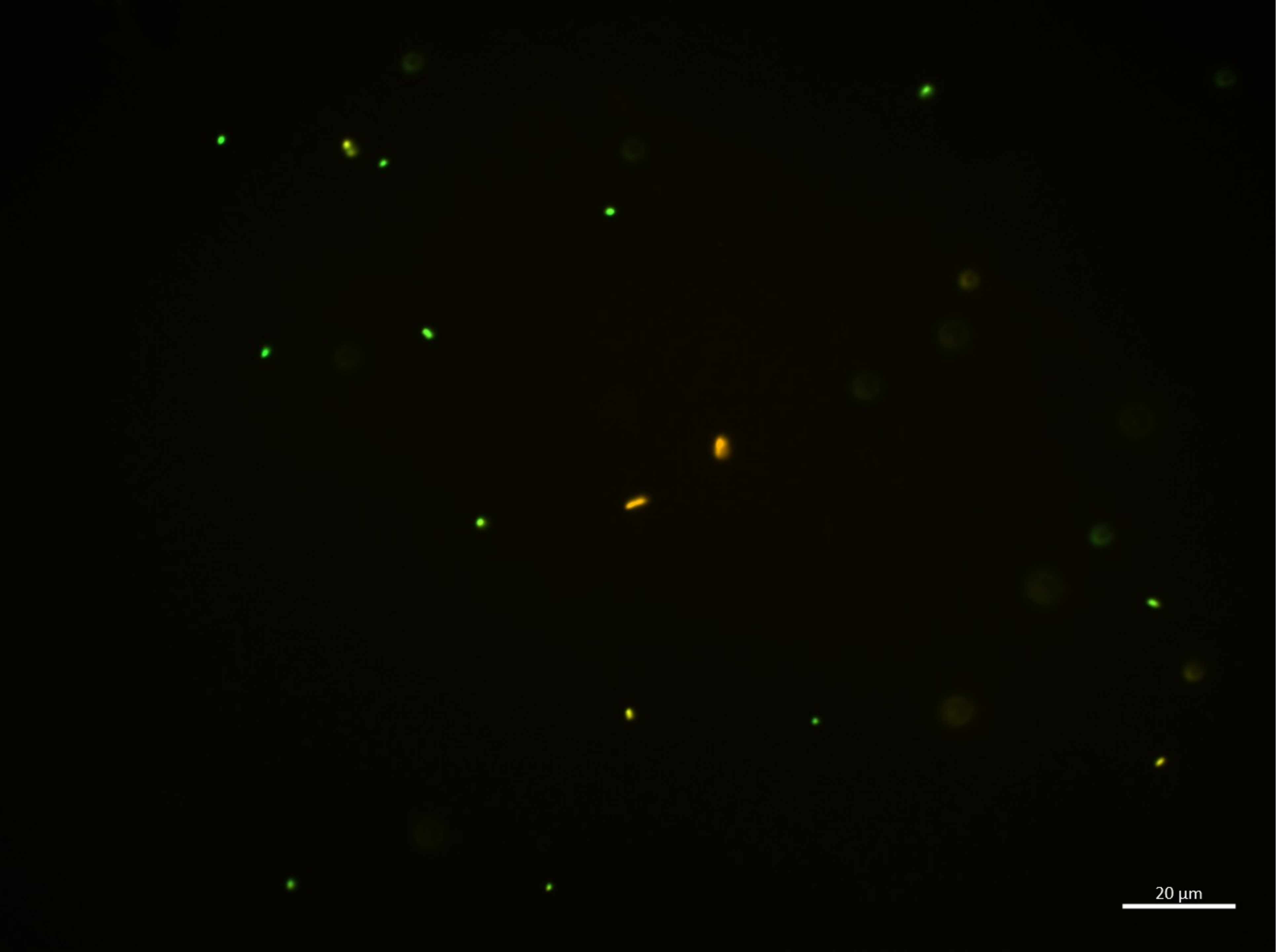

| | |
|---|---|
| B.bifidum | 3 |
| E.coli | 12 |
| Sum | 15 |

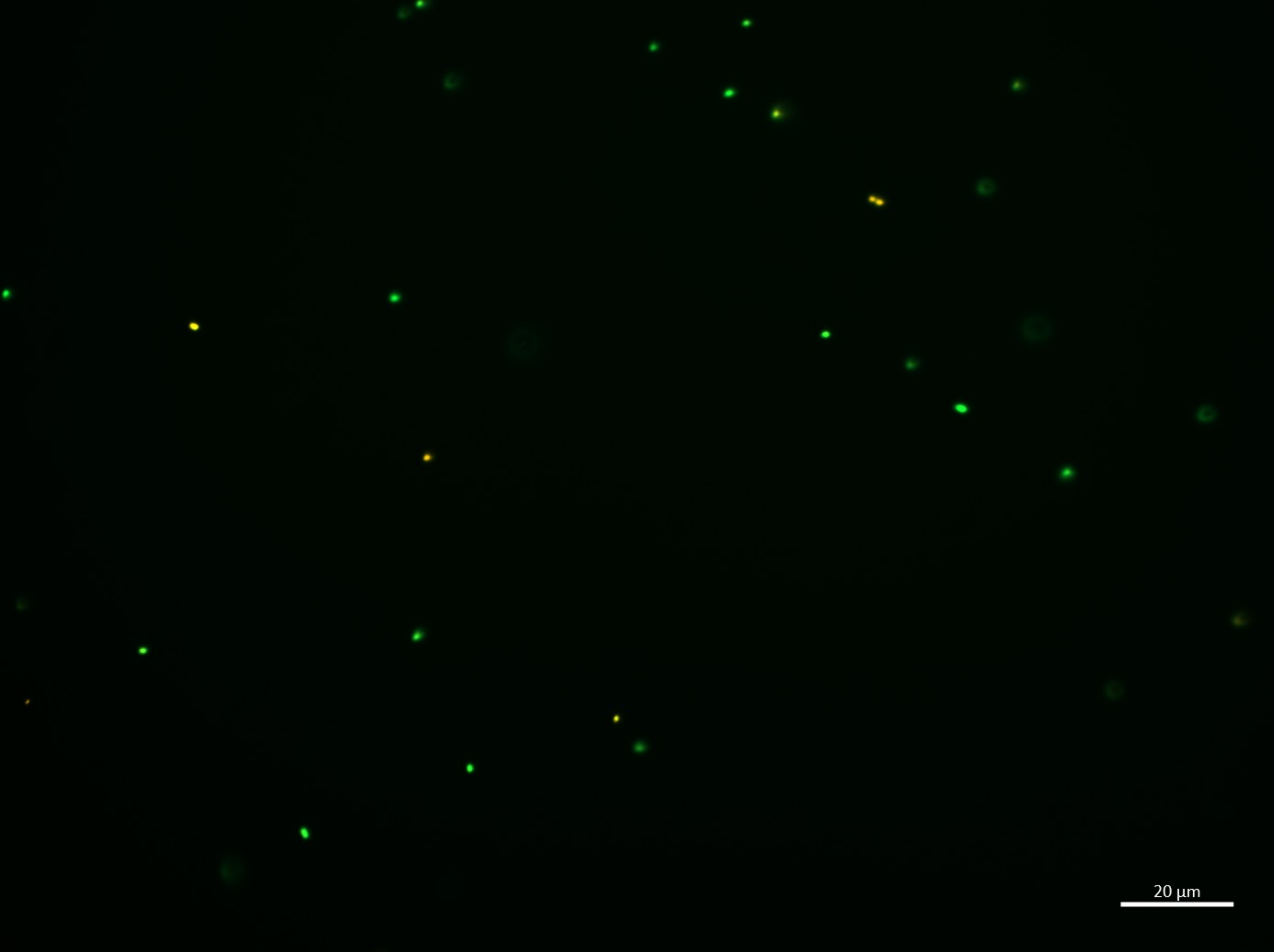

18:00-G

| | |
|---|---|
| **B.bifidum** | **3** |
| **E.coli** | 22 |
| **Sum** | 25 |

18:00-H

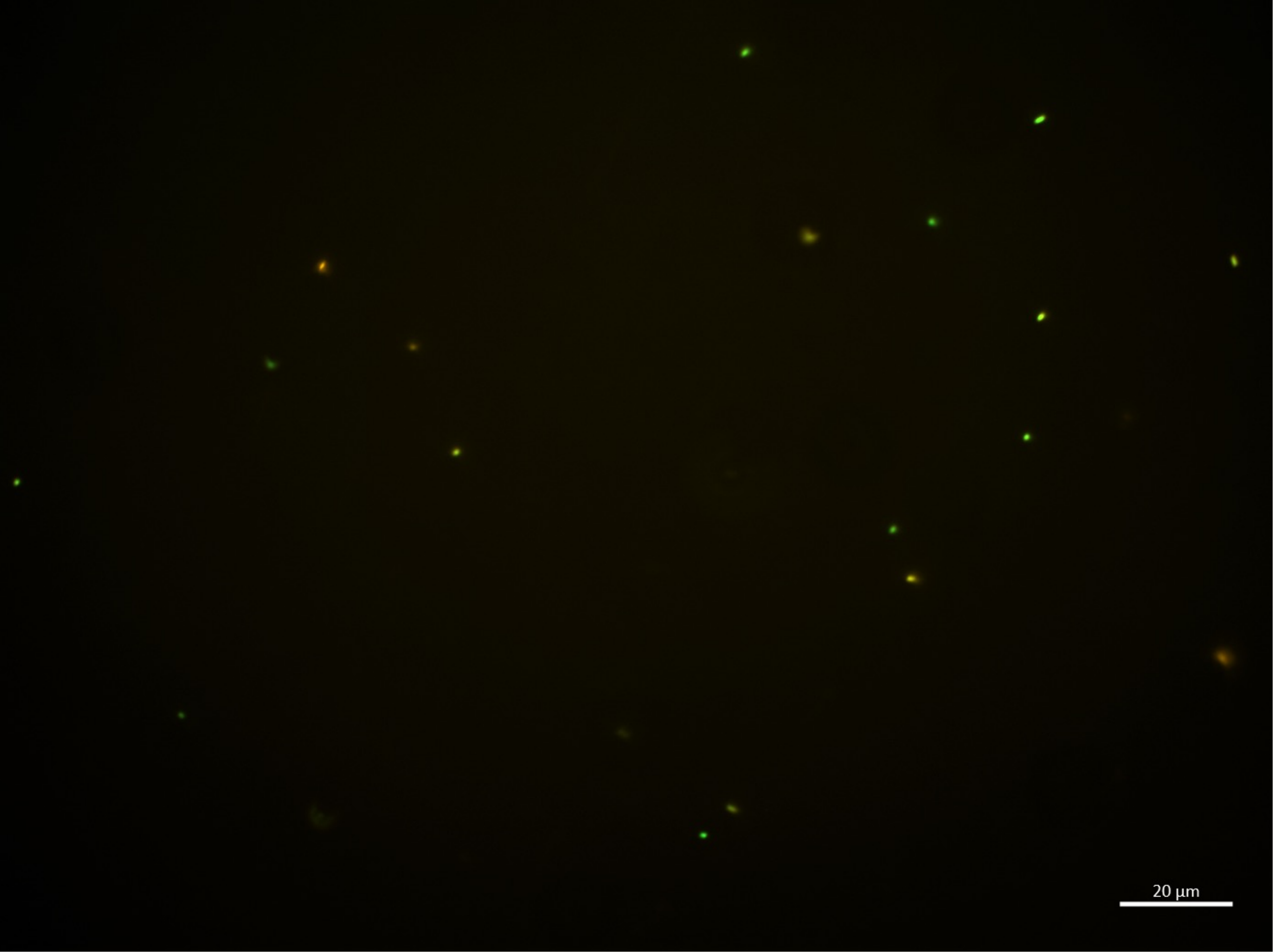

| | |
|---|---|
| **B.bifidum** | **5** |
| **E.coli** | 14 |
| **Sum** | 19 |

18:00-I

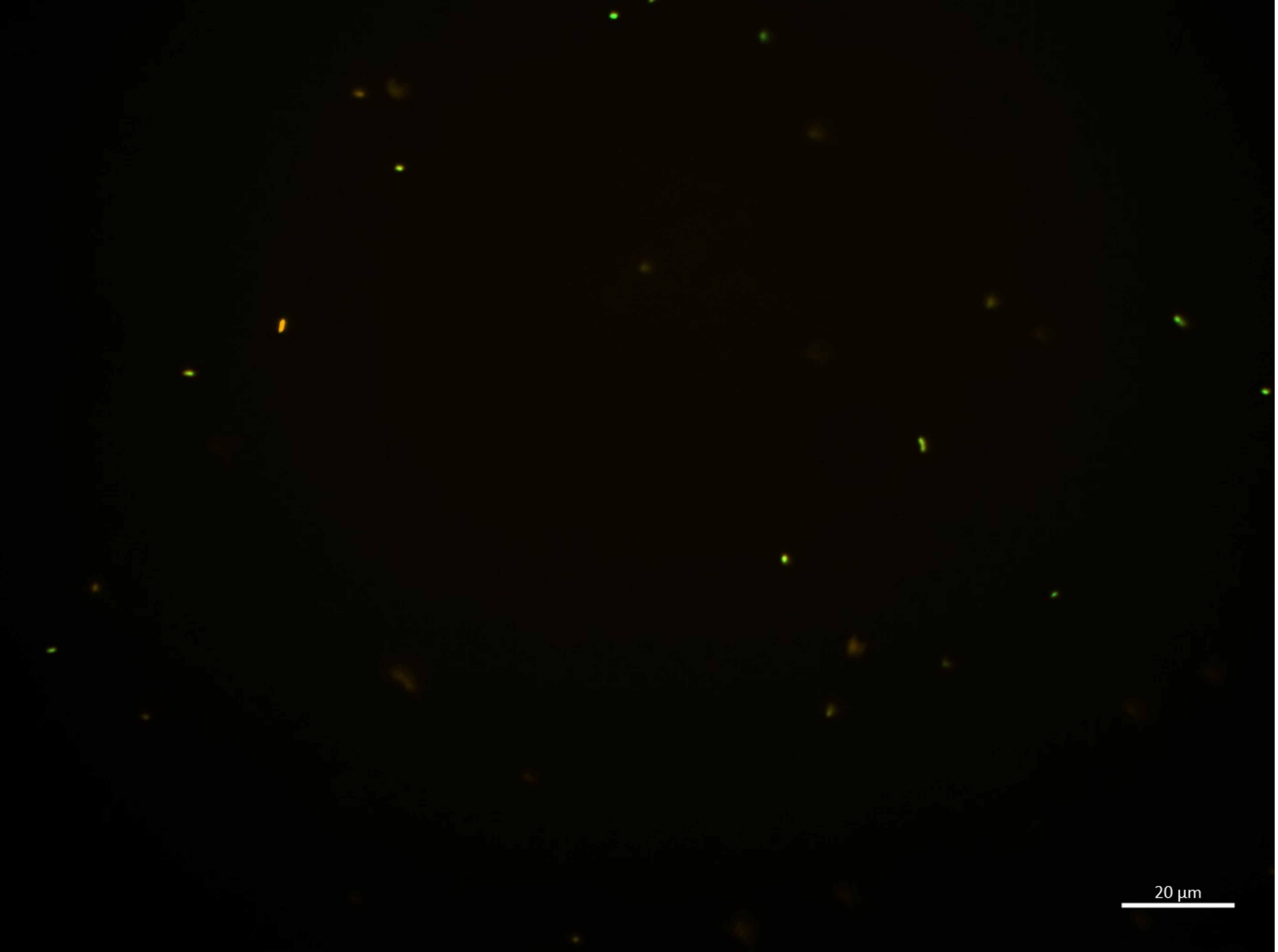

| | |
|---|---|
| **B.bifidum** | **8** |
| **E.coli** | 12 |
| **Sum** | 20 |

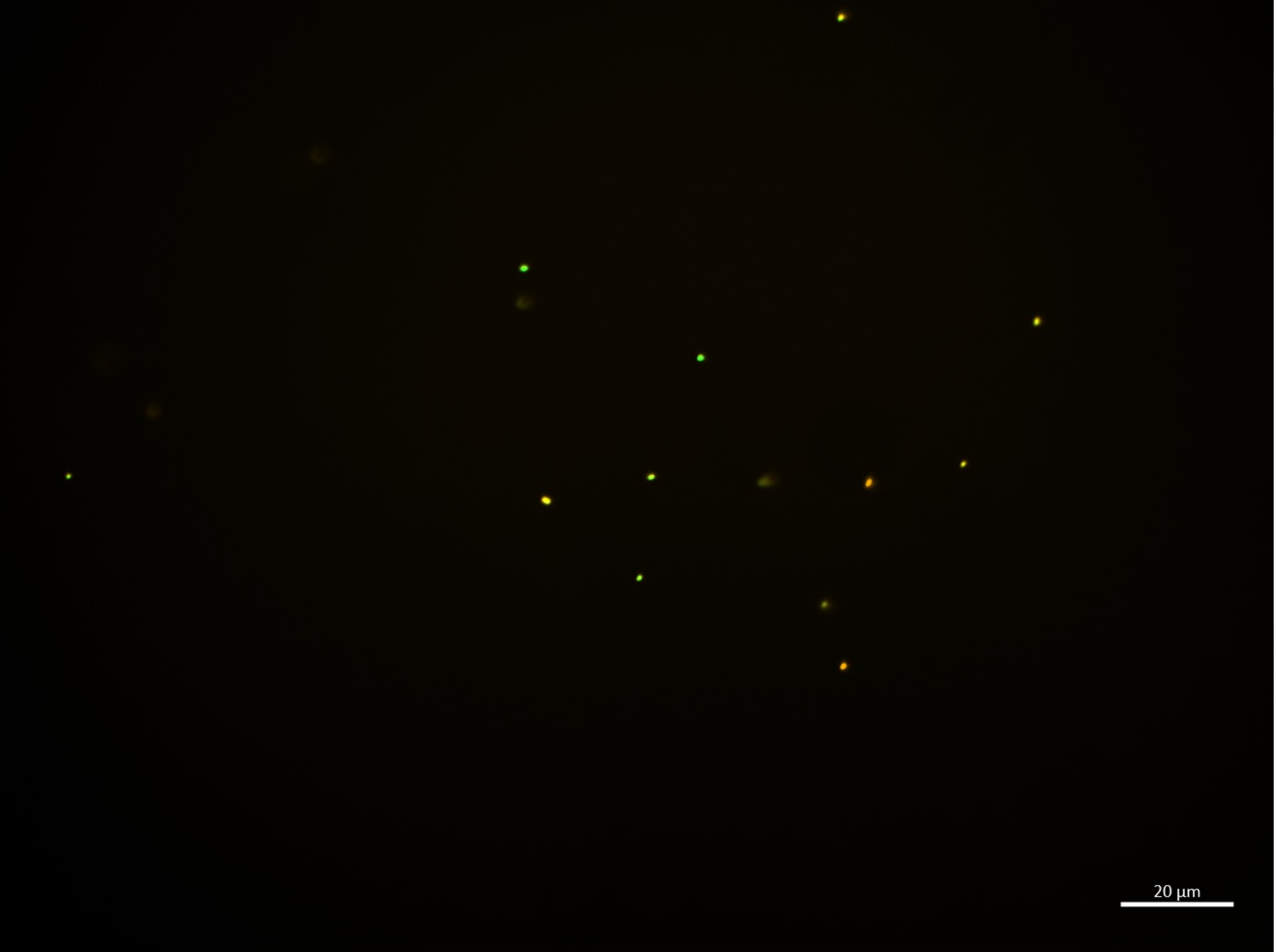

18:00-J

| | |
|---|---|
| B.bifidum | **5** |
| E.coli | 9 |
| **Sum** | 14 |

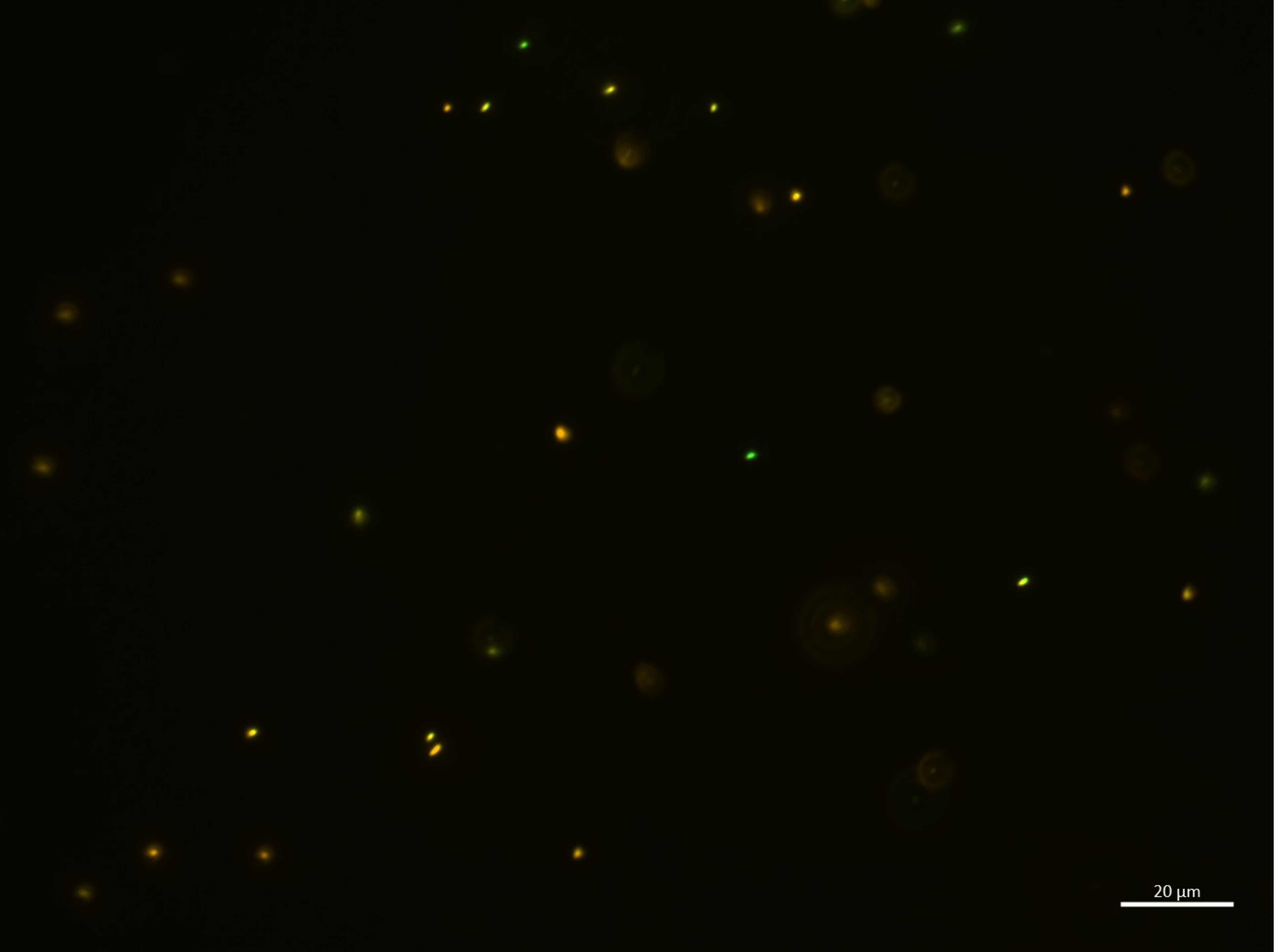

20:00-A

| | |
|---|---|
| B.bifidum | **33** |
| E.coli | 4 |
| **Sum** | 37 |

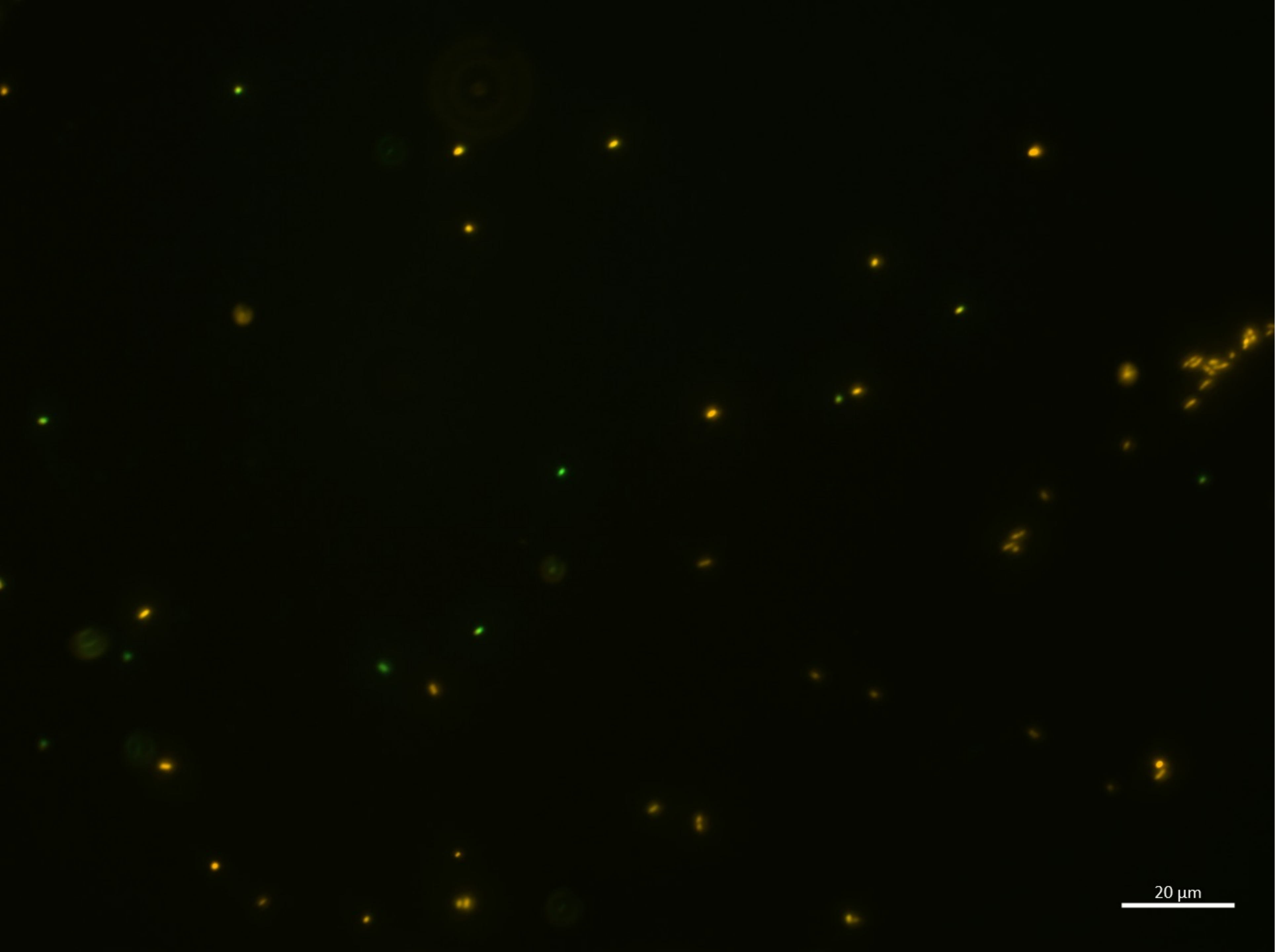

20:00-B

| | |
|---|---|
| B.bifidum | **51** |
| E.coli | 6 |
| **Sum** | 57 |

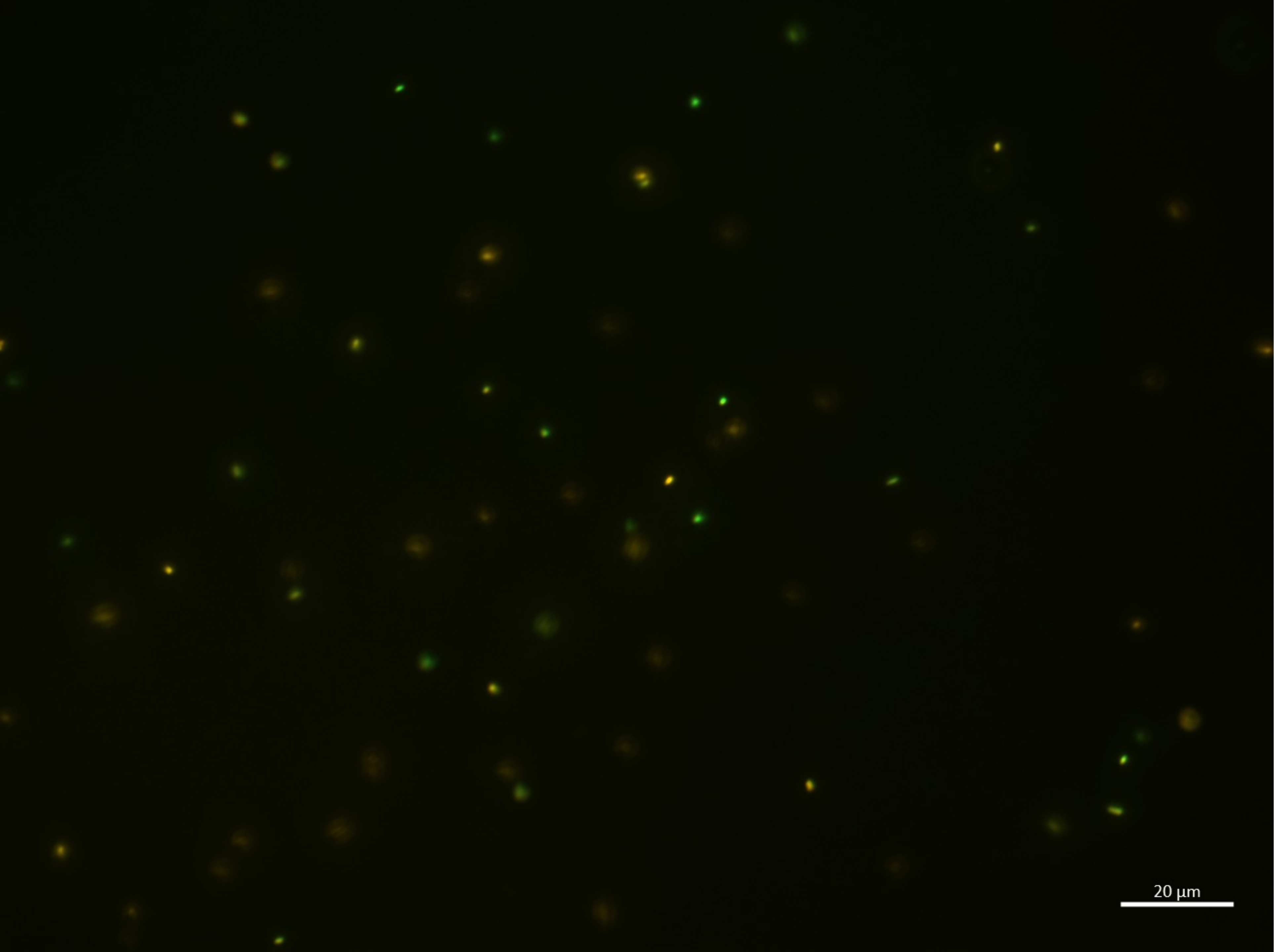

20:00-C

| | |
|---|---|
| B.bifidum | **55** |
| E.coli | 13 |
| **Sum** | 68 |

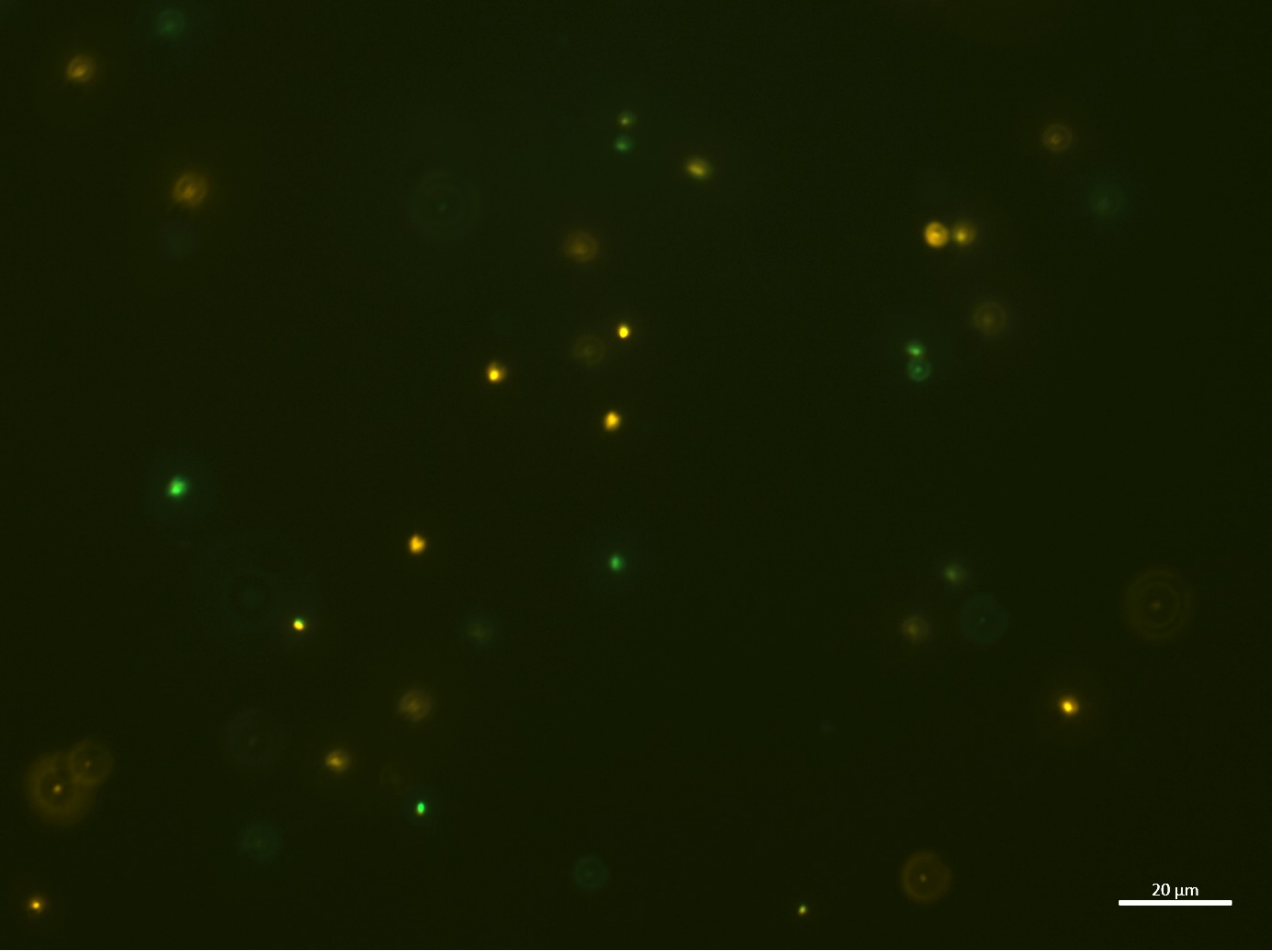

20:00-D

| | |
|---|---|
| **B.bifidum** | **20** |
| **E.coli** | 6 |
| **Sum** | 26 |

20:00-E

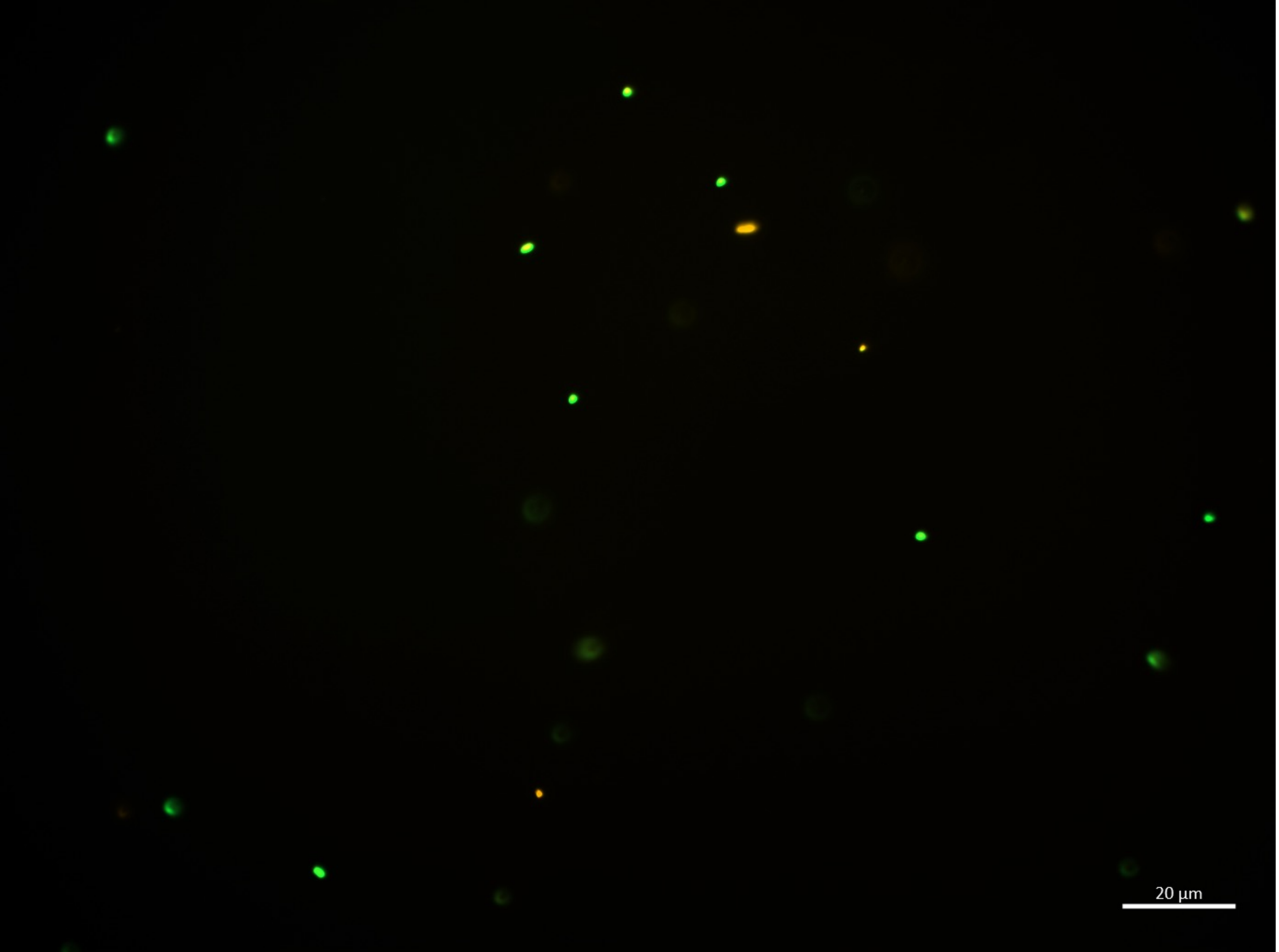

| | |
|---|---|
| **B.bifidum** | **7** |
| **E.coli** | 10 |
| **Sum** | 17 |

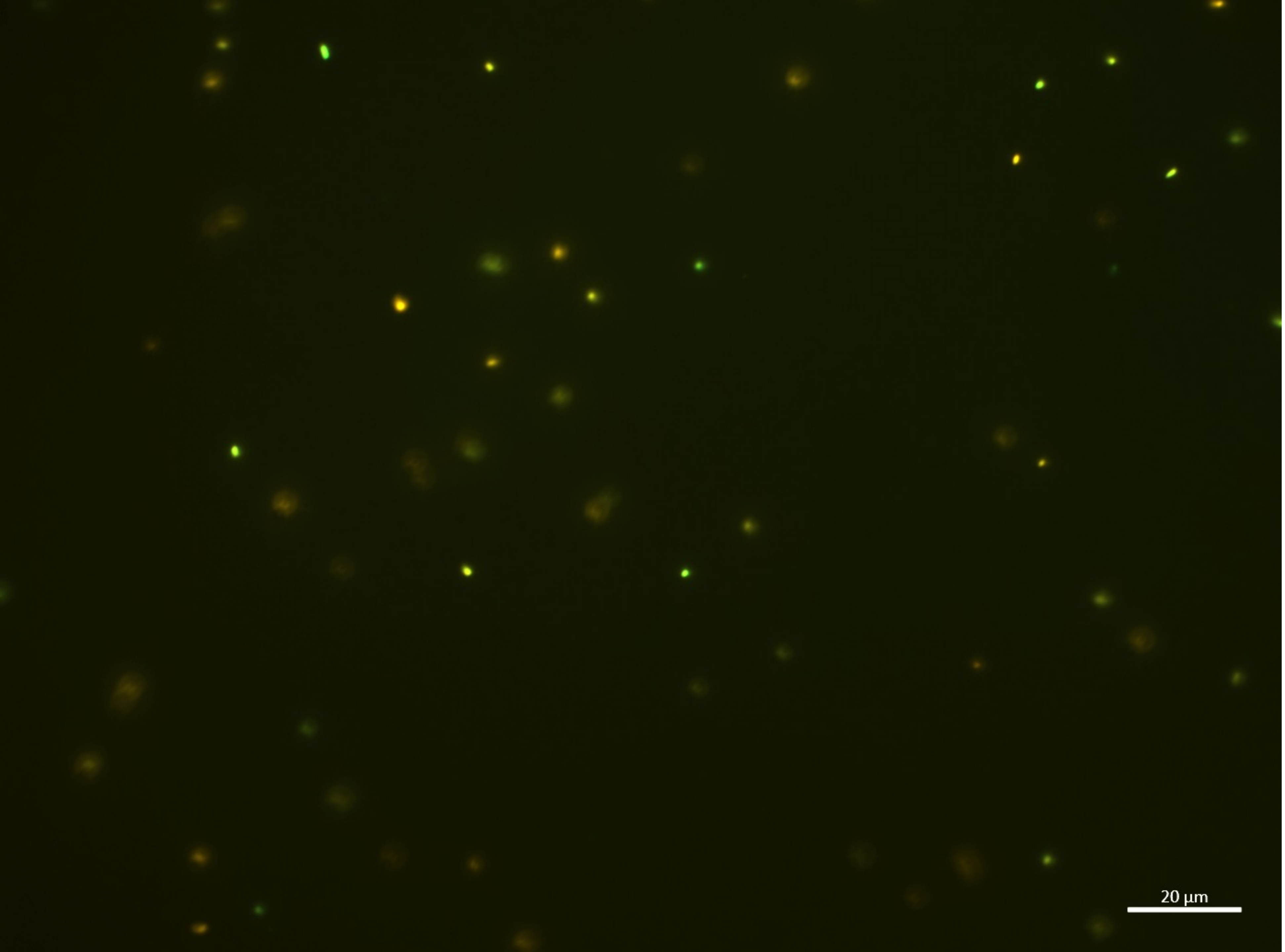

20:00-F

| | |
|---|---|
| **B.bifidum** | **36** |
| **E.coli** | 10 |
| **Sum** | 46 |

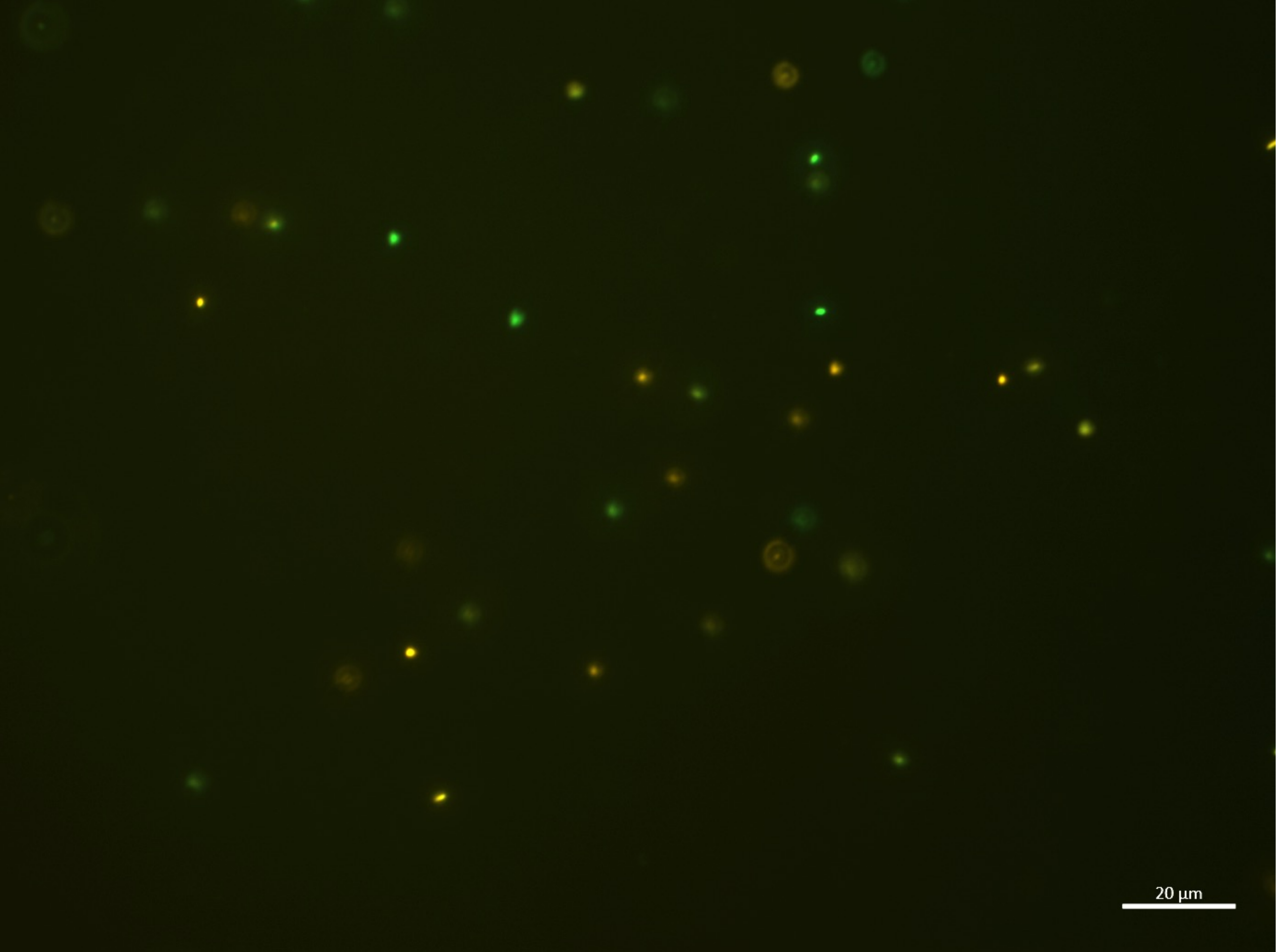

20:00-G

| | |
|---|---|
| B.bifidum | **22** |
| E.coli | 12 |
| **Sum** | 34 |

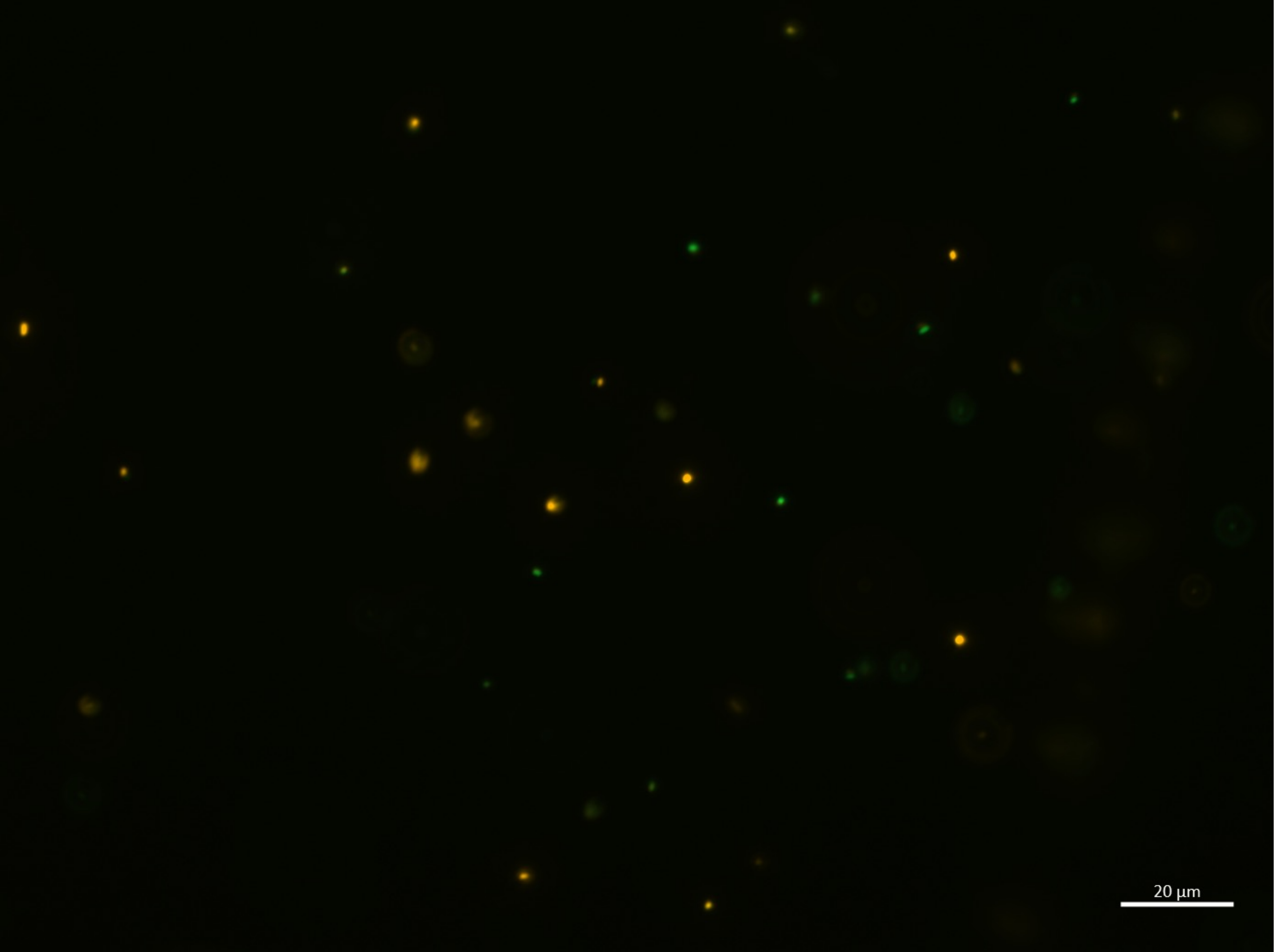

20:00-H

| | |
|---|---|
| **B.bifidum** | **25** |
| **E.coli** | 10 |
| **Sum** | 35 |

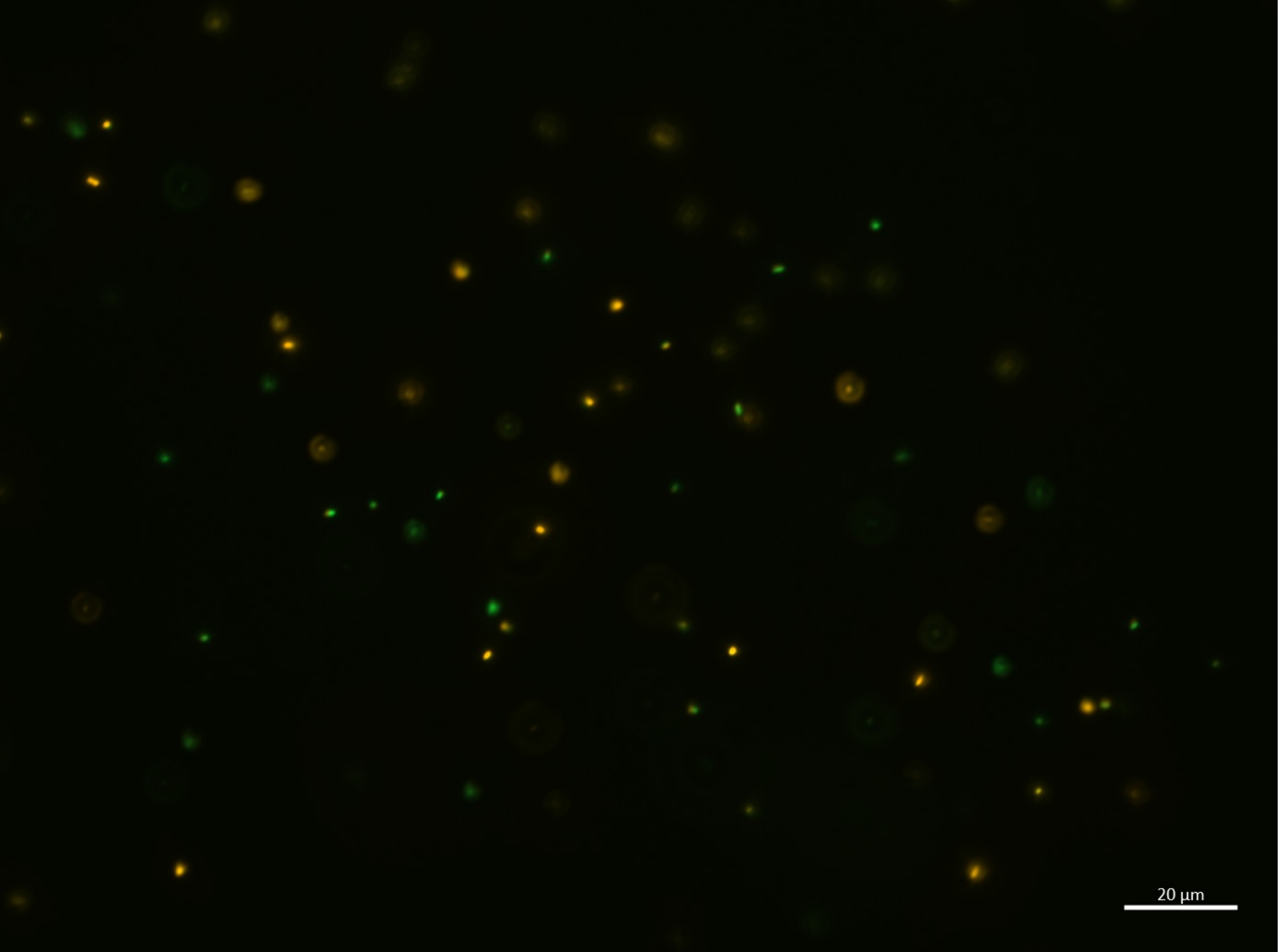

20:00-I

| | |
|---|---|
| B.bifidum | **50** |
| E.coli | 18 |
| **Sum** | 64 |

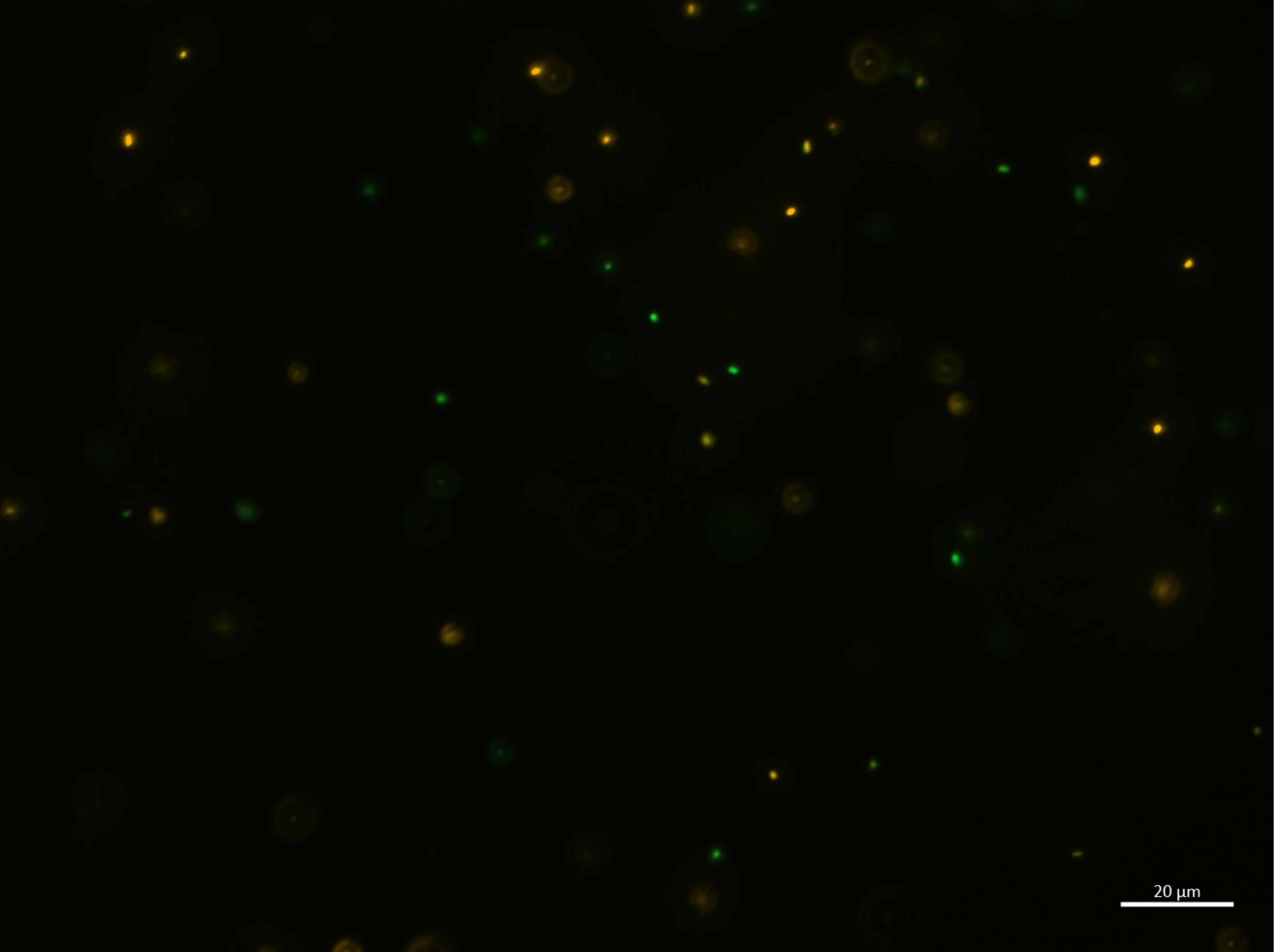

20:00-J

| | |
|---|---|
| B.bifidum | **38** |
| E.coli | 13 |
| **Sum** | 51 |

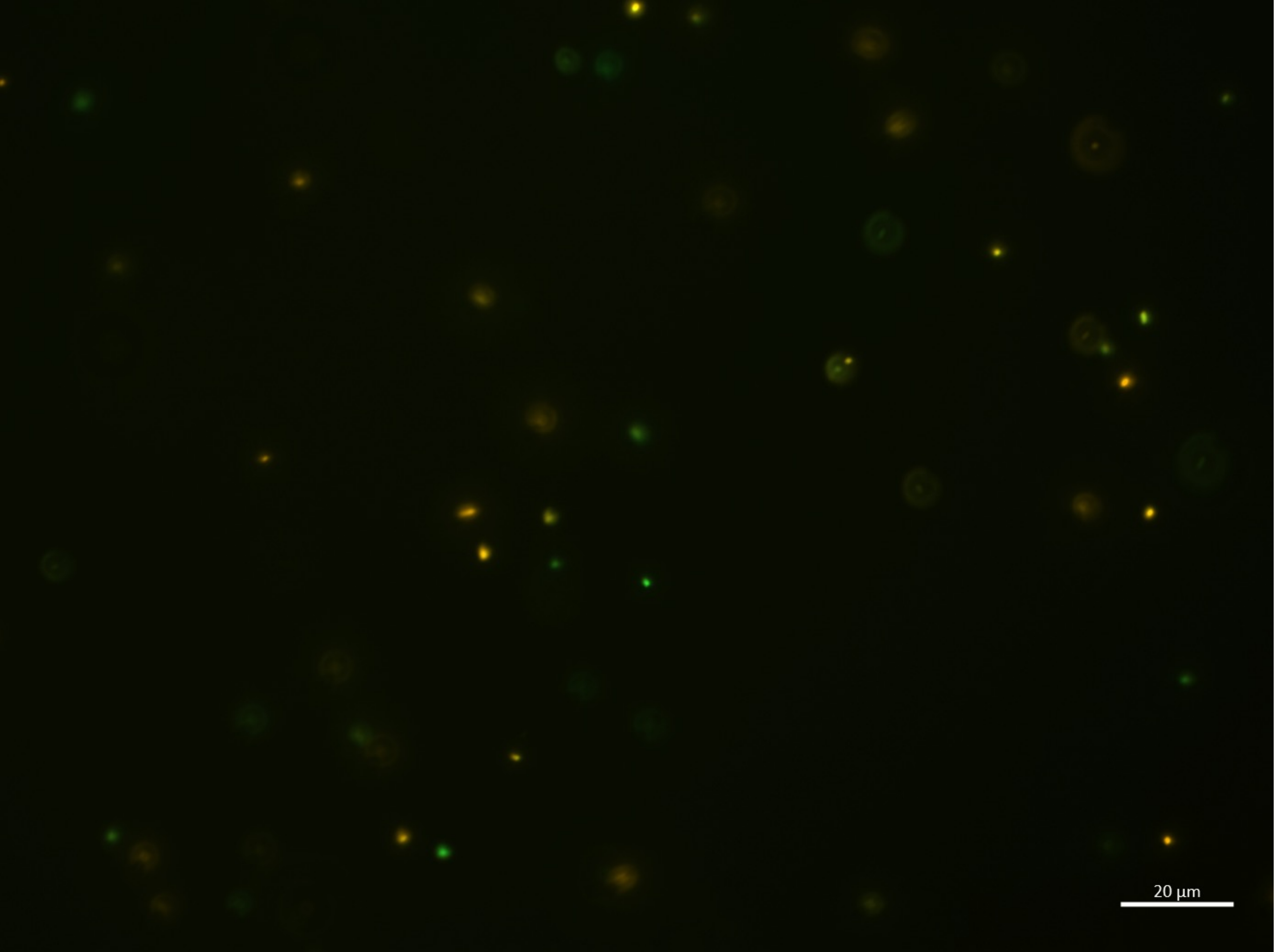

20:00-K

| | |
|---|---|
| **B.bifidum** | **36** |
| E.coli | 12 |
| **Sum** | 48 |

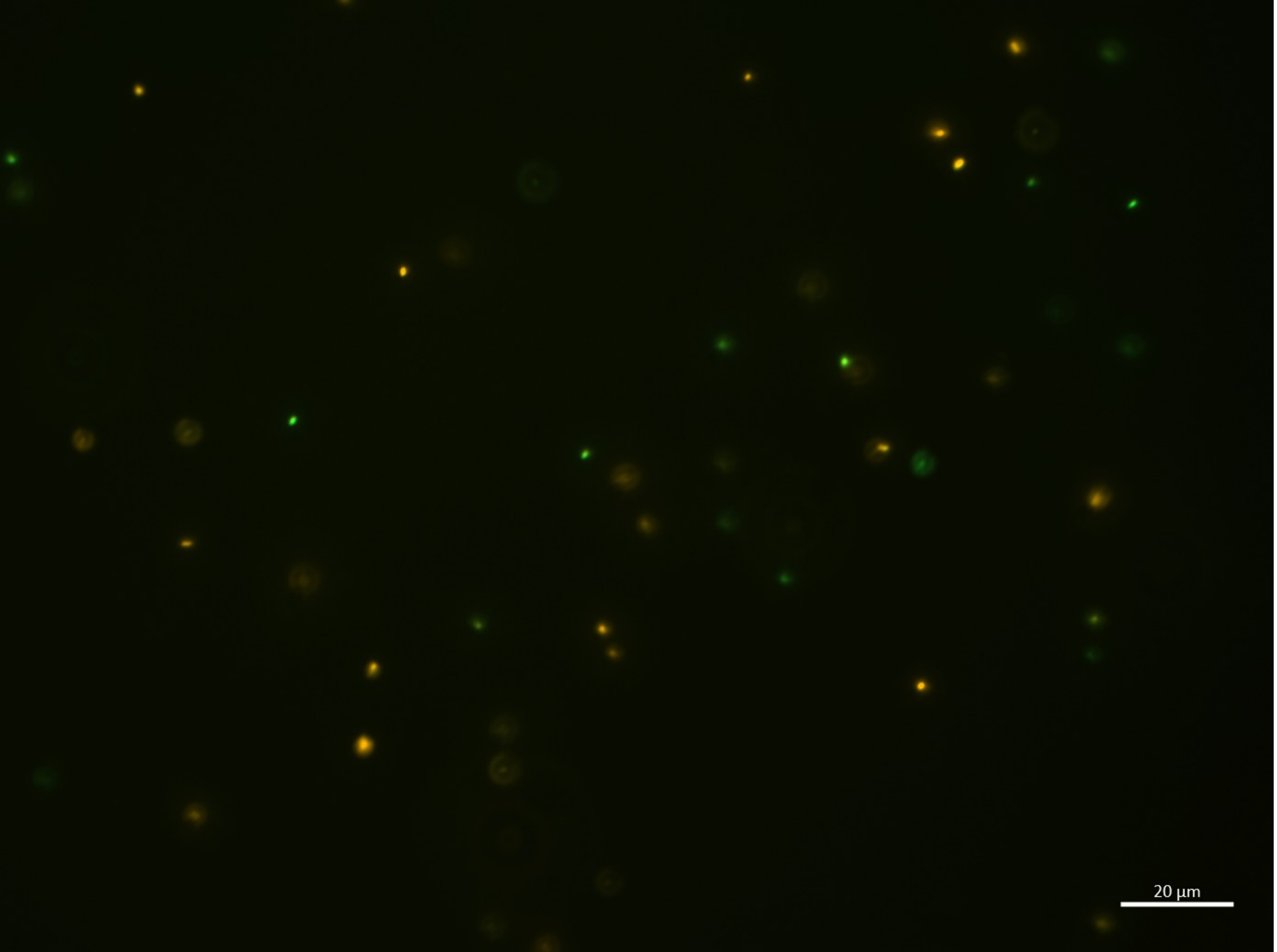

20:00-L

| | |
|---|---|
| B.bifidum | **33** |
| E.coli | 13 |
| **Sum** | 46 |

20:00-M

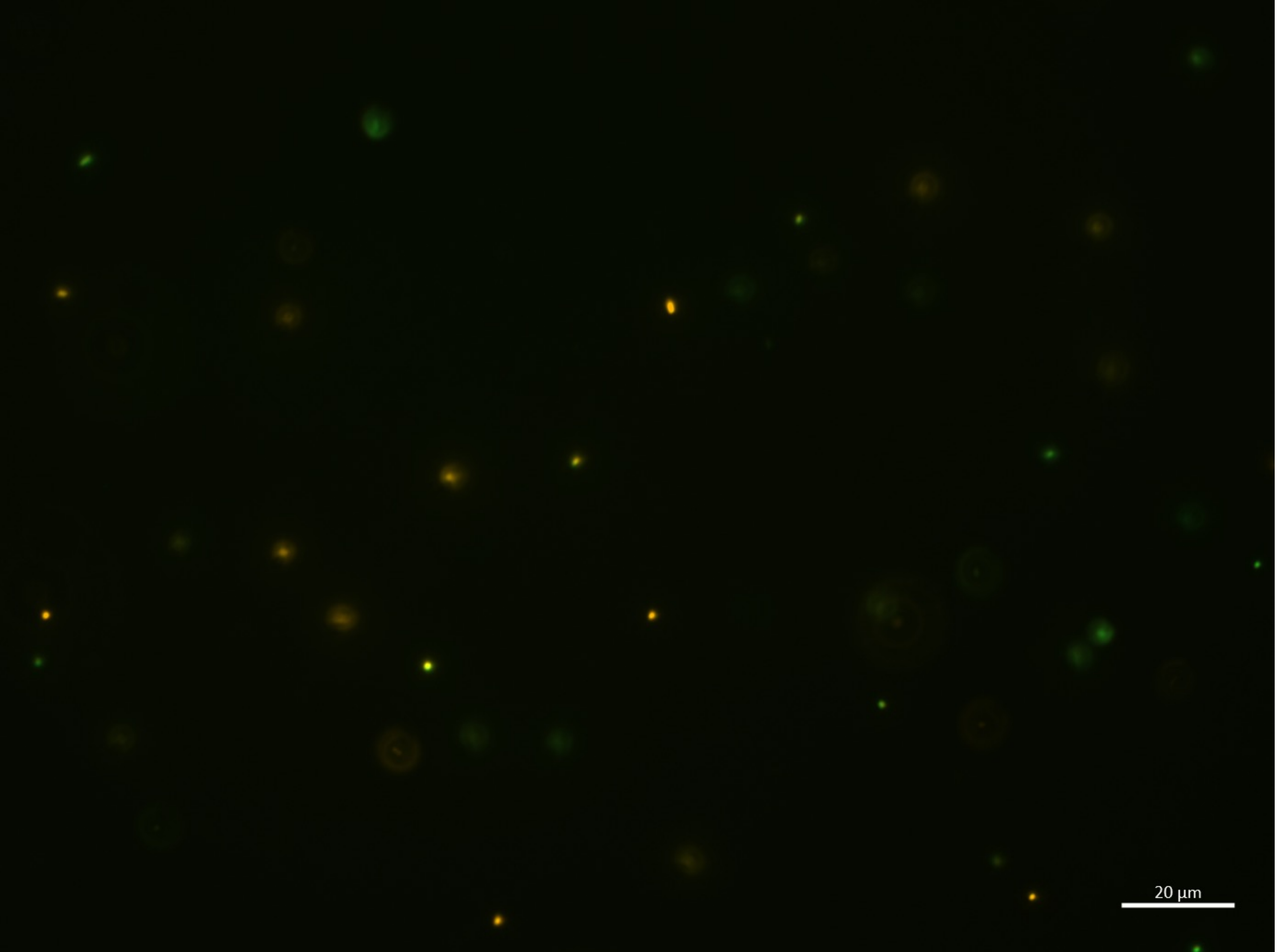

| | |
|---|---|
| B.bifidum | **29** |
| E.coli | 6 |
| **Sum** | 35 |

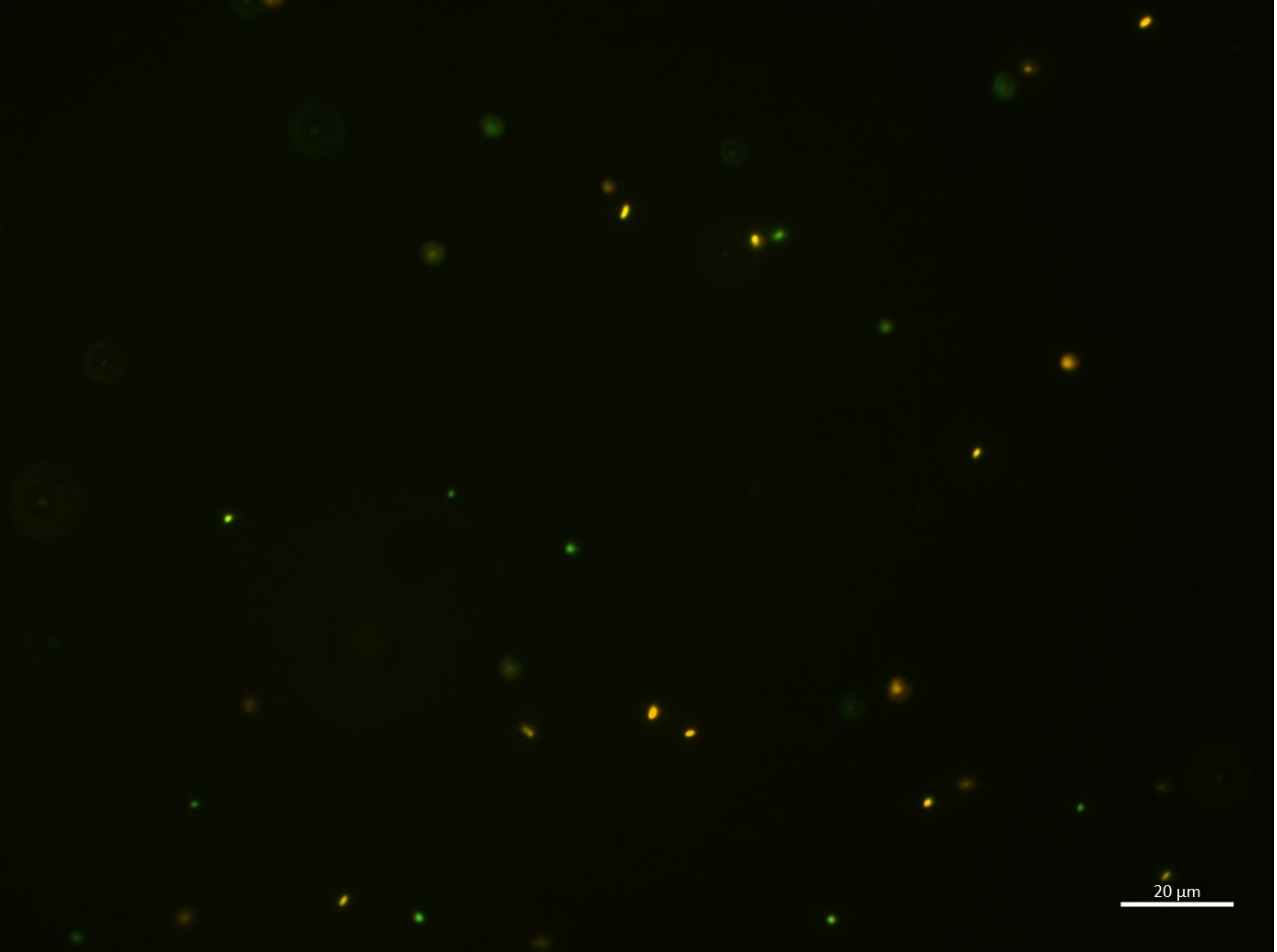

20:00-N

| | |
|---|---|
| B.bifidum | **24** |
| E.coli | 11 |
| **Sum** | 35 |

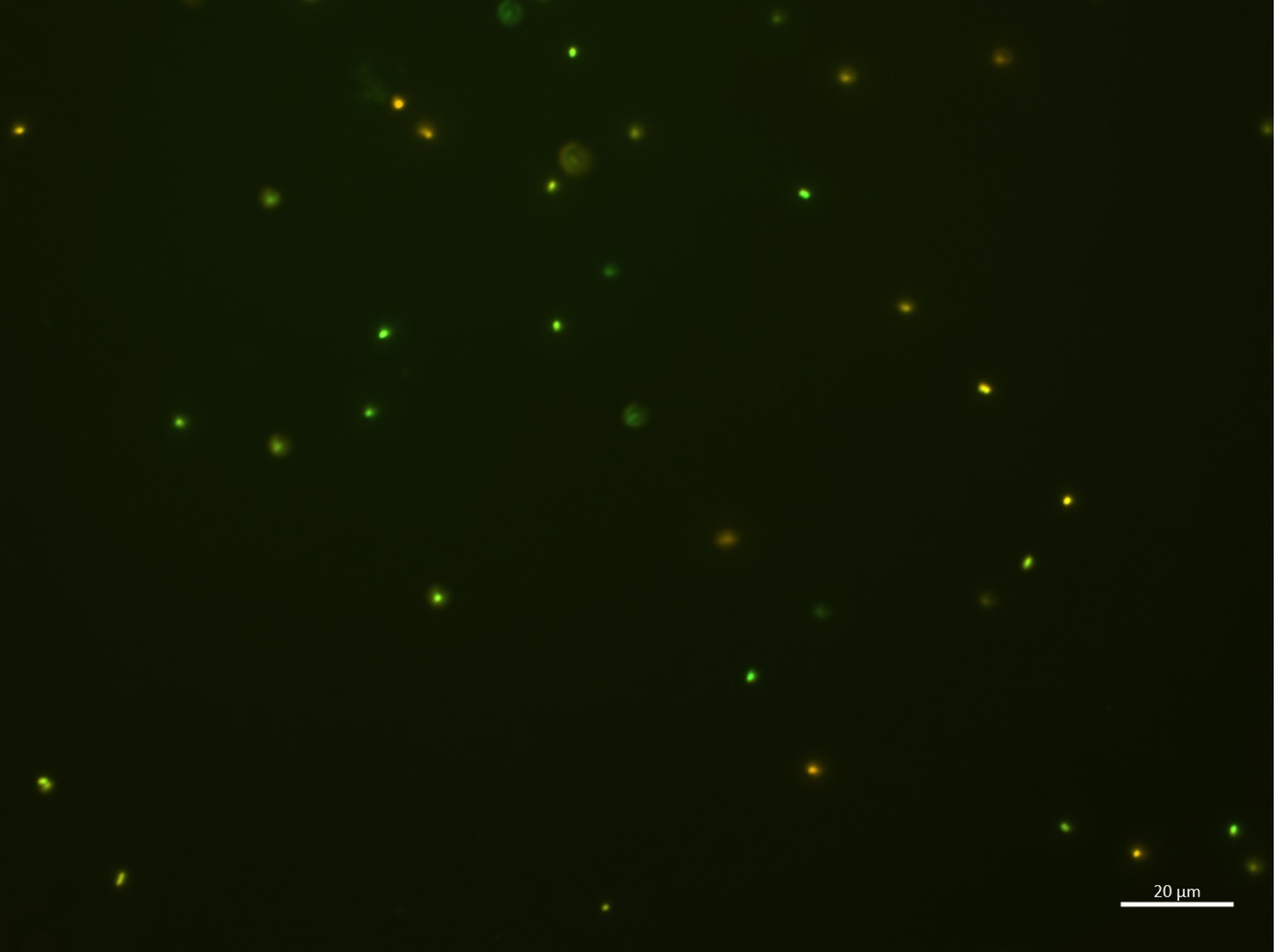

20:00-O

| | |
|---|---|
| B.bifidum | **22** |
| E.coli | 13 |
| **Sum** | 35 |

24:00-A

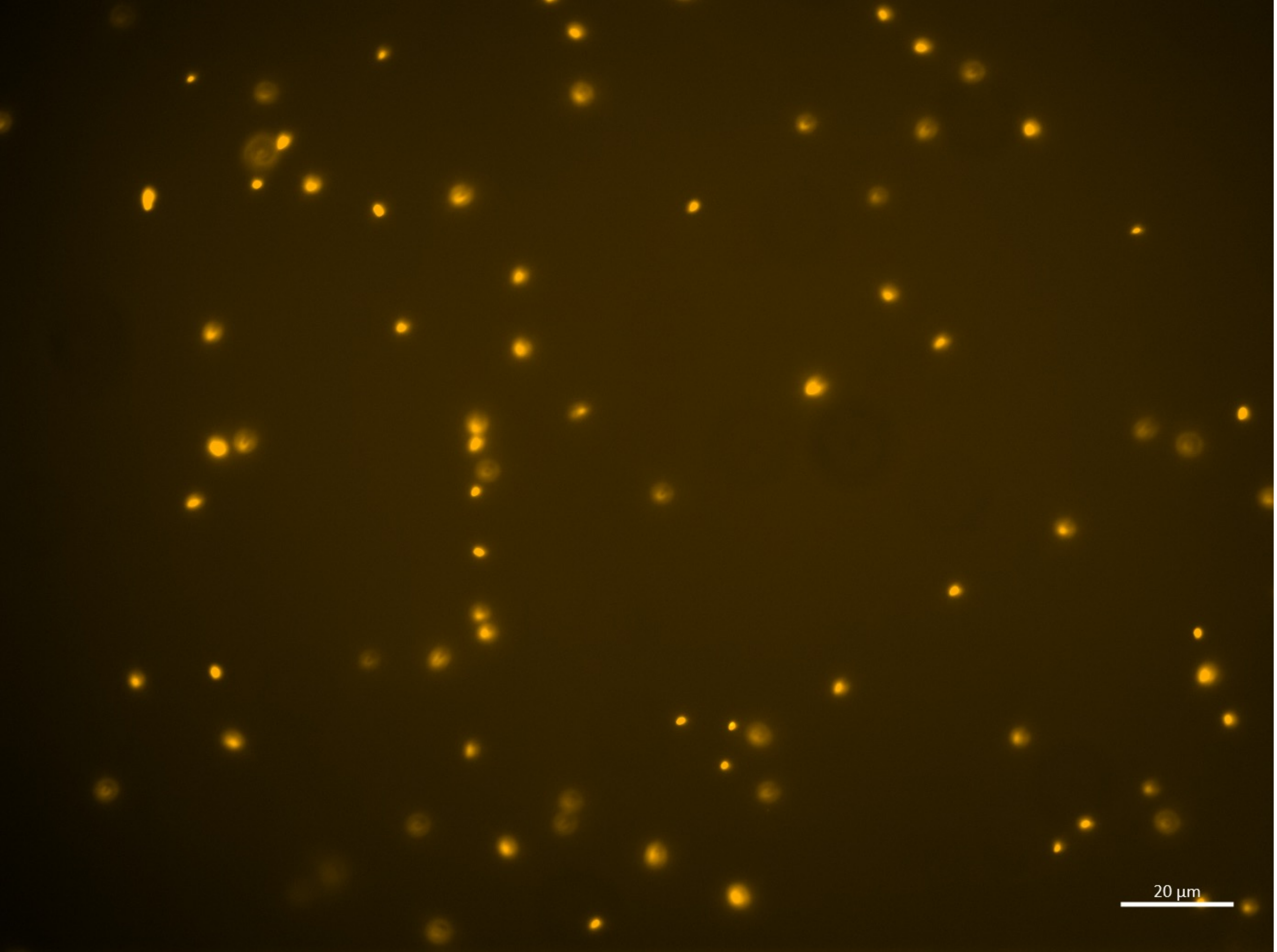

| |
|---|
| B.bifidum |
| E.coli |
| Sum |

| | |
|---|---|
| B.bifidum | 100 % |

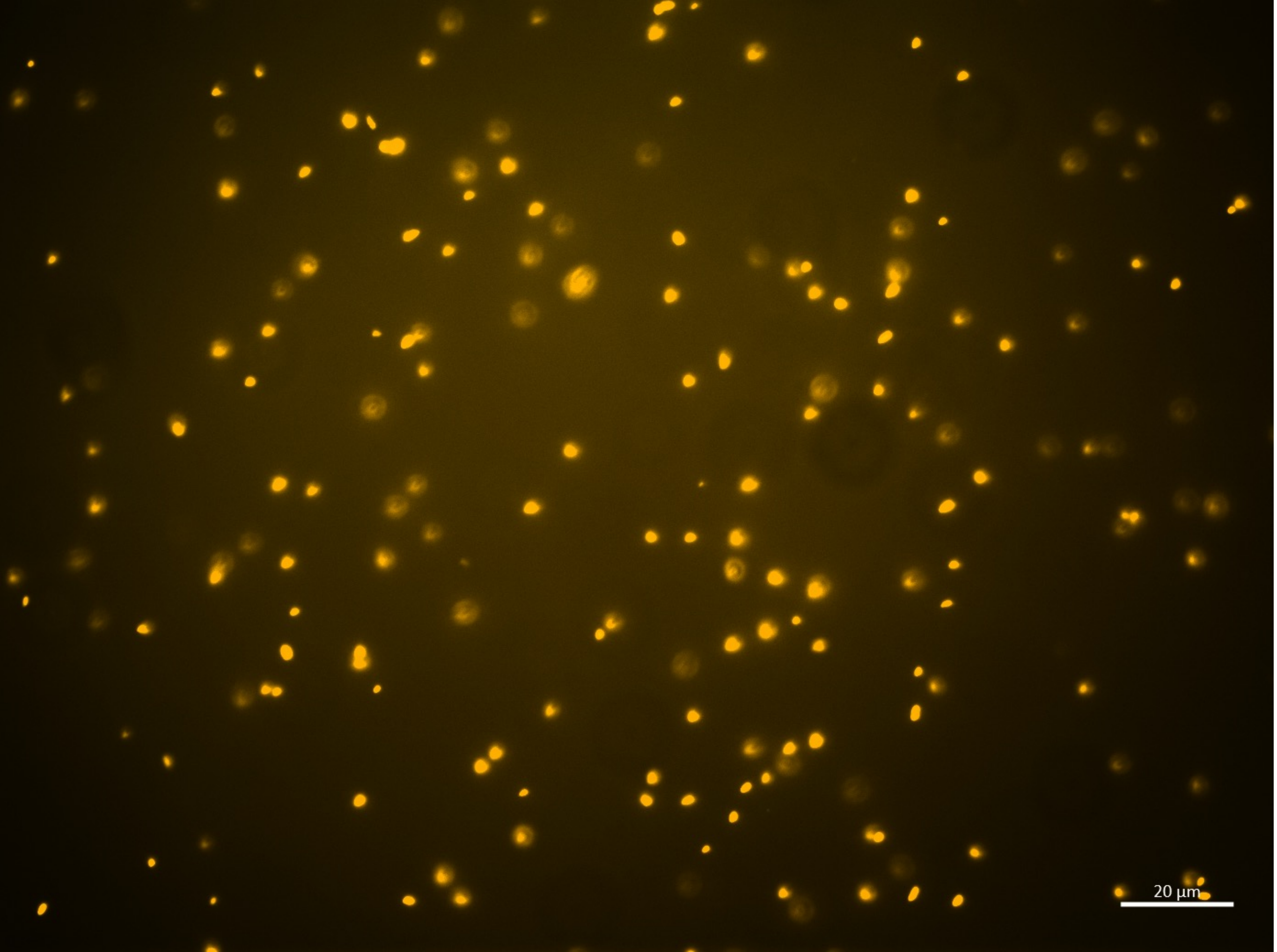

24:00-A

| |
|---|
| B.bifidum |
| E.coli |
| Sum |

| | |
|---|---|
| B.bifidum | 100 % |

24:00-A

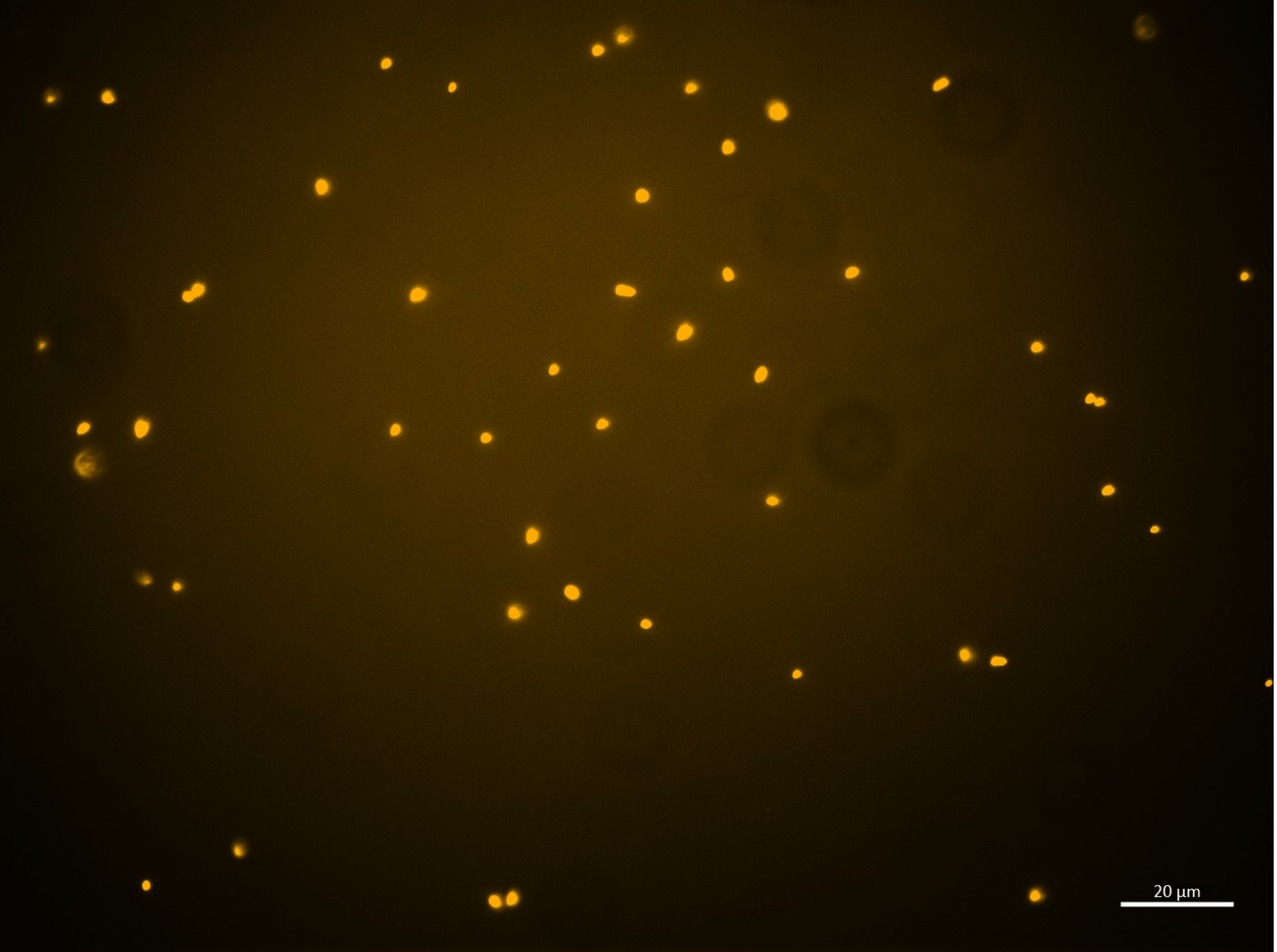

| |
|---|
| B.bifidum |
| E.coli |
| Sum |

| | |
|---|---|
| B.bifidum | 100 % |

24:00-A

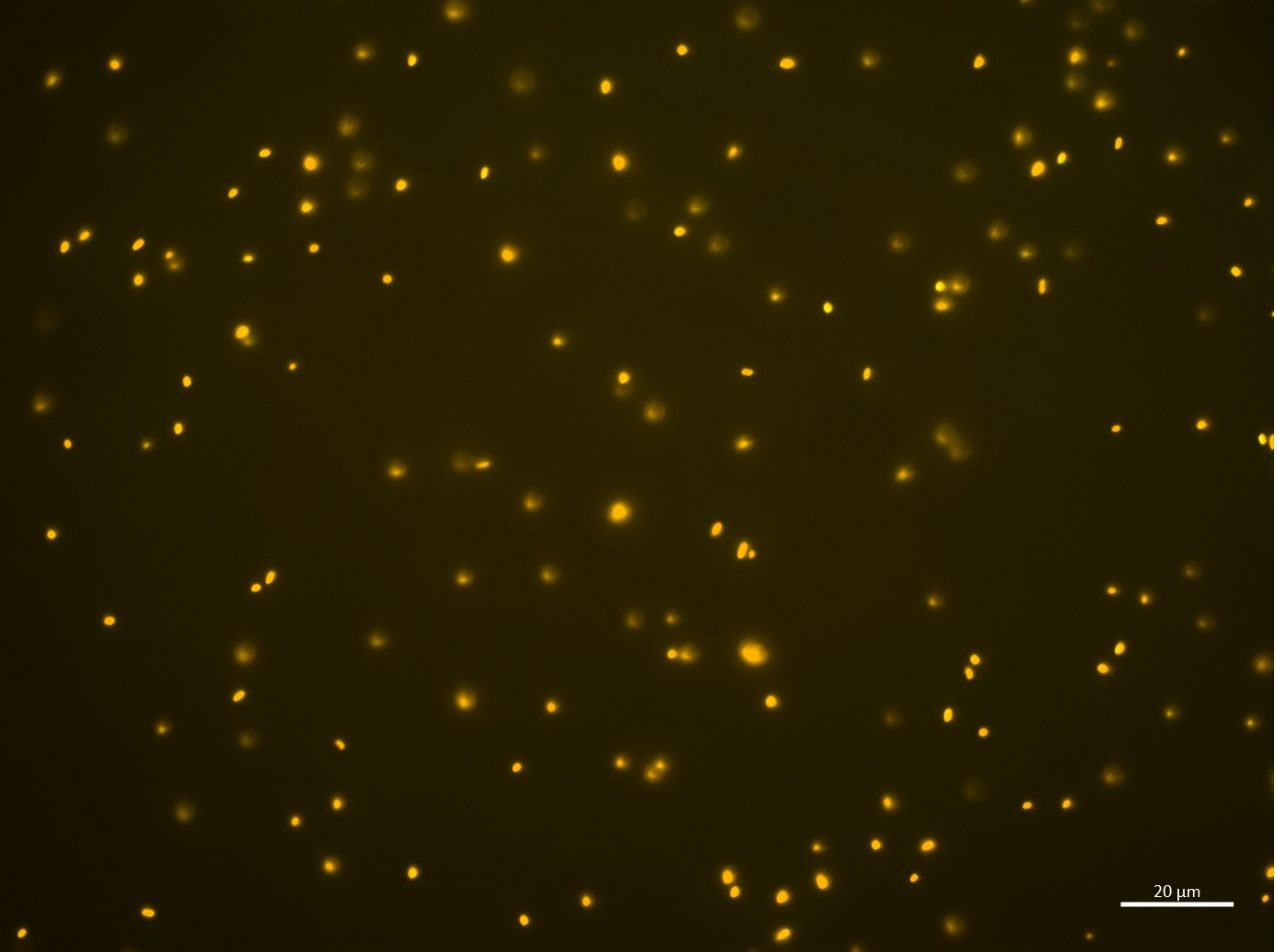

| |
|---|
| B.bifidum |
| E.coli |
| Sum |

| | |
|---|---|
| B.bifidum | 100 % |

24:00-A

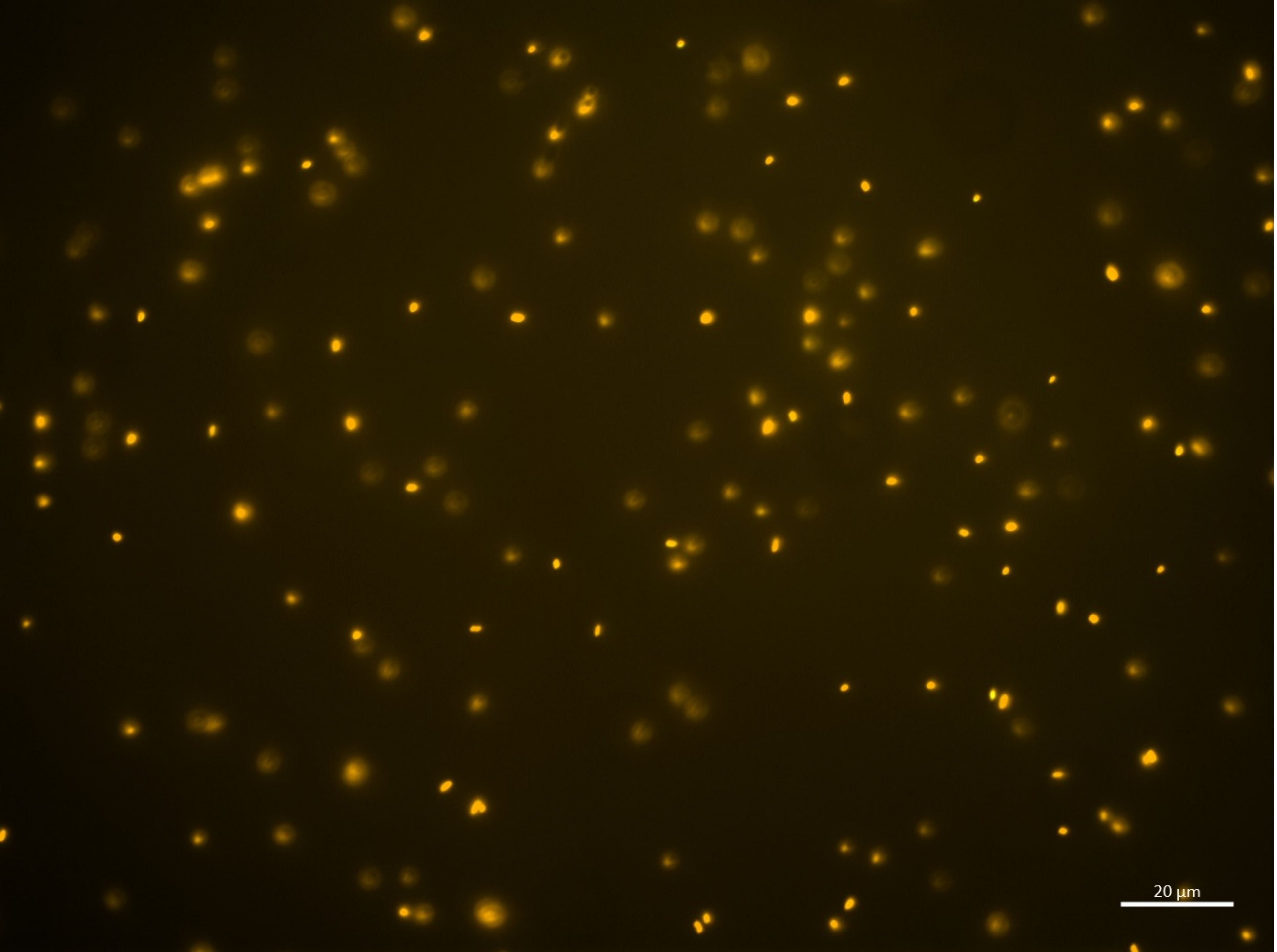

| |
|---|
| B.bifidum |
| E.coli |
| Sum |

| | |
|---|---|
| B.bifidum | 100 % |

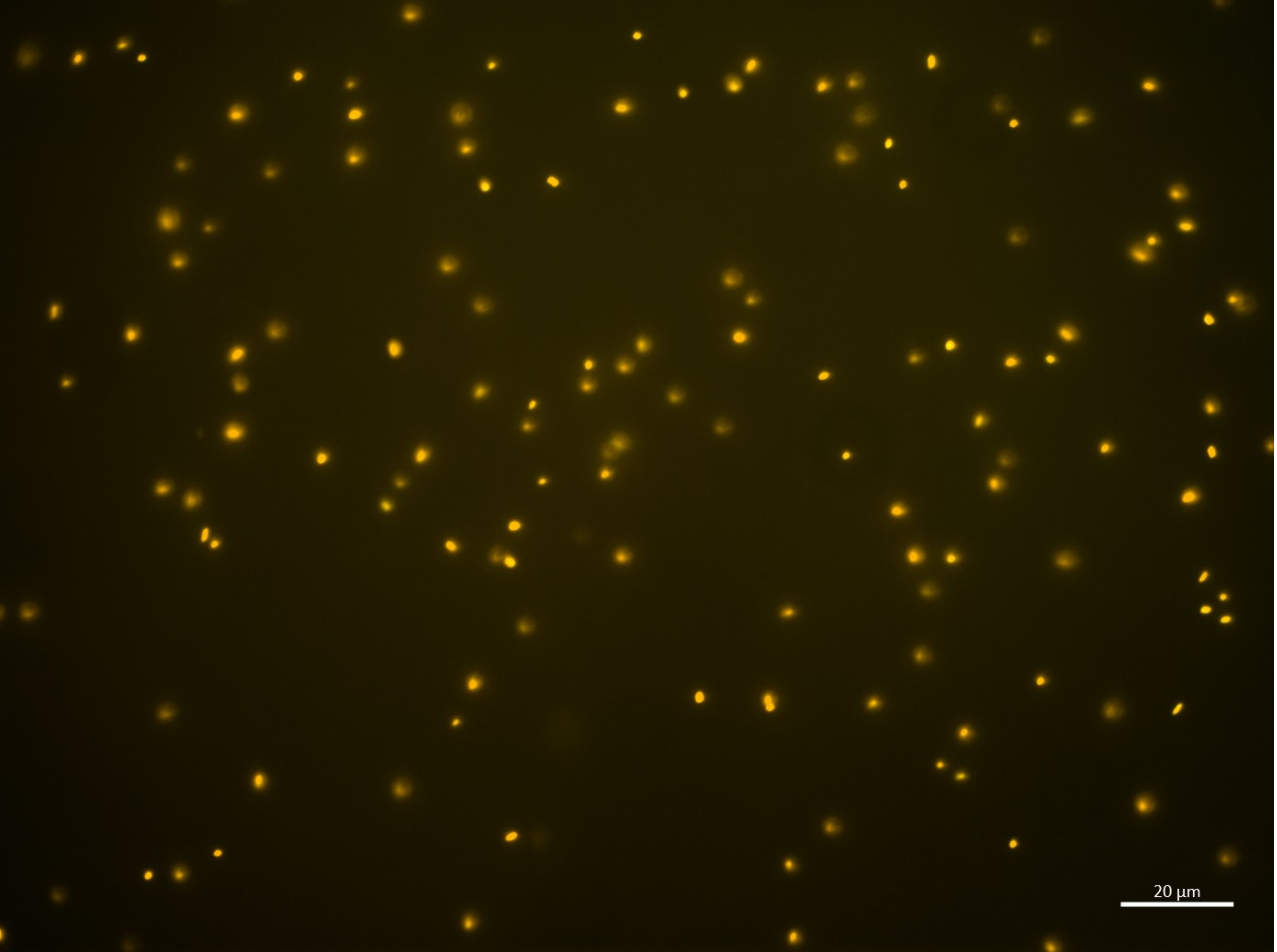

24:00-A

| |
|---|
| B.bifidum |
| E.coli |
| Sum |

| | |
|---|---|
| B.bifidum | 100 % |

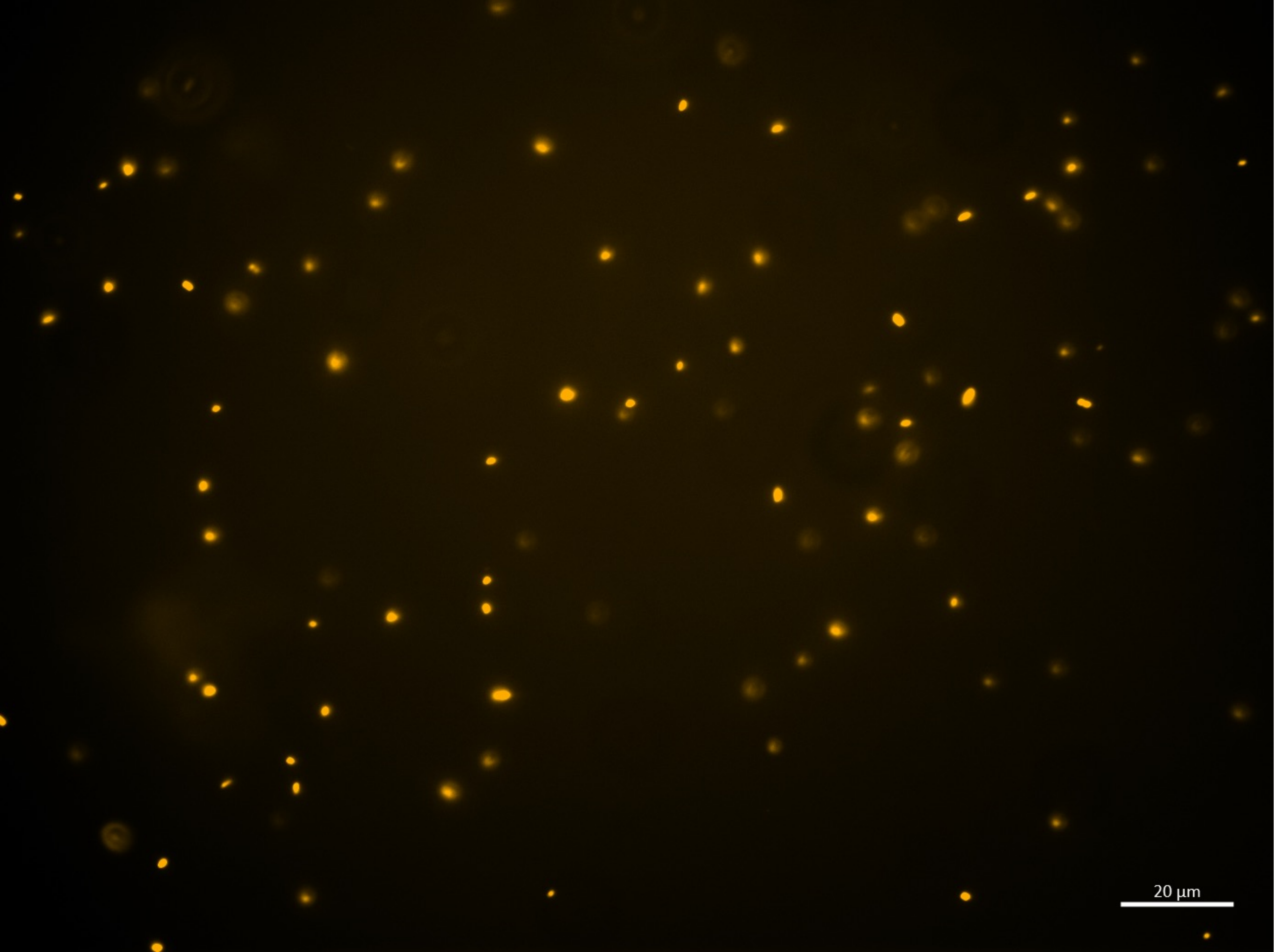

24:00-A

| |
|---|
| B.bifidum |
| E.coli |
| Sum |

| | |
|---|---|
| B.bifidum | 100 % |

# 24:00-A

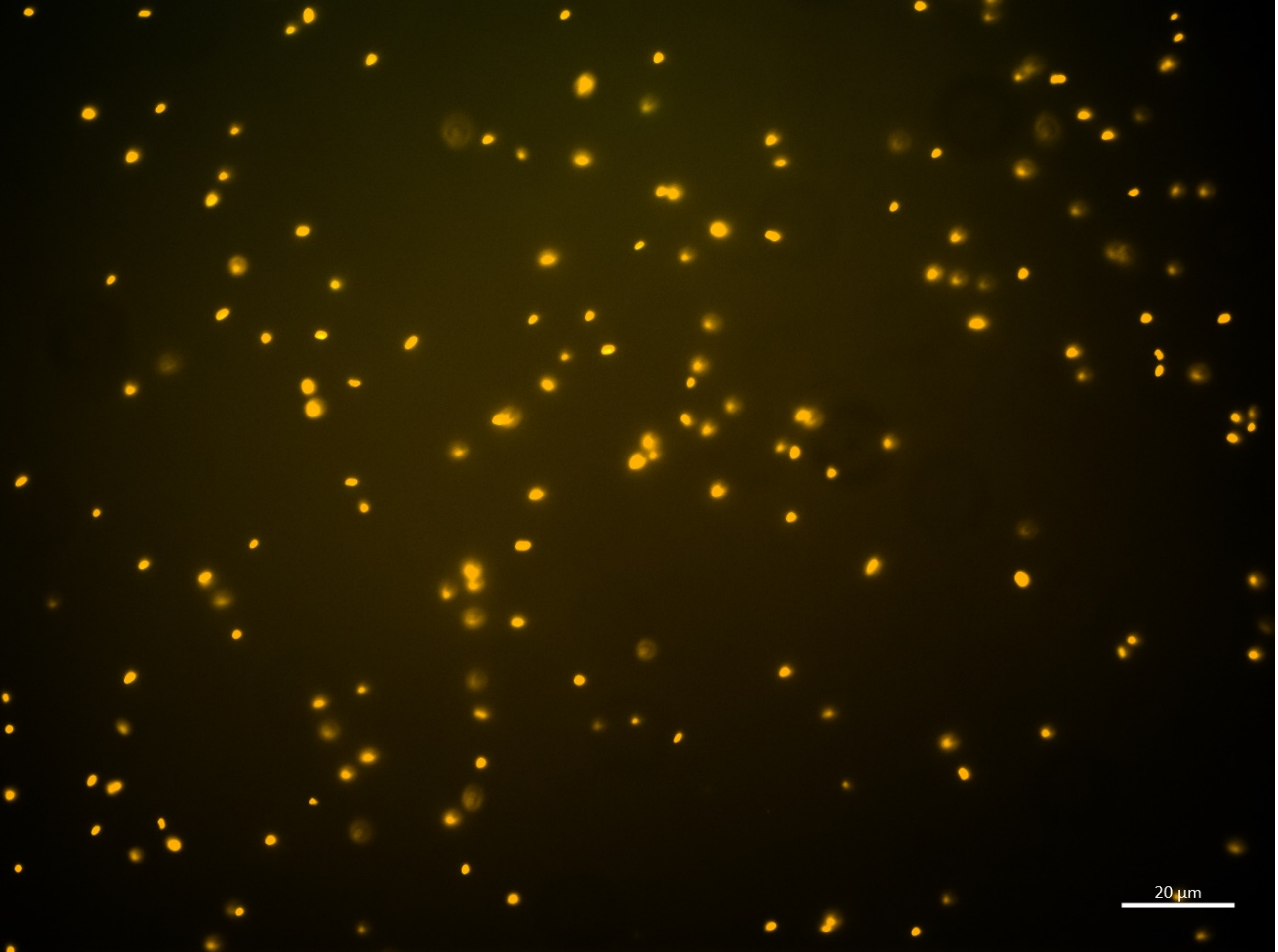

| |
|---|
| B.bifidum |
| E.coli |
| Sum |

| | |
|---|---|
| B.bifidum | 100 % |

24:00-A

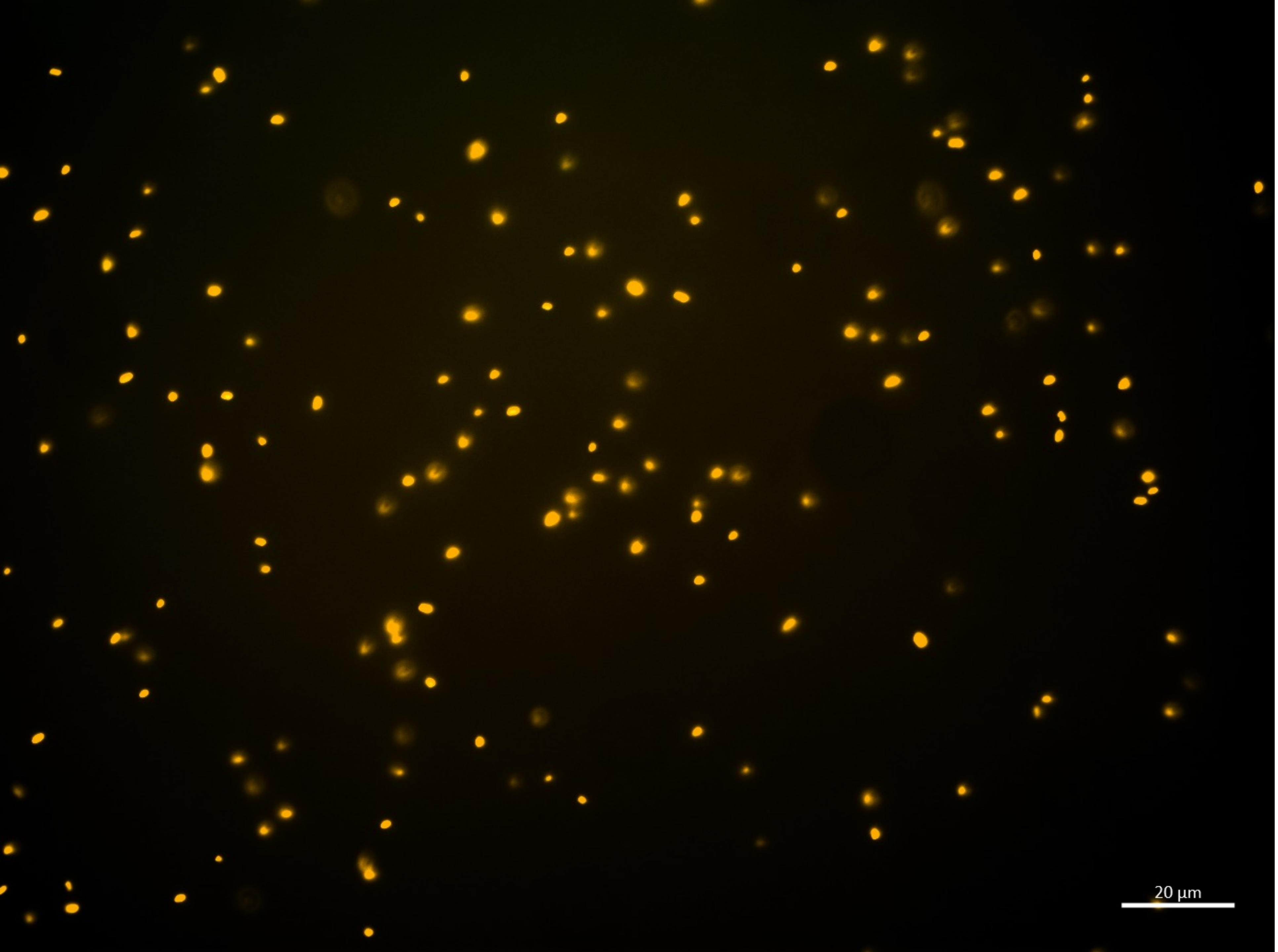

| |
|---|
| B.bifidum |
| E.coli |
| Sum |

| | |
|---|---|
| B.bifidum | 100 % |
